# Long-term survival of asexual *Zymoseptoria tritici* spores in the environment

**DOI:** 10.1101/2024.02.29.582720

**Authors:** William T. Kay, P. O’Neill, Sarah J. Gurr, Helen N. Fones

## Abstract

The fungal phytopathogen *Zymoseptoria tritici*, causal agent of the economically damaging Septoria tritici blotch of wheat, is different from most foliar fungal pathogens in that its germination occurs slowly and apparently randomly after arrival on the leaf surface and is followed by a potentially prolonged period of epiphytic growth and even reproduction, during which no feeding structures are formed by the fungus. Thus, understanding the cues for germination and the mechanisms that underpin survival in low-nutrient environments could provide key new avenues for disease control. In this work, we examine survival, culturability, and virulence of spores following transfer from a high nutrient environment to water. We find that a sub-population of *Z. tritici* spores can survive and remain virulent for at least 7 weeks in water alone, during which time multicellular structures split to single cells. The fungus relies heavily on stored lipids; however, if cell suspensions in water are dried, the cells survive without lipid utilisation. Changes in gene expression in the first hours after suspension in water reflect adaptation to stress, while longer term starvation (7 days) induces changes particularly in primary metabolism and cytochrome P450 (CYP) gene expression. Importantly, we also found that *Z. tritici* spores are equally or better able to survive in soil as in water, and that rain-splash occurring 49 days after soil inoculation can transfer cells to wheat seedlings growing in inoculated soil and cause Septoria leaf blotch disease.

## Introduction

*Zymoseptoria tritici* is an ascomycete fungus that causes the economically damaging wheat disease, Septoria tritici blotch. Despite significant research effort, open questions remain around the strategy used by this fungal pathogen to obtain nutrients [1]. When infecting wheat, it forms no feeding structures and is generally considered a ‘stealth’ pathogen, a biotroph that evades detection by its host partially through slow initial growth [2]. However, some doubt has been cast upon this lifestyle description [1, 3, 4]. [4] demonstrated that the heavily-studied isolate IPO323 can spend over ten days on the leaf surface prior to invasion. This period of surface dwelling is not passive, but can include hyphal extension and exploration of the leaf surface, reproduction by budding [5–7] and even the formation of biofilms [8]. It has been shown that germination, hyphal extension and the subsequent phases of leaf infection, including leaf penetration and the formation of fruiting bodies, are all asynchronous [9]. Following an initial resting phase that lasts up to 15 days, growth on the leaf surface prior to penetration continues for between 2 and 17 days, giving a total of up to 18 days on the leaf surface under optimal conditions [9]. This long, variable period of surface survival and growth stands in sharp contrast to many other fungal plant pathogens, which, with limited energy stores in the spore, have a short time frame for leaf entry and nutrient uptake from the plant and are thus adapted to navigate the leaf surface efficiently [4, 10, 11]. These fungi have highly predictable developmental processes from the detection of a host surface to the formation of feeding structures inside the leaf; while *Z. tritici* does follow a series of predictable steps from germination to entry via stomata to colonisation of the apoplast and pycnidiation in substomatal spaces [12, 13], the timing and the extent of growth at each stage is variable [7, 9]. Thus, rather than a carefully choreographed process of germination and host invasion that is efficient and responds to host cues, *Z. tritici* presents a picture of a fungus whose germination occurs at a random time after arrival on the leaf surface, whose growth is random with respect to entry points, variable in extent and prone to deviating into alternative developmental processes such as blastosporulation or biofilm formation, whether it is on a susceptible or resistant host [4–7, 9].

These unusual epiphytic behaviours in *Z. tritici* provoke questions: primarily, how does *Z. tritici* survive these long periods in the low-nutrient epiphytic environment? Arrival of pycnidiospores - the secondary inoculum of *Z. tritici* produced repeatedly during polycyclic infection of wheat and responsible for disease spread across a field and vertically through a canopy - generally occurs via rain-splash. During this process, the nutrient rich cirrus in which pycnidiospores are extruded from the infected leaf and which protects them from desiccation [13, 14] is diluted and the spores are therefore exposed to a rapid drop in nutrient availability and osmotic pressure. Previously, we showed that in the first few minutes after blastospores are transferred from YPD to water - experiencing a similar drop in osmotic pressure and nutrient availability - both their virulence and culturability fall sharply, unless protected by added osmolytes [15]. This is a counter-intuitive finding, given the similarity of the change in environment between these experiments and rain-splash dispersal. However, rain-splash is an inefficient method of finding a susceptible host, with many spores landing on non-host surfaces or soil. We hypothesised that the short-term drop in culturability and virulence might indicate the initiation of a process that would allow longer term survival of spores in unfavourable environments. In this work, we therefore tested the hypothesis that *Z. tritici* spores would be able to survive for extended periods without nutrients. We find that a sub-population of spores remains culturable and virulent in water for at least 49 days, and we investigate nutrient use and gene expression during this prolonged starvation period.

## Results

### Survival of *Z tritici* blastospores in water

Following rain-splash dispersal, asexual spores of *Z. tritici* are most likely to land on host or non-host plant surfaces, or soil. They will be suspended in rainwater which may, or may not, include some dissolved nutrients from the cirrus [16–19]. This is likely to represent a low-nutrient environment. To determine how well *Z. tritici* spores can survive in such environments, we investigated their survival in autoclaved MilliQ water, representing the most extreme version of these conditions, starvation (Figure 1). Spores were assessed in multiple ways over a 49-day period. Live/dead staining with propidium iodide revealed that the proportion of live spores (defined as a fungal structure having at least one live constituent cell; Fig. 1A,C; Fig. S1) fell slowly for the first five days and then declined very rapidly between days five and seven. However, from day seven onwards, the rate of decline slowed again, with the proportion of live spores remaining steady after around 20 days at approximately 10%. Visual inspection shows that there is an increased number of dead cells per spore and more visible lysed cells at the end of the experimental time course (Fig. S1). Percentage culturability (defined as the ability of a spore to form a colony on YPD agar; Fig. 1B) was lower than the % live cells at all time points, but followed a similar pattern, with around five percent of spores still culturable after 49 days in water. Spore size, measured by the mean number of cells in each spore, followed a hollow curve with rapid initial decline slowing over time and reaching the minimum of one cell per spore by day 49 (Fig. 1D). Strikingly, the total number of spores per ml of suspension increased over the first ∼7 days, reaching a plateau of over twice the starting number, which was maintained until around day 40 (Fig. 1E). This increase suggests either budding is occurring or that larger spores can split. Instances of budding were seen (Fig. 1C; yellow arrows), as were instances of death in non-end-cells (Fig. 1C; white arrows), which might represent points at which spores may later split into two or more. Exemplar images taken after 5 and 42 days in water are given in Fig. S2. Collectively, these results show that ∼10% of blastospores remained viable and culturable after 49 days in water.

**Figure 1.**
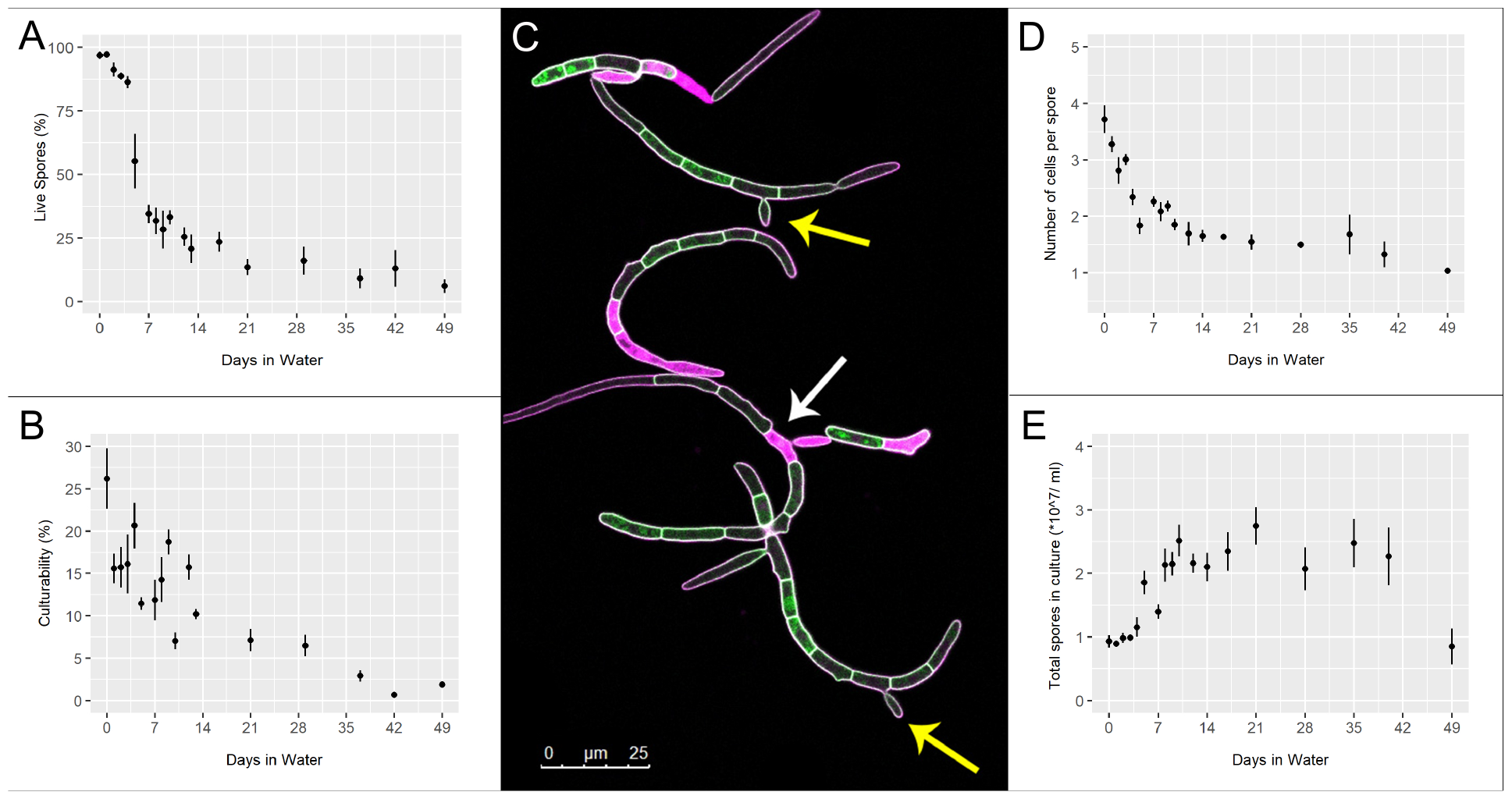
Assessment of *Z. tritici* blastospore populations over 49 days suspended in water. **A:** Percentage viable blastospores (spores with at least one live cell; assessed by live/dead staining with 0.05% (w/v) propidium iodide). **B:** Percentage culturability - number of colonies from plating 100 blastospores as quantified by haemocytometer. **C:** Blastospores expressing cytoplasmic ZtGFP (green) were stained with propidium iodide (pink) after 5 days in water. White arrow highlights an example dead cell; yellow arrows highlight instances of budding growth. **D:** Mean number of cells per blastospore. **E:** Total blastospore count of populations over time using a haemocytometer. In all experiments, data at each time point are means of counts from 4 independent experiments, each containing at least 4 confocal images totalling at least 100 blastospores (A, D); or 3 spread plates (B) or 3 haemocytometer counts (E). Error bars show SE.

### Nutrient stores utilised by *Z tritici* blastospores in water

To gain insight into the possible energy sources used by *Z. tritici* to survive this extended period of starvation, we investigated the use of three common fungal storage compounds, lipids [20], glycogen [21] and trehalose [21, 22]. Lipid droplets were visualised using the fluorescent lipid stain BODIPY^®^ 493/503 and the percentage of spores occupied calculated (Figure 2). Mean lipid content decreased rapidly for the first ∼14 days, and then remained at a low plateau of ∼10% for the rest of the time course (Figure 2A). There is a clear visual difference in the amount of BODIPY^®^ staining in cells at the beginning and end of the time course (Figure 2B, D) and a strong positive correlation (Pearson’s product-moment correlation = 0.94) between the percentage of live spores and the mean spore lipid content (Figure 2C). In addition, we measured the glycogen and trehalose content of spores at days 0, 4 and 8 after suspension in MilliQ. These measurements were carried out by measuring glucose release following enzymatic breakdown of the two compounds; control samples without enzyme treatment therefore measured native free glucose in *Z. tritici* cells. Native free glucose was low (<2 *µ*g/ml per 10^8^ cfu) prior to starvation, and fell to <0.2 *µ*g/ml per 10^8^ cfu after 8 days of starvation (Figure 3), although this decrease is non-significant (ANOVA, P = 0.52). *α*-amyloglucosidase treatment yielded increases in measured glucose up to 64.4 *µ*g/ml per 10^8^ cfu on day 0. The concentration of glucose liberated by *α*-amyloglucosidase appeared to fall over the time course of the experiment (this reduction was non-significant; ANOVA, p = 0.11) but always remained significantly greater than the glucose concentration of controls (*t* -tests, p = 0.0014, 0.005 and 0.002 respectively for days 0. 4 and 8), indicating the presence of glycogen in *Z. tritici* cells grown on YPD agar (Fig 3A). The glucose concentration in samples treated with trehalase showed no significant difference from the controls on any day (*t* -tests, p = 0.45, 0.49 and 0.21, respectively for days 0, 4 and 8), indicating that *Z. tritici* does not produce trehalose as a storage compound during growth on YPD agar (Fig 3B). Taken together, these results indicate that lipids are likely the main energy source for *Z. tritici* spores during starvation.

**Figure 2.**
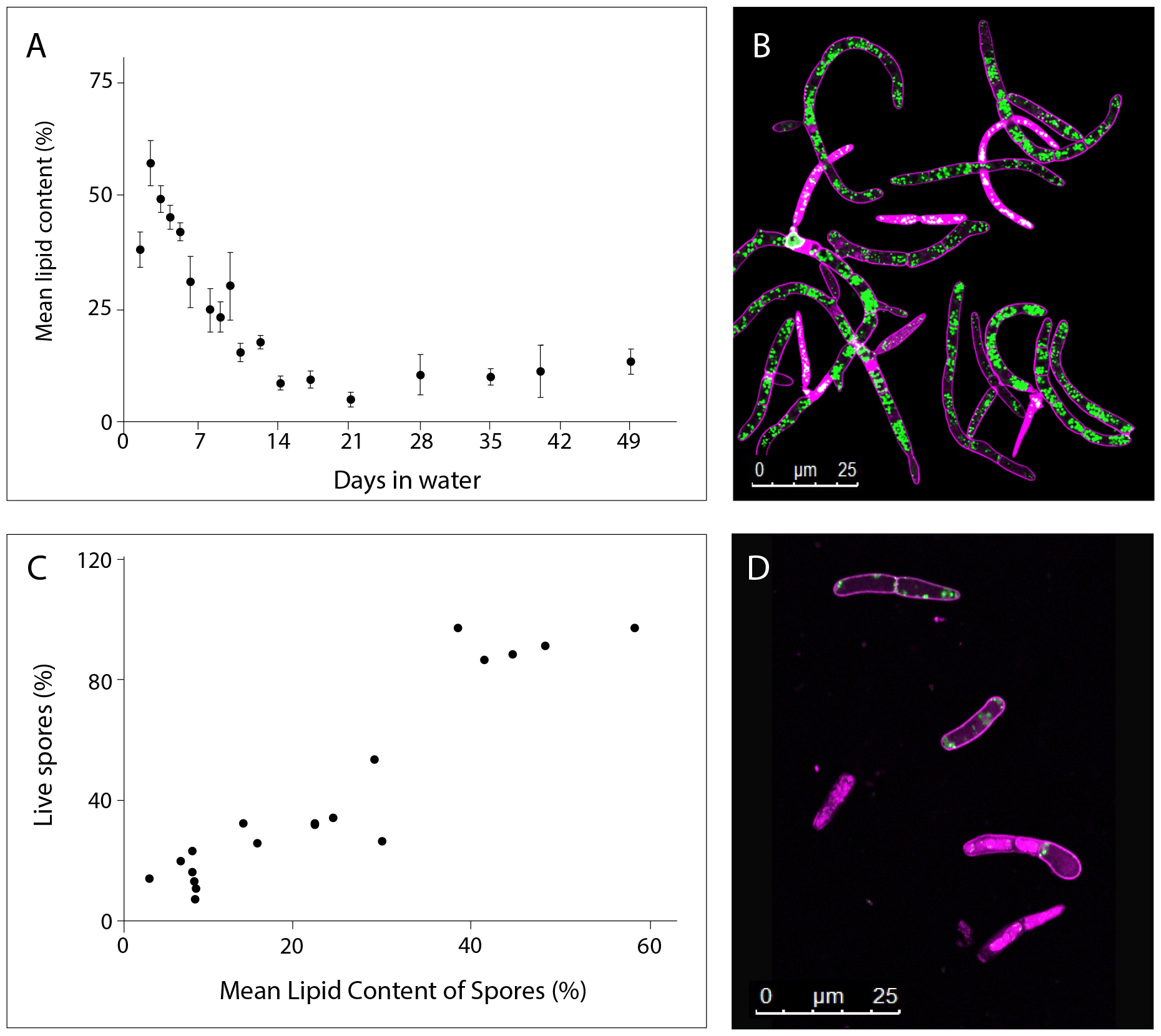
Blastospores show depletion of lipids over time when suspended in water. **A**: average percentage area of spores stained by BODIPY^®^ 493/503. Lipid content of fungal blastospore population is calculated as the percentage of image filled with green fluorescence (lipid granules), divided by area of image representing fungal tissue within the bounds of plasma membranes stained by propidium iodide (PI). Data are means of assessments from 4 independent experiments, each containing at least 4 confocal images of spore populations for each time point. **B**: Example image of PI (pink) and BODIPY^®^ (green) -stained cells after 0 days in suspended in water. **C**: Correlation between lipid content and spore viability. Experimental data for spore viability taken from a 49-day time spore-viability course shown in (Figure 1A). Data show positive correlation of 0.94 (Pearson’s product-moment correlation, t = 5.23, df = 16, p < 0.00005. **D**: Example image of PI and BODIPY^®^-stained cells after 49 days in suspended in water. The higher proportion of dead cells, flooded with PI, can be seen in this image, as well as the large reduction in BODIPY^®^-stained lipid granules.

**Figure 3.**
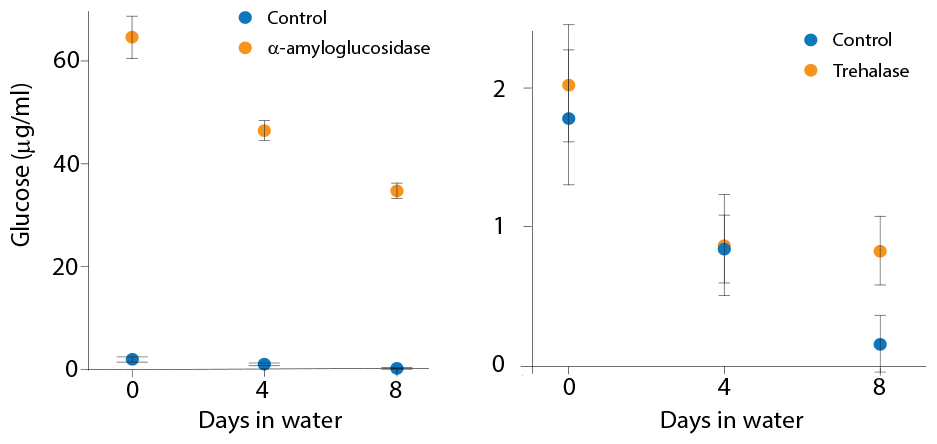
Assessment of glycogen (A) and trehalose (B) concentrations over time in *Z. tritici* cells suspended in water. Blastospores were treated with either *Aspergillus niger α*-amyloglucosidase (A) or porcine trehalase (B). Glucose liberated from each reaction was assayed using a Glucose (GO) Assay Kit. Controls were not treated with either enzyme and so reflect the native free glucose content of *Z. tritici* cells. Sample optical density was measured at 540 nm using a spectrophotometer and compared against prepared glucose standards. Data are means of two experiments, each containing three replicate samples.

### Longevity is maintained with reduced nutrient utilisation during starvation under dry conditions

After *Z. tritici* spores are dispersed from pycnidia by rain-splash, a potential source of abiotic stress is drying. To determine whether drying would reduce the longevity or alter the rate at which lipids are depleted by *Z. tritici* spores during starvation, cells were suspended in sterile MilliQ water as previously but then the suspension was allowed to dry on a sterile plastic surface. At 7-day intervals, dried cells were resuspended in 2 ml sterile distilled water and 100 *µ*l aliquots plated onto YPD agar. Resuspended cells were found to be culturable for at least 56 days (Figure S3). Live-dead staining and lipid content measurements were undertaken for cells resuspended after 28 days, using the same methods as for cells suspended in water (Figure 4). While viability declines over this time period (ANOVA, P < 0.0001), no difference in viability is seen between wet and dry cells at day 28 (Tukey’s simultaneous comparisons, P = 0.49). Significant differences in lipid content were found between samples (ANOVA, P = 0.001), reflecting the same decline in lipids in spores in aqueous suspension as reported above, but there was no significant decline in lipid content for cells in dried suspensions (Tukey’s simultaneous comparisons, P = 0.0006 and P = 0.07, respectively). This suggests that *Z. tritici* spores do not metabolise lipids - or do so at a much reduced rate - when exposed to both starvation and drying, compared to starvation alone.

**Figure 4.**
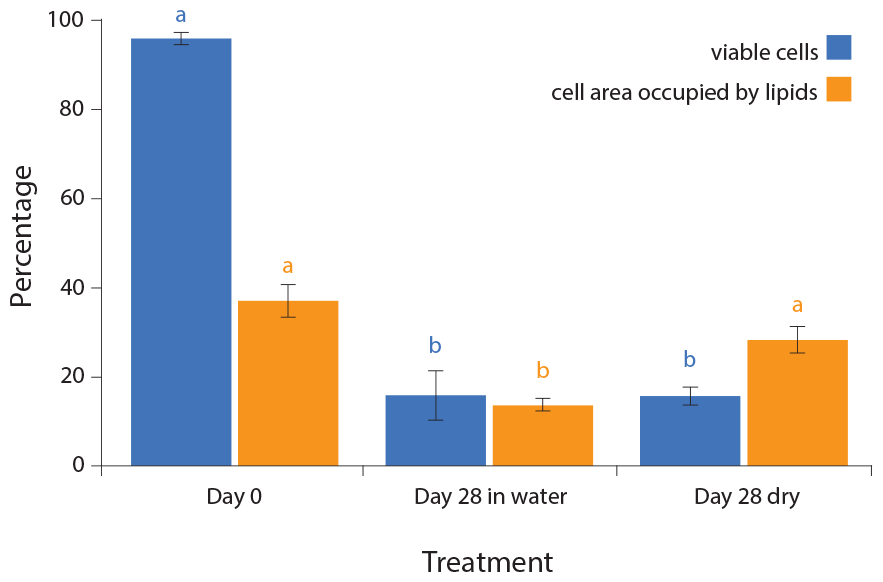
Survival of *Z. tritici* spores suspended in water and allowed to dry out is comparable to spore suspensions maintained in water, but lipid depletion is reduced. Blastospores were suspended in sterile MilliQ water as before. Spore suspensions were either maintained in water or spread onto sterile petri dishes and allowed to dry under sterile air before the dishes were sealed. Percentage viable spores (PI staining) and lipid content (BODIPY^®^ staining) are shown for cell suspensions at day 0 and after 28 days after suspension in water with or without subsequent drying. Values are means of 3 independent experiments and error bars show SE. Significant differences in ANOVAs with Tukey’s simultaneous comparisons are indicated by different letters above bars. Letters apply only to the data whose colour they match.

### Surviving spores retain the ability to cause disease on wheat following starvation

Following the discovery that *Z. tritici* spores can survive at least 49 days of starvation, with or without water, the virulence of these surviving starved spores was assessed. Wheat leaves were inoculated with spores either taken directly from YPD plates or maintained for 49 days in MilliQ water, as before. A range of inoculum densities were used in order to compare virulence more accurately [23] and to account for the 90% drop in spore viability seen after 49 days in water. Inoculation with starved spores led to the production of pycnidia on the wheat leaves, indicating that virulence can be maintained. The number of pycnidia produced by starved inoculum was equivalent to that produced by fresh spore suspensions at 10x lower cfu/ml (Figure 5). When the drop to 10% viability in the starved spore population is taken into account, this equates to an equivalent rate of pycnidium formation per viable spore in the inoculum.

**Figure 5.**
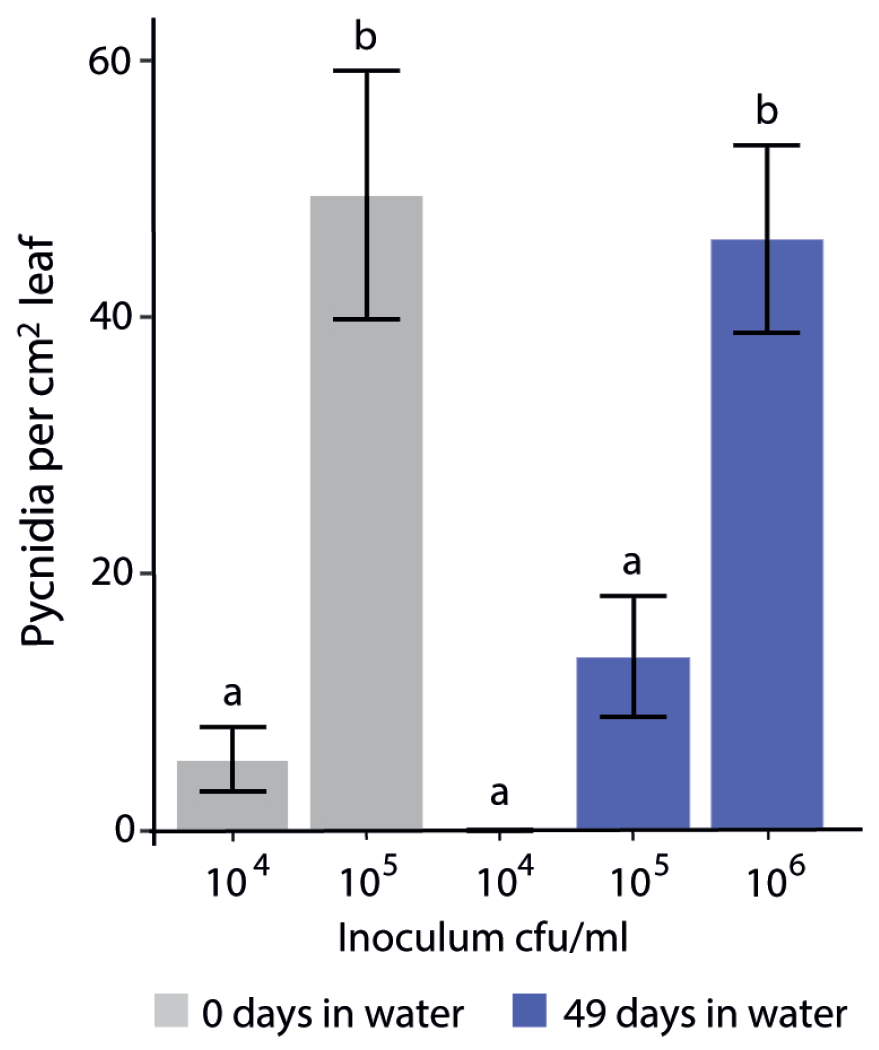
Virulence of *Z. tritici* spores after 49 days in water is comparable to that of fresh spore suspensions. Blastospores were suspended in sterile MilliQ water and maintained for 49 days as before, or suspended from YPD plates immediately prior to use. Spore suspensions were adjusted to 10^4^, 10^5^, or 10^6^ cfu per ml and inoculated onto wheat leaves. Pycnidia were enumerated after 28 days. Values are means of seven independent experiments and error bars show SE. Different letters above bars indicate significant differences in ANOVA (P < 0.0001) with Tukey’s simultaneous comparisons.

### Survival and virulence of blastospores in soil

To increase the field-relevance of these findings, the survival and subsequent virulence of *Z. tritici* blastospores in soil was assessed. Spreading samples of autoclaved, inoculated soil onto YPD plates yielded *Z. tritici* colonies for at least 49 days. To test the virulence of spores ‘stored’ in soil, two experiments were conducted. Firstly, wheat seeds were planted in autoclaved soil which was inoculated with *Z. tritici* blastospores (5 ml of 10^6^ cfu/ml per pot) and subsequently subjected to simulated rainfall. Neither simulated rain-splash from uninoculated soil nor the growing of plants in inoculated soil without rain-splash yielded pycnidia, as expected. However, rain-splash from soil inoculated either on the day of the simulated rainfall or 14 days prior led to pycnidiation (Figure 6A), albeit at a significantly lower rate per cm^2^ than seen with the positive control of brush inoculation with a 10^6^ cfu/ml blastospore suspension (Figure 6A; ANOVA with Tukey’s simultaneous comparisons; P < 0.0001). There was no significant difference in the amount of pycnidia produced by spores that had been in soil for 14 days vs 0 days (Figure 6A; ANOVA with Tukey’s simultaneous comparisons; P = 0.08).

**Figure 6.**
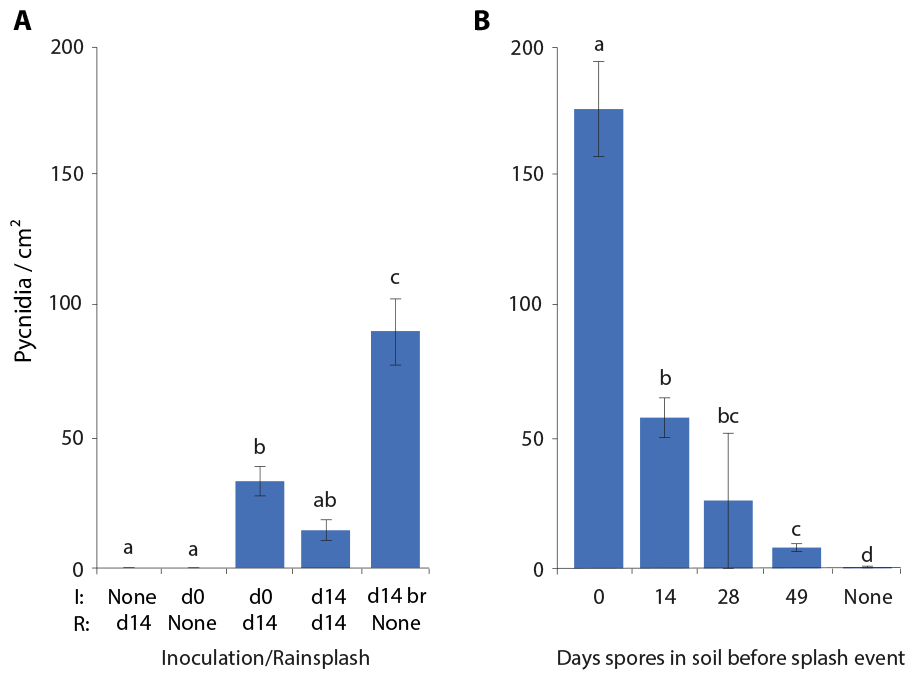
*Z. tritici* spores remain virulent after 49 days in soil, and can infect plants during rain-splash events. **A:** Wheat was grown in autoclaved soil. Either when the first shoots emerged (d0) or when the plants were 14 days old (d14), the soil was inoculated with *Z. tritici* blastospores (5 ml of 10^6^ cfu/ml per cell of a 24-cell tray). Rainfall was simulated on d14 by watering from a height of 2 metres at a rate of 4 litres of sterile distilled water per 24-cell tray from a rose head watering can. Negative controls without *Z. tritici* or without rainfall were included, and brush inoculation (br) on d14 was used as a positive disease control. **B:** Wheat was subjected to simulated rainfall at the indicated intervals after the soil had been inoculated with *Z. tritici* blastospores (5 ml of 10^7^ cfu/ml per cell of a 24-cell tray). Uninoculated soil (‘None’) was used as a negative control. In both **A** and **B**, pycnidia per cm^2^ of leaf was calculated for the cotyledon, first and second leaf 28 days after the rain-splash or brush inoculation. Values are means of three independent experiments and error bars show SE. Different letters above bars indicate significant differences in ANOVA with Tukey’s simultaneous comparisons.

Secondly, soil was inoculated with *Z. tritici* blastospores (5 ml of 10^7^ cfu/ml per pot) and seeds were sown periodically such that plants were 14 days old at various times after the inoculation of the soil. Rainfall was simulated at these time points and plants assessed for the development of pycnidia (Figure 6B). In line with the decline of spore viability seen in water, these rain-splash events yielded progressively fewer pycnidia as the spores aged (ANOVA: P < 0.0001; Figure 6B), but, again in line with previous results, pycnidia were formed even when the simulated rain occurred 49 days after the soil was inoculated. Thus, as in water, spores survive and remain virulent in soil for at least 49 days.

### Changes in gene expression during suspension in water

We hypothesised that the ability of *Z. tritici* spores to survive extended periods of starvation while suspended in water must be underpinned by significant changes in gene expression. To test this and to gain insight into the nature of these changes, we carried out RNAseq to compare gene expression in *Z. tritici* blastospores on YPD to blastospores grown on YPD and then suspended in MilliQ water. Given that the virulence of *Z. tritici* blastospores has been shown to decline rapidly in water [15], we further hypothesised that changes in gene expression would be rapid. We therefore included samples of cells harvested at 1, 4 and 24 hours post suspension in water in this RNAseq experiment. To reveal gene expression associated with longer term starvation, we also included samples of cells harvested after 7 days in water. It proved challenging to extract RNA from later time points due to a large drop in RNA content of cell suspensions, suggesting that global gene expression might be much lower in starved cells, as well as reflecting the high proportion of cells already shown to be dead by this time (only ∼35% of spores contained a live cell and only ∼12% were culturable after 7 days in water - see Fig. 1A&B).

We compared the expression of genes at each of these starvation time-points to their expression under nutrient-replete conditions (YPD agar). There are 108 genes up-regulated and 8 down-regulated in common at all time-points; conversely, there are genes uniquely up-regulated or down-regulated at each time point (Figure 7 & 8). The largest number of uniquely up-regulated genes (437) is seen after 7 days in water, and the least (52) at the end of the first hour. Similarly, the largest number of down-regulated genes (272) is seen after 7 days in water. The lowest number of uniquely down-regulated genes is seen following 4 h of starvation in water. A heatmap of differentially expressed genes shows that the changes in gene expression after seven days are distinct from those at the earlier time points (Figure 9).

**Figure 7.**
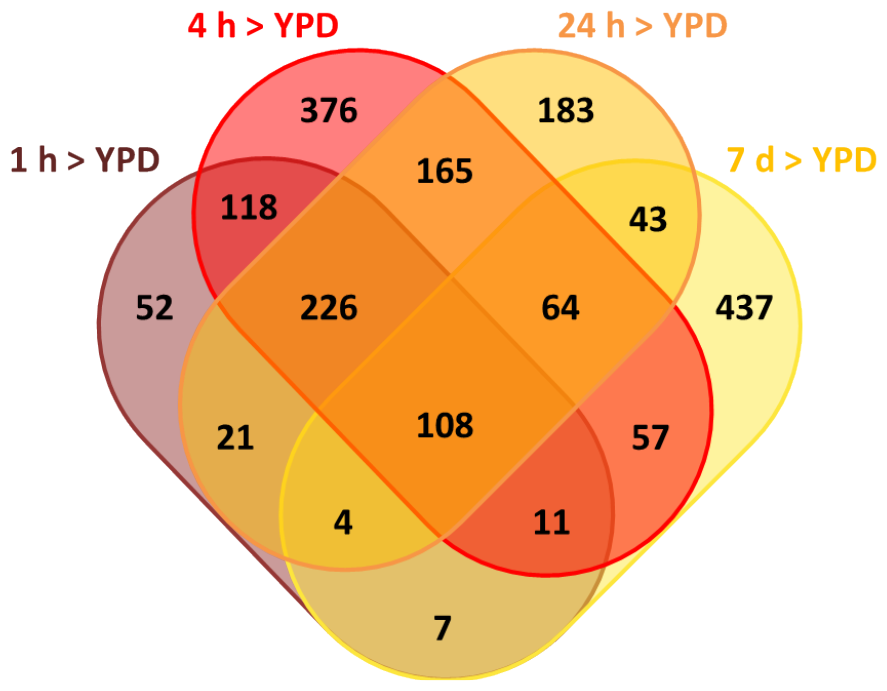
Venn diagram showing numbers of genes up-regulated uniquely or in common with specific other time-points. Venn diagram produced using http://bioinformatics.psb.ugent.be/webtools/Venn/

**Figure 8.**
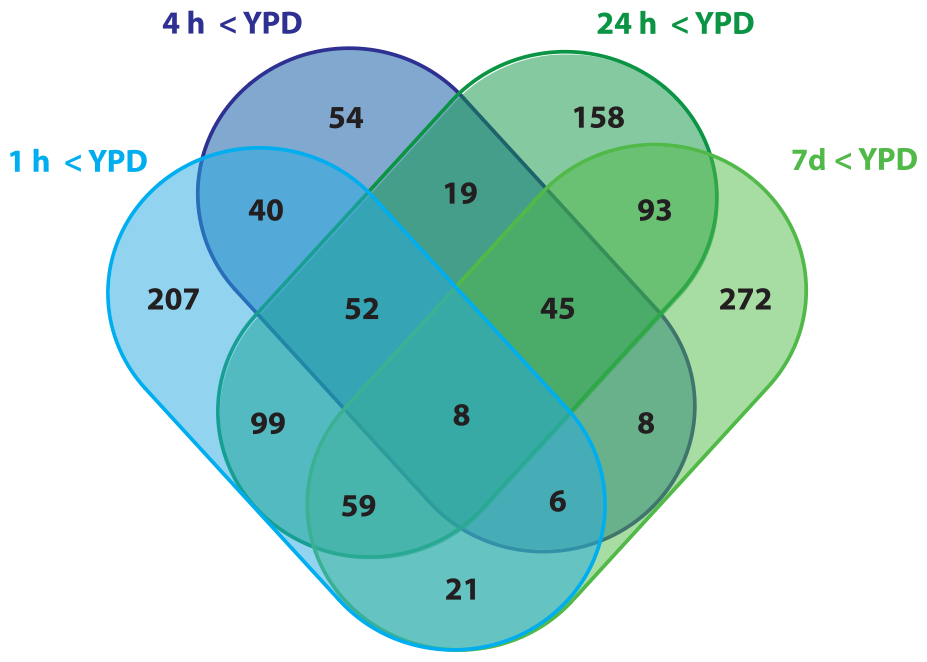
Venn diagram showing numbers of genes down-regulated uniquely or in common with specific other time-points. Venn diagram produced using http://bioinformatics.psb.ugent.be/webtools/Venn/

**Figure 9.**
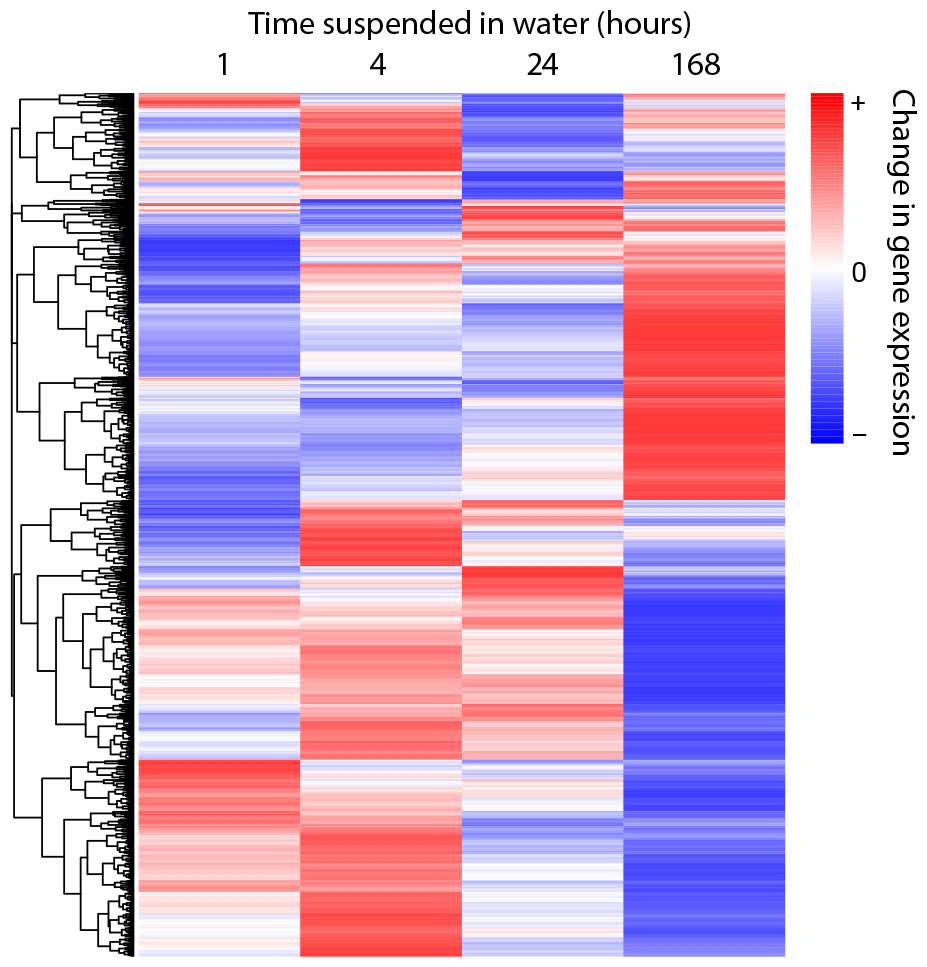
Heatmap of differentially expressed genes at 1, 4, 24 h or 7 days in water, compared to expression on YPD agar. Produced using SR plot [24]

### Genes up-regulated uniquely at 1 h (vs YPD)

Genes uniquely up-regulated at this time are found only on core chromosomes. The ten most up-regulated genes include five which encode proteins with homology to known proteins from other fungi: ZTRI_8.214, encoding a *β*-glucanase; ZTRI_5.41, encoding an N-glycosylated membrane protein; ZTRI_11.419, encoding a cell wall galactomannoprotein; ZTRI_8.260, encoding a methyltransferase, and ZTRI_1.1034, encoding a glycosyl hydrolase (Table S1). The set of 52 uniquely up-regulated genes is significantly enriched (p = 0.016) for GO term GO:0005618, indicating cell wall localisation. There is also enrichment for GO terms denoting molecular functions DNA photolyase activity (p = 0.0032) and for those relating to biological processes involved with circadian rhythm and cell wall organisation (p = 0.0032 and 0.0096) (Table 1). In addition, metabolic pathways relating to peptidoglycan synthesis and cross-linkage are significantly over-represented among the up-regulated genes (Table S2). Other pathways showing enrichment among this gene set compared to the genome as a whole are epoxysqualene and lineolate biosynthesis (Table S2). Searching the list of genes uniquely up-regulated at 1 h for the string ‘transcr*’ in FungiDB [25] returned two genes, ZTRI_5.380 and ZTRI_6.433. Of these, ZTRI_5.380 has orthology to C6 zinc finger domain proteins from other fungi and predicted molecular functions of zinc ion binding and RNA polymerase II-specific DNA-binding transcription factor activity. ZTRI_6.443 is orthologous to genes in *Neurospora crassa* and *Fusarium* sp. encoding the clock protein FRQ, which is not itself a transcription factor but interacts with the transcription factors white collar 1&2 in regulation of the circadian rhythm.

**Table 1:** GO term enrichment among genes uniquely up-regulated after 1 h in water. GO enrichment carried out using tools in fungiDB [25]. Threshold for inclusion in table P < 0.02

| GO ID | GO Term | Genes in background with this term | Genes uniquely up-regulated at 1 h | Fold enrichment | Odds ratio | P-value | Gene IDs |
| --- | --- | --- | --- | --- | --- | --- | --- |
| GO:0030312 | external encapsulating structure | 5 | 1 | 62.24 | 82.33 | 0.0160 | ZTRI_1.1034 |
| GO:0005618 | cell wall | 5 | 1 | 62.24 | 82.33 | 0.0160 | ZTRI_1.1034 |
| GO:0004455 | ketol-acid reductoisomerase activity | 1 | 1 | 311.18 | Infinity | 0.0032 | ZTRI_1.395 |
| GO:0004506 | squalene monooxygenase activity | 1 | 1 | 311.18 | Infinity | 0.0032 | ZTRI_3.484 |
| GO:0003913 | DNA photolyase activity | 1 | 1 | 311.18 | Infinity | 0.0032 | ZTRI_4.76 |
| GO:0007623 | circadian rhythm | 1 | 1 | 311.18 | Infinity | 0.0032 | ZTRI_6.443 |
| GO:0048511 | rhythmic process | 1 | 1 | 311.18 | Infinity | 0.0032 | ZTRI_6.443 |
| GO:0045229 | external encapsulating structure organization | 3 | 1 | 103.73 | 164.72 | 0.0096 | ZTRI_1.1034 |
| GO:0071555 | cell wall organization | 3 | 1 | 103.73 | 164.72 | 0.0096 | ZTRI_1.1034 |
| GO:0071554 | cell wall organization or biogenesis | 4 | 1 | 77.79 | 109.79 | 0.0128 | ZTRI_1.1034 |

### All genes up-regulated at 1 h (vs YPD)

There are 547 genes up-regulated after 1 h in water, the majority of which (533) are on the core chromosomes. The top ten most up-regulated genes (Table S3) include a gene likely to encode a proteolytic enzyme (ZTRI_11.422), a HSP-like protein (ZTRI_6.402) with homology to *Aspergillus* Hsp30/Hsp42, which is implicated in the response to a number of stresses including heat, antifungals and oxidative stress [26–28], two genes with homology to alcohol dehydrogenases (ZTRI_1.494, ZTRI_1.498), two genes encoding major facilitator superfamily transporters (ZTRI_-8.748, ZTRI_11.277) of which one has homology to mannitol dehydrogenase in several fungi and to *Candida albicans* gene C6_02480W_A, known to be part of the core stress response in that organism [29], and two genes predicted to encode trichodiene synthases (ZTRI_ 5.833, ZTRI_1.1590). The other genes encode predicted or hypothetical proteins. ZTRI_ 11.277 encodes a protein likely able to transport sugar and/or phosphate based on PFAM domains and homology; its homologue in *C. albicans, C1_11480W_A* being a stress-induced phosphate transporter under the control of *Hog1*. ZTRI_8.748 has homology to tetracycline efflux transporter genes in other fungi.

Genes up-regulated at 1 h were also enriched in a number of GO terms including molecular functions oxidoreductase activity (GO:0016491; P = 0.0013), unfolded protein binding, heat shock protein binding and chaperone binding (GO:0051082, 0031072, 0051087; P = 0.0017, 0.012, 0.019), as well as tetrapyrrole/haem binding (GO:0046906/0020037; P = 0.018) and lipase activity (GO:0004806, P = 0.0190) (Table 2). Biological processes for which the genes up-regulated at 1 h showed significant enrichment were protein folding (GO:0006457; P <0.0001), oxidation-reduction (GO:0055114; P = 0.0028), fatty acid metabolic process (GO:0006631; P = 0.0019) and a number of related GO terms (oligo-/di-saccharide/trehalose biosynthesis; P = 0.0061 or metabolic process; P = 0.0019) attached to the same two genes, ZTRI_6.64 and ZTRI_7.164 - trehalose phosphatase and trehalose phosphatase synthase. In addition, metabolic pathways relating to fatty acid *β*-oxidation, cis- and trans-hepta- and octadecenoyl-CoA degradation; gingerol, lineolate, sulphur amino acid, methionine, ribonucleotide and trehalose biosynthesis and fatty acid salvage are significantly over-represented among the up-regulated genes (Table S4).

**Table 2:**
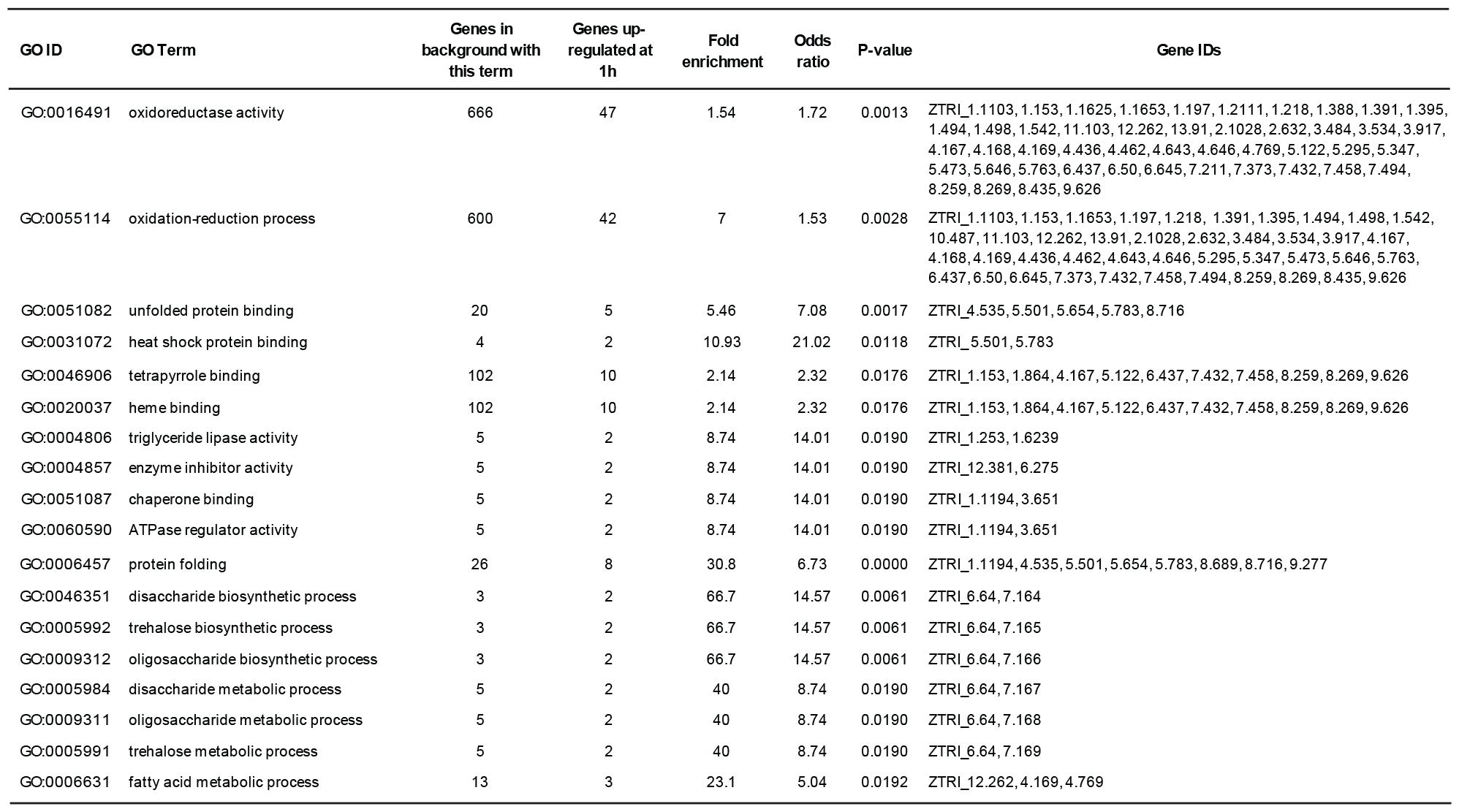
GO term enrichment among all genes up-regulated after 1 h in water. GO enrichment carried out using tools in fungiDB [25]. Threshold for inclusion in table P < 0.02

### Genes up-regulated uniquely at 7 days

There are 732 genes up-regulated after 7 days in water compared to YPD. Of these, 437 are not up-regulated at any other time point. These are mostly on the core chromosomes, but 5 of the 8 accessory chromosomes are also represented. The ten most up-regulated genes unique to the 7-day time point (Table S5) are mostly hypothetical or predicted proteins. GO terms enriched among the 437 genes up-regulated uniquely at 7 d include those representing molecular functions or biological processes relating to oxidation/reduction (GO: 0016705, 0016491, 0055114; P < 0.0005) and binding to cations - in particular, transition metals including specifically iron ion binding, as well as haem binding (GO: 0005506, 0020037, 0046914; P < 0.0005, < 0.0005, 0.001). GO terms relating to transmembrane transport and membrane localisation are also over-represented (e.g. GO:0055085, 0016021; P <0.0005) (Table 3).

**Table 3:** GO terms significantly enriched in genes up-regulated uniquely after 7 days in water. Enrichment analysis carried out using tools in FungiDB [14] P-value threshold for inclusion is 0.02.

| GO ID | GO Term | Genes in background with this term | Genes uniquely up-regulated at 7d | Fold enrichment | Odds ratio | P-value | Gene IDs |
| --- | --- | --- | --- | --- | --- | --- | --- |
| GO:0016705 | oxidoreductase activity, acting on paired donors, with incorporation or reduction of molecular oxygen | 126 | 20 | 4.31 | 5.3800 | 0.0000 | ZTRI_1.1603, 1.1735, 1.1783, 1.1784, 1.187, 1.1950, 1.834, 10.145, 10.489, 11.126, 12.61, 2.10, 2.1087, 3.103, 4.395, 6.233, 6.595, 7.340, 7.9, 8.549 |
| GO:0005506 | iron ion binding | 113 | 18 | 4.32 | 5.3500 | 0.0000 | ZTRI_1.1603, 1.1735, 1.1783, 1.1784, 1.187, 1.834, 10.145, 10.489, 11.126, 12.61, 2.1138, 2.570, 3.103, 4.395, 6.595, 7.340, 7.9, 8.549 |
| GO:0046906 | tetrapyrrole binding | 102 | 16 | 4.26 | 5.2100 | 0.0000 | ZTRI_1.1603, 1.1735, 1.1783, 1.1784, 1.187, 1.834, 10.145, 10.489, 11.126, 12.61, 3.103, 4.395, 6.595, 7.340, 7.9, 8.549 |
| GO:0020037 | heme binding | 102 | 16 | 4.26 | 5.2100 | 0.0000 | ZTRI_1.1603, 1.1735, 1.1783, 1.1784, 1.187, 1.834, 10.145, 10.489, 11.126, 12.61, 3.103, 4.395, 6.595, 7.340, 7.9, 8.549 |
| GO:0016491 | oxidoreductase activity | 666 | 48 | 1.96 | 2.3700 | 0.0000 | ZTRI_1.1215, 1.1603, 1.1735, 1.1783, 1.1784, 1.187, 1.1950, 1.724, 1.834, 10.145, 10.35, 10.489, 10.546, 11.126, 12.28, 12.61, 13.227, 1.253, 2.10, 2.1015, 2.1087, 2.1138, 2.57, 2.570, 2.646, 2.751, 3.103, 3.1037, 3.875, 4.120, 4.395, 4.484, 4.613, 4.675, 4.724, 5.424, 6.2333, 6.247, 6.595, 7.139, 7.340, 7.431, 7.9, 8.140, 8.548, 8.549, 9.334, 9.542 |
| GO:0048037 | cofactor binding | 428 | 33 | 2.09 | 2.4200 | 0.0000 | ZTRI_1.1215, 1.1603, 1.1735, 1.1783, 1.1784, 1.187, 1.1950, 1.724, 1.834, 10.145, 10.35, 10.489, 10.546, 11.126, 12.28, 12.61, 2.10, 2.1015, 2.1087, 3.103, 4.395, 4.484, 4.613, 6.247, 6.595, 7.340, 7.9, 8.4458, 8.549, 9.334, 9.542, 9.475, 8.749, 11.273, 11.184 |
| GO:0046914 | transition metal ion binding | 420 | 30 | 1.94 | 2.1900 | 0.0003 | ZTRI_1.1162, 1.1603, 1.1735, 1.1783, 1.1784, 1.187, 1.834, 1.2033, 2.1138, 2.282, 2.370, 2.563, 2.570, 2.851, 3.103, 3.875, 4.395, 5.64, 5.757, 6.595, 7.340, 7.421, 7.578, 7.9, 8.549, 10.145, 10.489, 11.126, 11.185, 12.61 |
| GO:0046872 | metal ion binding | 562 | 31 | 1.5 | 1.6200 | 0.0134 | ZTRI_1.1162, 1.1603, 1.1735, 1.1783, 1.1784, 1.187, 1.2033, 1.834, 2.1138, 2.282, 2.370, 2.563, 2.570, 2.851, 3.103, 3.875, 4.395, 5.64, 5.757, 6.595, 7.340, 7.421, 7.578, 7.9, 8.549, 10.145, 10.489, 11.126, 11.185, 12.61, 13.308 |
| GO:0043169 | cation binding | 565 | 31 | 1.49 | 1.6100 | 0.0144 | ZTRI_1.1162, 1.1603, 1.1735, 1.1783, 1.1784, 1.187, 1.2033, 1.834, 2.1138, 2.282, 2.370, 2.563, 2.570, 2.851, 3.103, 3.875, 4.395, 5.64, 5.757, 6.595, 7.340, 7.421, 7.578, 7.9, 8.549, 10.145, 10.489, 11.126, 11.185, 12.61, 13.308 |
| GO:0050660 | flavin adenine dinucleotide binding | 125 | 10 | 2.17 | 2.3400 | 0.0164 | ZTRI_1.1950, 1.724, 10.35, 2.10, 2.1015, 2.1087, 4.484, 4.613, 9.475, 9.542 |
| GO:0055085 | transmembrane transport | 506 | 41 | 2.2 | 2.6500 | 0.0000 | ZTRI_1.1859, 1.1864, 1.726, 10.85, 2.1016, 2.1018, 2.1168, 2.328, 3.207, 3.224, 3.642, 3.672, 3.731, 3.82, 4.179, 4.204, 4.255, 4.459, 4.621, 4.645, 4.723, 4.879, 5.184, 5.511, 5.516, 5.61, 5.637, 6.6, 6.83, 7.15, 7.154, 7.843, 8.278, 8.286, 8.537, 11.146, 11.264, 12.259, 13.195, 13.32, 13.39 |
| GO:0055114 | oxidation-reduction process | 600 | 40 | 1.81 | 2.0900 | 0.0001 | ZTRI_1.1215, 1.1603, 1.1735, 1.1783, 1.1784, 1.187, 1.1950, 1.724, 1.834, 10.145, 10.35, 10.489, 10.546, 11.126, 12.61, 13.227, 2.10, 2.1015, 2.1087, 2.1138, 2.57, 2.570, 3.103, 3.1037, 3.875, 4.120, 4.395, 4.484, 4.613, 4.724, 5.424, 6.233, 6.247, 6.595, 7.340, 7.9, 8.548, 8.549, 9.334, 9.542 |
| GO:0006810 | transport | 677 | 42 | 1.68 | 1.9300 | 0.0004 | ZTRI_1.1193, 1.1859, 1.1864, 1.726, 10.85, 2.1016, 2.1018, 2.1168, 2.328, 3.207, 3.224, 3.642, 3.672, 3.731, 3.82, 4.179, 4.204, 4.255, 4.459, 4.621, 4.645, 4.723, 4.879, 5.184, 5.511, 5.516, 5.61, 5.637, 6.6, 6.83, 7.15, 7.154, 7.843, 8.278, 8.286, 8.537, 11.146, 11.264, 12.259, 13.195, 13.32, 13.39 |
| GO:0051234 | establishment of localization | 679 | 42 | 1.68 | 1.9200 | 0.0004 | ZTRI_1.1193, 1.1859, 1.1864, 1.726, 10.85, 2.1016, 2.1018, 2.1168, 2.328, 3.207, 3.224, 3.642, 3.672, 3.731, 3.82, 4.179, 4.204, 4.255, 4.459, 4.621, 4.645, 4.723, 4.879, 5.184, 5.511, 5.516, 5.61, 5.637, 6.6, 6.83, 7.15, 7.154, 7.843, 8.278, 8.286, 8.537, 11.146, 11.264, 12.259, 13.195, 13.32, 13.39 |
| GO:0051179 | localization | 690 | 42 | 1.65 | 1.8800 | 0.0005 | ZTRI_1.1193, 1.1859, 1.1864, 1.726, 10.85, 2.1016, 2.1018, 2.1168, 2.328, 3.207, 3.224, 3.642, 3.672, 3.731, 3.82, 4.179, 4.204, 4.255, 4.459, 4.621, 4.645, 4.723, 4.879, 5.184, 5.511, 5.516, 5.61, 5.637, 6.6, 6.83, 7.15, 7.154, 7.843, 8.278, 8.286, 8.537, 11.146, 11.264, 12.259, 13.195, 13.32, 13.39 |
| GO:0051704 | multi-organism process | 2 | 2 | 27.13 | Infinity | 0.0014 | ZTRI_1.1979, 11.307 |
| GO:0016021 | integral component of membrane | 523 | 36 | 1.87 | 2.1400 | 0.0001 | ZTRI_1.1864, 1.726, 10.85, 2.1016, 2.1018, 2.1138, 2.1168, 2.328, 3.224, 3.41, 3.642, 3.672, 3.731, 4.179, 4.204, 4.255, 4.270, 4.621, 4.645, 4.723, 5.184, 5.511, 5.594, 5.61, 5.637, 5.720, 6.6, 6.83, 7.15, 7.154, 8.278, 8.286, 8.537, 11.264, 13.195, 13.32, 13.39 |
| GO:0031224 | intrinsic component of membrane | 523 | 36 | 1.87 | 2.1400 | 0.0001 | ZTRI_1.1864, 1.726, 10.85, 2.1016, 2.1018, 2.1138, 2.1168, 2.328, 3.224, 3.41, 3.642, 3.672, 3.731, 4.179, 4.204, 4.255, 4.270, 4.621, 4.645, 4.723, 5.184, 5.511, 5.594, 5.61, 5.637, 5.720, 6.6, 6.83, 7.15, 7.154, 8.278, 8.286, 8.537, 11.264, 13.195, 13.32, 13.39 |
| GO:0016020 | membrane | 819 | 47 | 1.56 | 1.7800 | 0.0009 | ZTRI_1.1859, 1.1864, 1.726, 10.85, 2.1016, 2.1018, 2.1138, 2.1168, 2.328, 3.207, 3.224, 3.41, 3.642, 3.672, 3.731, 4.179, 4.204, 4.255, 4.270, 4.459, 4.621, 4.645, 4.723, 4.879, 5.149, 5.184, 5.511, 5.516, 5.594, 5.61, 5.637, 5.720, 6.6, 6.83, 7.15, 7.154, 7.843, 8.278, 8.286, 8.537, 11.146, 11.264, 11.307, 12.259, 13.195, 13.32, 13.39 |
| GO:0044425 | membrane part | 603 | 37 | 1.66 | 1.8700 | 0.0011 | ZTRI_1.1864, 1.726, 10.85, 2.1016, 2.1018, 2.1138, 2.1168, 2.328, 3.207, 3.224, 3.41, 3.642, 3.672, 3.731, 4.179, 4.204, 4.255, 4.270, 4.621, 4.645, 4.723, 5.184, 5.511, 5.594, 5.61, 5.637, 5.720, 6.6, 6.83, 7.15, 7.154, 8.278, 8.286, 8.537, 11.264, 13.195, 13.32, |

All of the genes associated with enriched GO-terms involved in haem binding or tetrapyrrole binding are CYPs (cytochrome P450 oxidases). Many of these same CYP genes are associated with GO terms relating to iron ion binding or oxidation/reduction. Also associated with oxidation/reduction GO terms were two sphingolipid fatty acid hydroxylases (ZTRI_2.1138, 2.570). These and other over-represented GO-terms were associated with a range of genes with annotations that only arose once in the results. To better understand any commonalities in function between these genes, their PFAM domains were investigated and the number of times each domain occurs in gene products associated with enriched GO-terms is shown in Figure 10. Domain descriptions are given in Table S6. This analysis indicates that there are 12 PFAM domains that occur in >1 of these gene products. The most common domain is PF0067 (Cytochrome p450; 15 gene products), followed by PF00264 (Tyrosinase; 4 gene products). Each of PF00743 (flavin-binding monooxygenase), PF00550 (Phosphopantetheine attachment site), PF07732 (Cu-oxidase-3) and PF01266 (DAO FAD dependent oxidoreductase) occur in 3 gene products.

**Figure 10.**
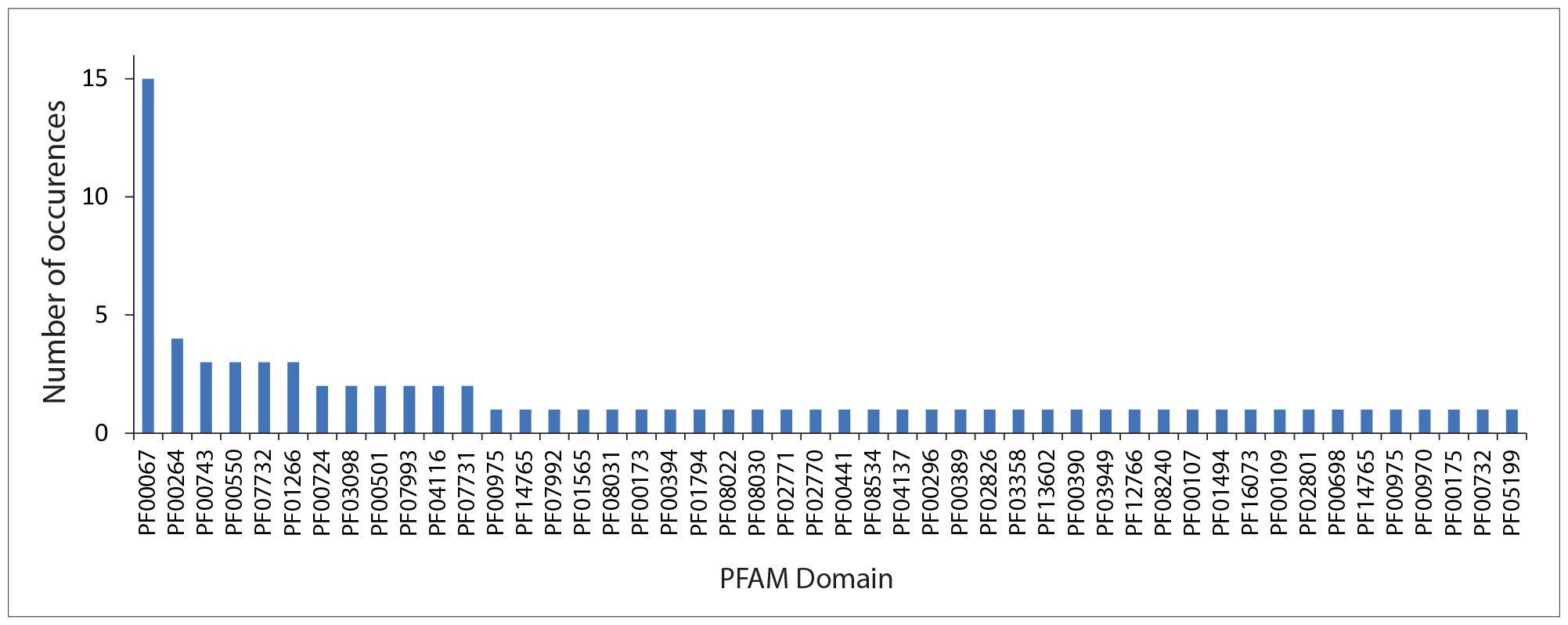
PFAM domains associated with over-represented GO terms enrichment among genes uniquely up-regulated after 7 d in water. GO enrichment carried out using tools in FungiDB [25]. PFAM domains associated with the 52 genes located on core chromosomes and associated with significantly enriched GO terms (Bonferroni corrected P<0.05) were investigated. A list of PFAM domains (49 domains) was manually curated by inspection of the results of FungiDB’s InterPro Domain search for each gene product. The number of occurrences of each domain is shown.

Searching the list of genes uniquely up-regulated after 7 days in water for the string ‘transcr*’ returned eleven genes (Table S7). Predicted domains in all eleven were fungal specific, with ten predictions being ‘fungal Zn(2)-Cys(6) binuclear cluster domain’, suggesting that they are zinc cluster proteins, fungal-specific regulators of metabolism and stress response.

### Genes up-regulated at all time points

There are 108 genes that are up-regulated at every time point. These are found largely on the core chromosomes, with chromosomes 15 and 19 also represented. As for the seven-day time point, GO terms enriched among genes up-regulated at all times (Table 4) include tetrapyrrole/haem/iron ion binding (GO:0046906, GO:0020037, GO:0005506), and these, once again, are associated with genes annotated as CYP-450s. Flavin oxygenases also appear (GO:0016712), along with more general oxidation-reduction GO terms (GO:0016899, GO:0016491, GO:0016705, GO:0055114) as does transmembrane transport (GO:0055085). GO terms that are enriched only in the set of genes up-regulated at all time points include dolichyl-phosphate-mannose-glycolipid alpha-mannosyltransferase activity (GO:0004584; P = 0.008); long-chain-alcohol oxidase activity (GO:0046577; P = 0.008); aromatase activity (GO:0070330; P = 0.015) and acyl-CoA oxidase activity (GO:0003997; P = 0.015). A number of enriched GO terms also relate to lipid metabolism: fatty acid metabolic process (GO:0006631; P = 0.0043), lipid catabolic process (GO:0016042; P = 0.0058), fatty acid beta-oxidation (GO:0006635, P = 0.015), lipid oxidation (GO:0034440; P = 0.015) and fatty acid oxidation (GO:0019395; P = 0.015). This gene set contains no transcription factors.

**Table 4:** GO terms significantly enriched in genes up-regulated at all time-points investigated. Enrichment analysis carried out using tools in FungiDB [25] P-value threshold for inclusion is 0.02.

| GO ID | GO Term | Genes in background with this term | Genes up-regulated at all time points | Fold enrichment | Odds ratio | P-value | Gene IDs |
| --- | --- | --- | --- | --- | --- | --- | --- |
| GO: 0046906 | tetrapyrrole binding | 102 | 6 | 7.59 | 9.2 | 0.0001 | ZTR1_1.153, 4.167, 5.122, 7.432, 7.458, 9.626 |
| GO: 0020037 | heme binding | 102 | 6 | 7.59 | 9.2 | 0.0001 | ZTR1_1.153, 4.167, 5.122, 7.432, 7.458, 9.626 |
| GO: 0016491 | oxidoreductase activity | 666 | 14 | 2.71 | 3.66 | 0.0003 | ZTR1_1.153, 1.194, 1.498, 1.542, 12.262, 2.1028, 4.167, 4.168, 4.169, 4.646, 5.122, 7.432, 7.458, 9.626 |
| GO: 0055114 | oxidation-reduction process | 600 | 13 | 2.8 | 3.69 | 0.0004 | ZTR1_1.153, 1.494, 1.498, 1.542, 12.262, 2.028, 4.167, 4.168, 4.169, 4.646, 7.432, 7.458, 9.626 |
| GO: 0055085 | transmembrane transport | 506 | 10 | 2.55 | 3.09 | 0.0043 | ZTR1_1.1497, 10.200, 11.214, 11.277, 2.813, 3.156, 4.5, 5.179, 8.407, 8.748 |
| GO: 0006631 | fatty acid metabolic process | 13 | 2 | 19.85 | 24.42 | 0.0043 | ZTR1_12.262, 4.169 |
| GO: 0048037 | cofactor binding | 428 | 9 | 2.71 | 3.24 | 0.0046 | ZTR1_1.153, 12.262, 2.1028, 3.830, 4.167, 5.122, 7.432, 7.458, 9.626 |
| GO: 0016042 | lipid catabolic process | 15 | 2 | 17.2 | 20.65 | 0.0058 | ZTR1_12.262, 6.239 |
| GO: 0046577 | long-chain-alcohol oxidase activity | 1 | 1 | 129.02 | Infinity | 0.0078 | ZTR1_2.1028 |
| GO: 0004584 | dolichyl-phosphate-mannose-glycolipid alpha-mannosyltransferase activity | 1 | 1 | 129.02 | Infinity | 0.0078 | ZTR1_3.314 |
| GO: 0005506 | iron ion binding | 113 | 4 | 4.57 | 5.1 | 0.0109 | ZTR1_1.153, 4.167, 7.432, 7.458, 9.626 |
| GO: 0016899 | oxidoreductase activity, acting on the CH-OH group of donors | 2 | 1 | 64.51 | 131.2 | 0.0154 | ZTR1_2.1028 |
| GO: 0070330 | aromatase activity | 2 | 1 | 64.51 | 131.2 | 0.0154 | ZTR1_7.458 |
| GO: 0003997 | acyl-CoA oxidase activity | 2 | 1 | 64.51 | 131.2 | 0.0154 | ZTR1_12.262 |
| GO: 0016712 | oxidoreductase activity, acting on paired donors | 2 | 1 | 64.51 | 131.2 | 0.0154 | ZTR1_7.458 |
| GO: 0006635 | fatty acid beta-oxidation | 2 | 1 | 64.51 | 131.2 | 0.0154 | ZTR1_12.262 |
| GO: 0006777 | Mo-molybdopterin cofactor biosynthetic process | 2 | 1 | 64.51 | 131.2 | 0.0154 | ZTR1_13.333 |
| GO: 0034440 | lipid oxidation | 2 | 1 | 64.51 | 131.2 | 0.0154 | ZTR1_12.262 |
| GO: 0019395 | fatty acid oxidation | 2 | 1 | 64.51 | 131.2 | 0.0154 | ZTR1_12.262 |
| GO: 0019720 | Mo-molybdopterin cofactor metabolic process | 2 | 1 | 64.51 | 131.2 | 0.0154 | ZTR1_13.333 |
| GO: 0016705 | oxidoreductase activity, acting on paired donors, with incorporation or reduction of molecular oxygen | 126 | 4 | 4.1 | 4.54 | 0.0157 | ZTR1_1.153, 4.167, 7.432, 7.458 |

### Genes down-regulated uniquely at 1 h (vs YPD)

The GO-terms most enriched among genes down-regulated only after 1 h in water included 6 related to oxidation-reductase activity. There are also five GO terms associated with sphingolipid metabolism and eight relating to meiosis, karyogamy and related processes (Table 5). Most other enriched GO-terms in this set have roles in primary metabolism - including peptidase activity, mannosidase activity and carbohydrate binding (Table 5). The ten genes most strongly down-regulated at this time point are all annotated only as encoding predicted or hypothetical proteins (Table S8). Searching the list of genes uniquely down-regulated at 1 h for the string ‘transcr*’ in FungiDB [25] returned fourteen genes (Table S9). Annotations of homologous genes in *Neurospora crassa, Aspergillus flavus* and *Magnaporthe oryzae* reveal little about the downstream genes likely regulated by these transcription factors. Exceptions are ZTRI_2.860, which has homology to the *amdx* gene in *Aspergillus flavus* and *Magnaporthe oryzae* and ZTRI_-10.332, which has homology to *medA* in *A. flavus* and *M. oryzae*. AmdX regulates the expression of *amdS*, an acetamidase in *Aspergillus nidulans* that allows acetamide to be used as a source of either nitrogen or carbon [30, 31], and *medA* is known to be involved in adhesion to surfaces, biofilm formation, virulence and conidiogenesis [32]. The metabolic pathways significantly enriched among the genes uniquely down-regulated after one hour in water are shown in Table S10.

**Table 5:** GO terms significantly enriched in genes down-regulated uniquely after 1 h in water. Enrichment analysis carried out using tools in FungiDB [25] P-value threshold for inclusion is 0.02.

| GO ID | GO Term | Genes in background with this term | Genes uniquely down-regulated at 1 h | Fold enrichment | Odds ratio | P-value | Gene IDs |
| --- | --- | --- | --- | --- | --- | --- | --- |
| GO:0016705 | oxidoreductase activity, acting on paired donors, with incorporation or reduction of molecular oxygen | 129 | 9 | 5.42 | 6.35 | 0.0000 | ZTRI_1.1068, 1.1950, 1.549, 1.828, 3.31, 5.698, 7.219, 9.359, 10.506 |
| GO:0003824 | catalytic activity | 2985 | 55 | 1.43 | 2.37 | 0.0001 | ZTRI_1.1068, 1.1244, 1.1310, 1.1376, 1.1387, 1.1605, 1.1759, 1.1822, 1.1950, 1.549, 1.764, 1.828, 2.1193, 2.323, 2.525, 2.63, 2.96, 3.1013, 3.31, 3.368, 4.700, 4.732, 4.95, 5.341, 5.575, 5.698, 6.150, 7.219, 7.234, 7.431, 7.439, 8.140, 8.254, 10.193, 10.212, 10.262, 10.273, 10.375, 10.400, 10.429, 10.494, 10.506, 10.65, 11.122, 12.330, 12.438, 13.174, 13.198, 14.84 |
| GO:0004497 | monooxygenase activity | 91 | 7 | 5.98 | 6.9 | 0.0002 | ZTRI_1.1068, 1.1950, 1.828, 5.698, 7.219, 9.359, 12.330 |
| GO:0016709 | oxidoreductase activity, acting on paired donors, with incorporation or reduction of molecular oxygen, NAD(P)H as one donor, and incorporation of one atom of oxygen | 25 | 4 | 12.43 | 15.31 | 0.0003 | ZTRI_1.1068, 1.1950, 5.698, 7.219 |
| GO:0050661 | NADP binding | 39 | 4 | 7.97 | 9.17 | 0.0015 | ZTRI_1.1068, 1.1950, 7.219, 10.193 |
| GO:0016491 | oxidoreductase activity | 781 | 20 | 1.99 | 2.35 | 0.0017 | ZTRI_1.1068, 1.1950, 1.549, 1.828, 2.63, 3.31, 4.732, 5.698, 6.150, 7.219, 7.234, 7.431, 8.140, 8.548, 9.359, 10.193, 10.400, 10.494, 10.506, 12.330 |
| GO:0004499 | N,N-dimethylaniline monooxygenase activity | 21 | 3 | 11.1 | 13.24 | 0.0023 | ZTRI_1.1068, 1.1950, 7.219 |
| GO:0004222 | metalloendopeptidase activity | 32 | 3 | 7.28 | 8.2 | 0.0078 | ZTRI_2.1193, 2.252, 9.112 |
| GO:0030246 | carbohydrate binding | 34 | 3 | 6.86 | 7.67 | 0.0092 | ZTRI_1.549, 10.273, 10.429 |
| GO:0004517 | nitric-oxide synthase activity | 1 | 1 | 77.69 | Infinity | 0.0129 | ZTRI_5.698 |
| GO:0004567 | beta-mannosidase activity | 1 | 1 | 77.69 | Infinity | 0.0129 | ZTRI_1.1605 |
| GO:0004767 | sphingomyelin phosphodiesterase activity | 1 | 1 | 77.69 | Infinity | 0.0129 | ZTRI_10.375 |
| GO:0004348 | glucosylceramidase activity | 1 | 1 | 77.69 | Infinity | 0.0129 | ZTRI_3.1013 |
| GO:0090729 | toxin activity | 1 | 1 | 77.69 | Infinity | 0.0129 | ZTRI_1.1979 |
| GO:0006684 | sphingomyelin metabolic process | 1 | 1 | 76.74 | Infinity | 0.0130 | ZTRI_10.375 |
| GO:0048288 | nuclear membrane fusion involved in karyogamy | 1 | 1 | 76.74 | Infinity | 0.0130 | ZTRI_4.159 |
| GO:0000741 | karyogamy | 1 | 1 | 76.74 | Infinity | 0.0130 | ZTRI_4.159 |
| GO:1990456 | mitochondrion-endoplasmic reticulum membrane tethering | 1 | 1 | 76.74 | Infinity | 0.0130 | ZTRI_2.459 |
| GO:0000740 | nuclear membrane fusion | 1 | 1 | 76.74 | Infinity | 0.0130 | ZTRI_4.159 |
| GO:0006685 | sphingomyelin catabolic process | 1 | 1 | 76.74 | Infinity | 0.0130 | ZTRI_10.375 |
| GO:0070096 | mitochondrial outer membrane translocase complex assembly | 1 | 1 | 76.74 | Infinity | 0.0130 | ZTRI_2.459 |
| GO:0090173 | regulation of synaptonemal complex assembly | 1 | 1 | 76.74 | Infinity | 0.0130 | ZTRI_1.709 |
| GO:0060629 | regulation of homologous chromosome segregation | 1 | 1 | 76.74 | Infinity | 0.0130 | ZTRI_1.709 |
| GO:0060631 | regulation of meiosis I | 1 | 1 | 76.74 | Infinity | 0.0130 | ZTRI_1.709 |
| GO:0000742 | karyogamy involved in conjugation with cellular fusion | 1 | 1 | 76.74 | Infinity | 0.0130 | ZTRI_4.159 |
| GO:0006665 | sphingolipid metabolic process | 15 | 2 | 10.23 | 11.92 | 0.0158 | ZTRI_10.375, 3.1013 |
| GO:0051783 | regulation of nuclear division | 15 | 2 | 10.23 | 11.92 | 0.0158 | ZTRI_1.709, 1.1244 |
| GO:0005506 | iron ion binding | 115 | 5 | 3.38 | 3.65 | 0.0159 | ZTRI_1.549, 1.828, 3.31, 6.150, 9.359 |
| GO:0055114 | oxidation-reduction process | 643 | 15 | 1.79 | 1.99 | 0.0177 | ZTRI_1.1950, 1.549, 1.828, 10.193, 10.400, 10.494, 10.506, 3.31, 5.698, 6.150, 7.219, 7.234, 8.548, 9.359 |

### All genes down-regulated at 1 h in water (vs YPD)

There are a total of 492 genes significantly down-regulated after 1 h in water; 34 GO-terms are significantly enriched among them, as shown in Table 6. These encompass a diverse range of functions. Notable among these are redox functions including oxidoreductase and peroxidase activity, iron-ion and iron-chelate binding functions, transport functions and catabolism of various small molecules. The top 10 most down-regulated genes (Table S11) also include transporters, as well as a GFY plasma-membrane protein likely to function as an acetate channel [33] and an acetolactate synthase. Searching the list of all genes down-regulated at 1 h for the string ‘transcr*’ in FungiDB [25] returned twenty-three genes (Table S12). There are also over 100 metabolic pathways that are over-represented in this gene list (Table S13).

**Table 6:**
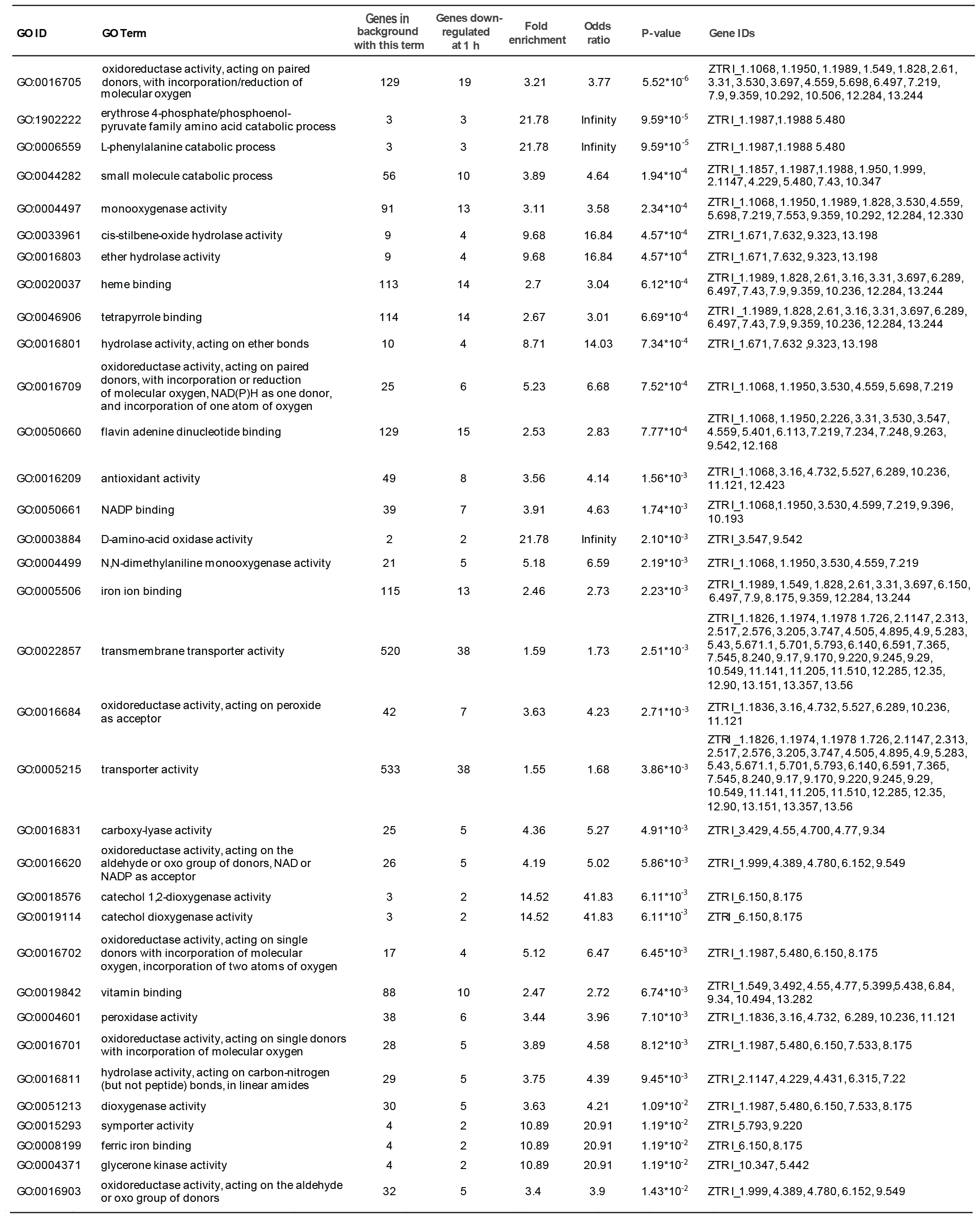
GO terms significantly enriched among all genes down-regulated after 1 h in water. Enrichment analysis carried out using tools in FungiDB [25] P-value threshold for inclusion is 0.02.

### Genes down-regulated after 7 d in water (vs YPD)

There are a total of 512 genes significantly down-regulated after 7 days in water; 272 are uniquely down-regulated at this time point. 29 GO-terms significantly are enriched among them, as shown in Table 7. Repeating motifs among these include glucan metabolism, transport and antioxidant activity. Searching the list of all genes down-regulated at 1 h for the string ‘transcr*’ in FungiDB [25] returned twenty-one genes (Table S14).

**Table 7:** GO terms significantly enriched among all genes down-regulated after 1 h in water. Enrichment analysis carried out using tools in FungiDB [25] P-value threshold for inclusion is 0.02.

| GO ID | GO Term | Genes in the bkgd with this term | Genes in result with this term | Percent of bkgd genes in result | Fold enrichment | Odds ratio | P-value |
| --- | --- | --- | --- | --- | --- | --- | --- |
| GO:0004601 | peroxidase activity | 38 | 8 | 21.1 | 11.52 | 15.32 | 3.02*10<sup>-7</sup> |
| GO:0016684 | oxidoreductase activity, acting on peroxide as acceptor | 42 | 8 | 19 | 10.42 | 13.51 | 6.85*10<sup>-7</sup> |
| GO:0016209 | antioxidant activity | 49 | 8 | 16.3 | 8.93 | 11.19 | 2.35*10<sup>-6</sup> |
| GO:0004553 | hydrolase activity, hydrolyzing O-glycosyl compounds | 130 | 10 | 7.7 | 4.21 | 4.81 | 1.19*10<sup>-4</sup> |
| GO:0016798 | hydrolase activity, acting on glycosyl bonds | 139 | 10 | 7.2 | 3.94 | 4.47 | 2.07*10<sup>-4</sup> |
| GO:0005975 | carbohydrate metabolic process | 218 | 12 | 5.5 | 3.01 | 3.38 | 5.83*10<sup>-4</sup> |
| GO:0055085 | transmembrane transport | 531 | 20 | 3.8 | 2.06 | 2.33 | 1.35*10<sup>-3</sup> |
| GO:0016491 | oxidoreductase activity | 781 | 26 | 3.3 | 1.82 | 2.1 | 1.50*10<sup>-3</sup> |
| GO:0022857 | transmembrane transporter activity | 520 | 19 | 3.7 | 2 | 2.24 | 2.56*10<sup>-3</sup> |
| GO:0006865 | amino acid transport | 17 | 3 | 17.6 | 9.66 | 11.79 | 3.35*10<sup>-3</sup> |
| GO:0005215 | transporter activity | 533 | 19 | 3.6 | 1.95 | 2.18 | 3.38*10<sup>-3</sup> |
| GO:0102250 | linear malto-oligosaccharide phosphorylase activity | 1 | 1 | 100 | 54.72 | Infinity | 1.83*10<sup>-2</sup> |
| GO:0004657 | proline dehydrogenase activity | 1 | 1 | 100 | 54.72 | Infinity | 1.83*10<sup>-2</sup> |
| GO:0004557 | alpha-galactosidase activity | 1 | 1 | 100 | 54.72 | Infinity | 1.83*10<sup>-2</sup> |
| GO:0046558 | arabinan endo-1,5-α-L-arabinosidase activity | 1 | 1 | 100 | 54.72 | Infinity | 1.83*10<sup>-2</sup> |
| GO:0102499 | SHG alpha-glucan phosphorylase activity cis-regulatory region | 1 | 1 | 100 | 54.72 | Infinity | 1.83*10<sup>-2</sup> |
| GO:0000987 | sequence-specific DNA binding | 1 | 1 | 100 | 54.72 | Infinity | 1.83*10<sup>-2</sup> |
| GO:0008184 | glycogen phosphorylase activity | 1 | 1 | 100 | 54.72 | Infinity | 1.83*10<sup>-2</sup> |
| GO:0004645 | 1,4-alpha-oligoglucan phosphorylase activity | 1 | 1 | 100 | 54.72 | Infinity | 1.83*10<sup>-2</sup> |
| GO:0046942 | carboxylic acid transport | 29 | 3 | 10.3 | 5.66 | 6.34 | 1.54*10<sup>-2</sup> |
| GO:0015849 | organic acid transport | 29 | 3 | 10.3 | 5.66 | 6.34 | 1.54*10<sup>-2</sup> |
| GO:0044419 | interspecies interaction between organisms | 11 | 2 | 18.2 | 9.95 | 12.13 | 1.63*10<sup>-2</sup> |
| GO:0009405 | pathogenesis | 11 | 2 | 18.2 | 9.95 | 12.13 | 1.63*10<sup>-2</sup> |
| GO:1902274 | positive regulation of (R)-carnitine transmembrane transport | 1 | 1 | 100 | 54.72 | Infinity | 1.83*10<sup>-2</sup> |
| GO:1902269 | positive regulation of polyamine transmembrane transport | 1 | 1 | 100 | 54.72 | Infinity | 1.83*10<sup>-2</sup> |
| GO:1902272 | regulation of (R)-carnitine transmembrane transport | 1 | 1 | 100 | 54.72 | Infinity | 1.83*10<sup>-2</sup> |
| GO:1902267 | regulation of polyamine transmembrane transport | 1 | 1 | 100 | 54.72 | Infinity | 1.83*10<sup>-2</sup> |
| GO:0006820 | anion transport | 57 | 4 | 7 | 3.84 | 4.16 | 1.99*10<sup>-2</sup> |

The metabolic pathways significantly enriched among the genes uniquely down-regulated after one hour in water are shown in Table S15. Most are related to primary metabolic functions such as fatty acid or amino acid biosythesis or degradation.

### Comparison of transcriptomic changes in water with those seen in other low nutrient environments

We compared the changes in gene expression that we observed with those seen in related studies. Rudd *et al*. (2015) [34] compared gene expression in cells on either the rich potato-dextrose broth (PDB) or the lower nutrient Czapek-Dox broth (CDB), as well as 24 h after inoculation on to a wheat leaf. Kilaru *et al*. (2022) [35] identified genes differentially regulated during the switch to hyphal growth across all isolates of *Z. tritici* investigated, which they termed Pan-strain Core Dimorphism Genes (PCDGs). Conditions found to induce hyphal growth and thus alterations in PCDG expression included low nutrient availability (minimal media) and the addition of host cues to the growth medium in the form of wheat leaf surface extracts (WLSEs). Using published lists of differentially expressed genes in those studies [34, 35], we carried out GO term analysis in FungiDB [25]. We then compared the numbers of genes associated with GO terms enriched in these gene lists with those associated with GO terms enriched in our lists of differentially expressed genes at 1 h or 7 days in water, or at all time points in this study. Figure 11 shows the GO terms enriched among up-regulated genes at any of the time points in our study, along with the number of genes associated with each term at each time point, in CDB [34], 24 h after inoculation onto the wheat leaf surface [34], in PCDGs [35], or in response to WLSE [35]. Figure 12 shows the same information but for GO terms enriched among down-regulated genes in each case. There are clear commonalities between the genes and GO terms that respond to each of the tested conditions, and also differences. Shared GO terms among upregulated genes are largely associated with redox, co-factor binding and primary metabolism, which the greatest similarities appearing between our 1 h time point and the conditions tested by Kilaru *et al*. [35]. For down-regulated genes, the shared GO terms are largely associated with redox, cofactor binding, transport, membranes and primary metabolism, with our 1 h time point most closely resembling WLSE [35] and 7 days having most in common with the CDB condition tested by Rudd *et al*. [34].

**Figure 11.**
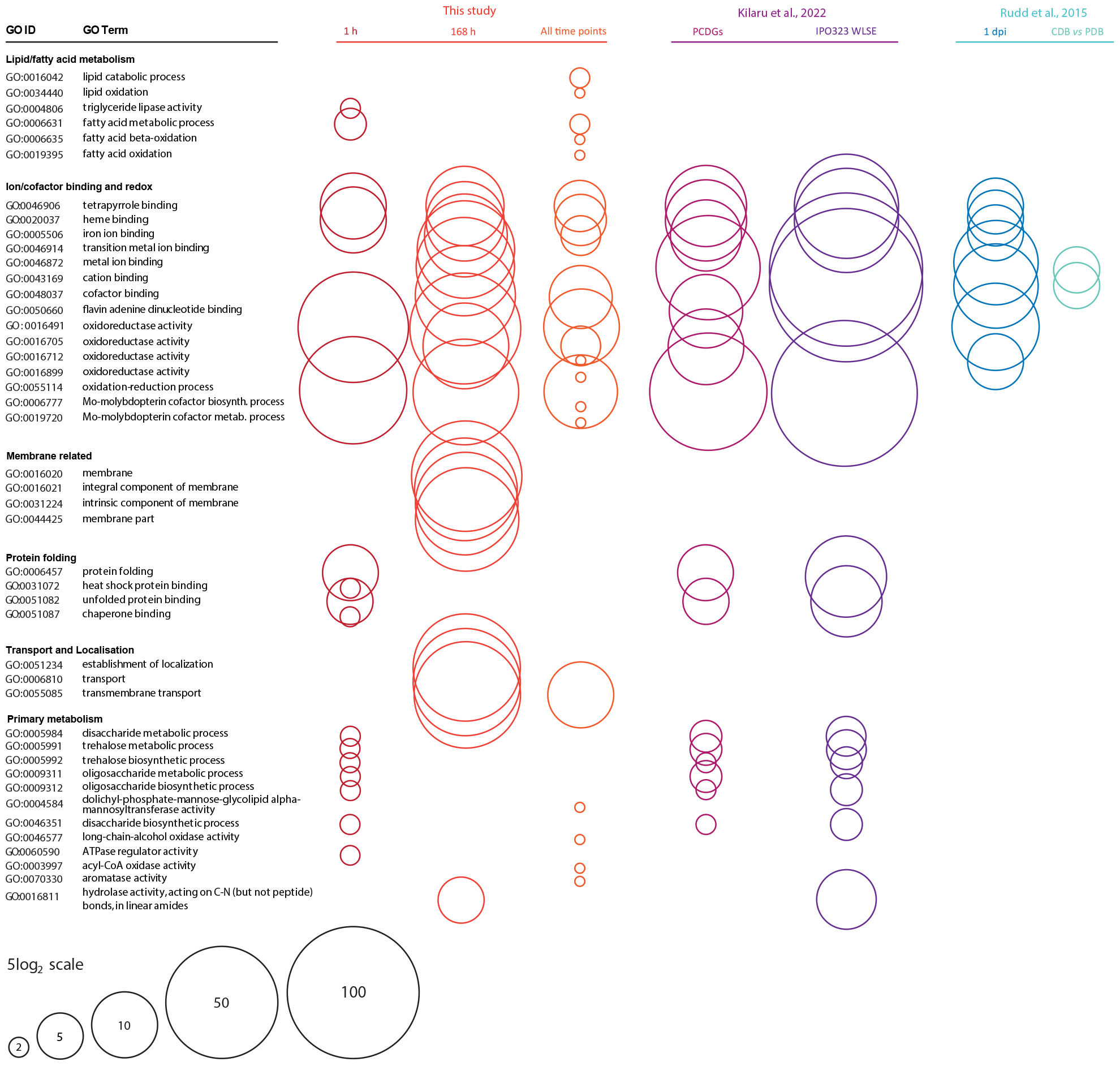
Comparison of numbers of genes related to GO terms enriched in the lists of genes up-regulated in this and studies with related, low nutrient conditions. GO terms enriched in the lists of genes up-regulated at either 1 h, 7 d or all time points in this study are shown on the left. Circle size represents the number of genes in the gene list associated with each Go term. Also shown are the numbers of genes associated with the same GO terms in other studies, if that GO term is also enriched in the relevant gene list. Studies and conditions included are: Rudd *et al*., 2015 [34] - genes up-regulated in cells at 1 day post inoculation onto wheat leaves vs *in vitro* growth on CDB and in cells grown on CDB (low nutrient) for 5 days vs PDB (high nutrient) for 3 days; Kilaru *et al*. (2022) [35] - genes identified as pan-strain core dimorphism genes up-regulated in hyphae, and genes up-regulated in cells grown for 2-3 days on minimal media + wheat leaf surface extract vs controls without extract.

**Figure 12.**
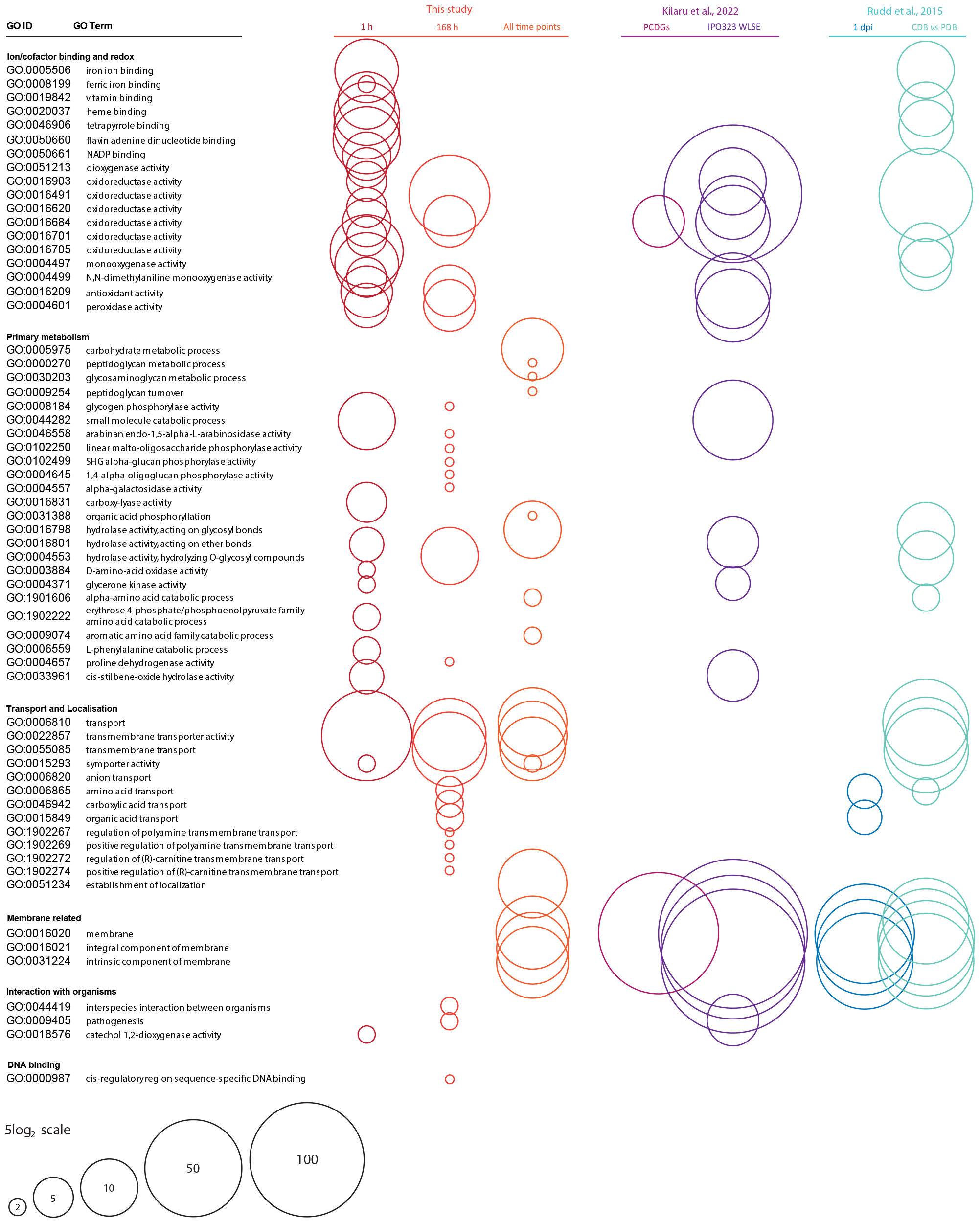
Comparison of numbers of genes related to GO terms enriched in the lists of genes down-regulated in this and studies with related, low nutrient conditions. GO terms enriched in the lists of genes down-regulated at either 1 h, 7 d or all time points in this study are shown on the left. Circle size represents the number of genes in the gene list associated with each Go term. Also shown are the numbers of genes associated with the same GO terms in other studies, if that GO term is also enriched in the relevant gene list. Studies and conditions included are: Rudd *et al*., 2015 [34] - genes down-regulated in cells at 1 day post inoculation onto wheat leaves vs *in vitro* growth on CDB and in cells grown on CDB (low nutrient) for 5 days vs PDB (high nutrient) for 3 days; Kilaru *et al*. (2022) [35] - genes identified as pan-strain core dimorphism genes down-regulated in hyphae, and genes down-regulated in cells grown for 2-3 days on minimal media + wheat leaf surface extract vs controls without extract.

## Discussion

### Long-term survival and maintenance of virulence in the absence of external nutrients has implications for our understanding of the life-cycle and population genetics of *Zymoseptoria tritici*

In this work, we have demonstrated that the blastospores of *Z. tritici* are able to survive for long periods without external nutrition. Previously, blastospores grown on YPD have been shown to be similar to pycnidiospores with respect to histology - including indistinguishable cell walls in TEM [5], virulence [4] and response to osmotic shock [15]. There are several relevant points of comparison between *in vitro* culture of blastospores on YPD agar and *in planta* pycnidiospore production. First, pycnidiospores are extruded from pycnidia in a gel-like cirrus. Although the cirrus of *Z. tritici* has not been widely studied, that of the wheat pathogen *Parastagonospora nodorum* is known to contain proteins and sugars, provide protection against dessication and inhibit germination [14]. Like that of *P. nodorum*, the cirrus of *Z. tritici* is described as mucilaginous [13]. It is hydrophobic and thought to aid in spore dispersal once free water is encountered, aiding in spread of spores by rain-splash [13]. In *P. nodorum*, another pathogen with rain-splash dispersed conidia, the cirrus needed to be diluted to around 10% of its original concentration to allow spore germination [14]. Together, these pieces of information indicate that cirrus material is likely to undergo very significant dilution during the process of rain-splash dispersal of pycnidiospores to a new host leaf. Thus, the spores undergo a severe alteration in their environment, from extrusion in a protective and nutrient dense cirrus matrix to dilution into rainwater. This process includes the sudden removal of external nutrients and an osmotic shock. Thus, the transition from YPD to water, investigated here, forms a reasonable proxy for dispersion from the cirrus via rain-splash.

Given the lack of nutrients available, growth and survival might be expected to be very limited after suspension of blastospores in distilled (MilliQ) water, and we have previously seen that culturability and virulence of suspended blastospores begins to fall within minutes [15]. However, we determined that some blastospores within a population remained alive for at least 49 days (7 weeks). Our conditions were very stringent - in the field, spores will have access to at least as many, or more, nutrients as in these experiments. Thus, the ability to survive starvation for at least 49 days is likely to have relevance under field conditions. In the UK, winter wheat is generally harvested in August, and the new crop drilled between September and November, depending on weather, soil type and wheat variety [36]. Thus, a period of seven weeks will often be sufficient to span the intercrop period for wheat. Following dispersal, spores also become subject to additional stresses whilst on the leaf surface, including a risk of desiccation. However, our data indicate that desiccation does not increase spore death rates over a period of 28 days, which reduces the likelihood that such stresses invalidate the long-term spore survival that we propose here. In line with this, Chaloner *et al*. (2019) [37] showed that after a small drop in viability associated with the drying process, proportion of viable spores was not greatly affected by 4 h of dryness. An increase in the death rate of dried spores was observed with increasing temperature, but significant differences between wet and dry spore viability were only seen at temperatures above the average for the UK in the intercrop period for wheat [37, 38]. Moreover, we show here that spore survival in soils is comparable to that seen in water. This is important both because wheat plants or stubble will not necessarily be available post-harvest, depending on the management of the field, and because soils retain more moisture and absorb UV light, mitigating the affects of non-starvation stresses. In addition, soil may be carried between fields on farm equipment and may thus be a mechanism for introducing *Z. tritici* spores into fields in which wheat is about to be sown. We also demonstrate that, once adjusted for the drop in viability associated with long-term storage, spores stored in either water or soil for 49 days retain their virulence, and that infection of wheat seedlings could occur as a result of rain-splash dispersal of 49 day old spores from soil. Taken together, these findings raise the possibility that soil-borne, asexual pycnidiospores, as well as wind-blown, sexual ascospores, can act as primary inoculum for outbreaks of Septoria tritici blotch. This may be relevant to disease management. Coupled with the production of blastospores *in planta* by virulent [5] and avirulent [6] *Z. tritici*, the long-term survival of blastospores in soils and low-nutrient environments such as non-host leaves provides a route by which genotypes less able to infect a given crop may remain in the population and potentially contribute genes to later generations via sexual reproduction or vegetative fusion. This has potential impact on the best strategies for managing the emergence of strains virulent on previously resistant wheat cultivars or resistant to fungicides. The ability of *Z. tritici* blastospores to survive desiccation also raises the possibility of aerial dispersal, for which this ability is essential. Like epiphytic bacteria, leaf surface fungi can become airborne [39, 40], providing an alternative route to dispersal. We note that if this were to occur with *Z. tritici*, for example following epiphytic blastosporulation or from leaf surface biofilms [5, 6, 8] our data suggest that desiccated spores would not use their lipid supplies, meaning that if rained out of the atmosphere onto a suitable host, they would still contain enough lipids to support colonisation. This possibility, to our knowledge, has never been explored in *Z. tritici*, whose ascospores are considered responsible for wind-blown dispersal [41, 42].

### Use of nutrient stores and changes to primary metabolism likely underpin survival in water

The data presented here indicate that *Z. tritici* blastospores rely on stored lipids for energy during starvation. Cells are seen packed with lipid droplets after the production of blastospores on YPD and these lipids become depleted over time in starvation conditions. Lipid content was also significantly positively correlated with spore viability, suggesting that as lipid stores become depleted, the probability of cell death rises. Trehalose and glycogen, by contrast, were either not present or not significantly depleted during starvation, indicating that these are not relied upon as nutrient sources under these conditions. Both are known to be important carbohydrate stores in yeasts such as *Saccharomyces cerevisiae* [21]. In *Aspergillus nidulans*, trehalose is a major storage compound in spores, and is rapidly metabolised following germination, with conidial viability in storage rapidly lost in trehalose biosynthesis mutants [22]. However, in many fungi, trehalose is mainly produced under stressful conditions, including those that trigger sporulation in many fungi, such as starvation [43]. The fact that *Z. tritici* blastospores raised on YPD do not accumulate trehalose is therefore consistent with previous findings in other fungi. The lack of trehalose production during subsequent starvation may indicate that *Z. tritici* does not accumulate trehalose in response to this stress, or that this takes place beyond the 8 day timepoint at which we ended trehalose measurement.

Intriguingly, GO terms and metabolic pathways associated with trehalose synthesis were enriched in the set of genes up-regulated after an hour in water. Genes linked to these terms included trehalose-6-phosphatase (T6Pase) and T6Pase synthase, supporting an increase in flux through the trehalose biosynthesis pathway. Since little or none is accumulated, this might be linked to rapid trehalose turnover. Additional functions of trehalose are reported to include temperature, oxidative and osmotic stress tolerance and signalling, as well as the virulence of several phytopathogens [22, 44–47]. Further, since trehalose synthesis begins with the conversion of glucose-6-phosphate to trehalose-6-phosphate (T-6-P), trehalose and T-6-P concentrations can be involved in the regulation of glycolysis [21]. Thus, up-regulation of trehalose synthesis may be related to survival during starvation in an indirect manner.

From the results presented here it is at least clear that neither glycogen nor trehalose stores provide energy for long-term blastospore survival in water. While pycnidiospores might in theory contain different storage compounds to blastospores, we consider that their similar histology and lipid content [5], germination behaviour *in planta* [4], virulence [23] and response to osmotic shock [15] makes large differences unlikely. Together, these results therefore suggest that lipid accumulation is key to long-term survival in low- or no-nutrient environments. From this finding, we hypothesise that the ability of *Z. tritici* to germinate, survive and even proliferate on the low-nutrient leaf surface [4–6, 8] is also likely to show strong dependence on prior lipid accumulation in spores. In this context, the finding that dry spores do not use their lipids, despite not showing a reduction in viability, is surprising. Moreover, desiccation survival in *Saccharomyces cerevisiae* is dependent on lipid breakdown [48]. This suggests that *Z. tritici* may have an as yet un-elucidated mechanism for surviving both starvation and desiccation. Perhaps most likely is a form of metabolic arrest, as seen during the desiccation of micro-colonial fungi on rock surfaces [49, 50].

Gene expression patterns suggest that the most significant changes undergone by *Z. tritici* cells under starvation are, indeed, linked to a global down-regulation of metabolism. After only 1 h in water, genes involved in primary metabolic functions such as peptidase activity were already significantly over-represented among down-regulated genes. At 1 h, around 10x more metabolic pathways were significantly over-represented among down-regulated genes, compared to up-regulated genes. Despite this, oligo- and disaccharide biosynthesis is up-regulated after 1 h in water, along with lipase activity and pathways relating to fatty acid *β*-oxidation and salvage. GO-terms relating to lipid and fatty acid metabolism are also enriched among up-regulated genes at all time points, which is consistent with the observed depletion of lipid droplets and our hypothesis concerning the importance of stored lipids for survival in water. After 7 days in water, further GO terms associated with primary metabolism, including carbohydrate, sugar, amino acid, carboxylic acid and polyamine catabolism, are down-regulated. Overall, gene expression results confirm a reduction in overall metabolic activity coupled with a likely reliance on stored lipids for energy. The transcriptome of dried cells would make an interesting point of comparison here.

### Relationship between cells in an individual during long-term survival

A striking feature of the *Z. tritici* spore population after an extended period of starvation was the observed reduction in average cells per spore, to a minimum of one after 49 days. This appeared to be due in part to budding reproduction, but primarily to the death of cells within the spore, which we propose is likely to lead to spores splitting into smaller units of fewer cells. The increase in total spore counts supports the hypotheses of either budding or breaking, with the decline in this measure from 42 to 49 days indicating that lysis of dead cells is also occurring, and outweighs the rate of budding and splitting once cells no longer have the resources to grow and are already composed of only one cell, which cannot split further. The death of cells within multicellular spores, while the remaining cells remain healthy, raises some important questions about the interactions between the cells of an individual spore. In common with other ascomycete fungi, *Z. tritici* is functionally coencytic, since the septae which divide the cells are perforated by pores and the cytoplasm is therefore continuous [51, 52]. These pores are kept open in an ATP-dependent manner and are blocked by Woronin bodies to prevent cytoplasmic bleeding if a cell is wounded [53, 54]. This explains how a spore can survive the death of certain cells within it. However, it remains unclear why cell death is triggered in some cells but not others under the same conditions. *Z. tritici* blastospore populations have been shown to behave in a highly asynchronous manner following inoculation onto wheat leaves [4, 9]. The findings here raise the possibility that this asynchronicity also exists at the level of cells within individuals. It is possible that certain cells, due to their status at the moment of suspension in water (for example, their cell cycle stage or nutrient content) are likely to die earlier than others. Cytoplasmic connectivity means that the nutrients within these cells could theoretically be scavenged by their neighbours. In live fungal cells, early endosomes and vacuoles mediate intercellular transport and can pass through septal pores, while transporters can mediate the selective exchange of nutrients between cells, even when septal pores are occluded by Woronin bodies [55–57]. It is thus theoretically possible that nutrients are passed from dying or even dead cells into their healthier neighbours. However, against this, it is notable that in the images of cells stained with propidium iodide and BODIPY^®^ in Figure 2, lipid bodies are visible in the dead cells, meaning that an efficient programme of lipid mobilisation and nutrient exchange does not occur prior to cell death. Given the apparent importance of stored lipids as an energy source during starvation, this failure to scavenge lipids from dead cells suggests that cell-cell communication and nutrient exchange is not an adaptation for survival of starvation. Notably, the mobilisation of lipid stores is thought to be important during the early phases of plant infection in *Z. tritici*, during which nutrient uptake from the host is thought to be extremely limited [1, 34]. There are therefore some parallels between early infection and the starvation conditions imposed here, which are borne out in the observed depletion of stored lipids in this work. However, *Z. tritici* is unusual in that its response to starvation is independent of autophagy [58]. While autophagy is repressed under nutrient-replete conditions in many fungi, including ascomycetes such as *Aspergillus nidulans* [59], this is not the case in *Z. tritici* [58]. Autophagy is often a precursor to programmed cell death in ascomyetes, is required to recycle nutrients from aging or obsolete cells and structures and is often essential for phytopathogen virulence [60, 61]. Recent work by Child *et al*. (2022) demonstrated that autophagy is not required for virulence in *Z. tritici* [58]. These findings cast doubt on the idea of nutrient scavenging from dying cells. An alternative hypothesis is that cell lysis, occurring after cell death, liberates cell contents into the growth media and allows their re-uptake by healthy cells, prolonging the life of the population. Such a mechanism would likely be maladaptive in widely dispersed cells, in a population of mixed genotypes or when part of a varied microbiome whose other members could compete for released nutrients. However, *Z tritici* undergoes blastosporulation (also called microcycle condiation) on the leaf surface [5] and may develop areas of dense, clonal epiphytic growth [6, 7]. These can develop into biofilms containing a mixture of live and dead cells [8]. Thus, there may be field-relevant circumstances in which the re-uptake of lysed cell contents would be almost exclusive to clonal *Z. tritici* cells. The gene ZTRI_6.83, a putative major facilitator superfamily transporter, is among the top ten most up-regulated genes after 7 days in water, and transport (GO:0006810) is significantly over-represented among genes up-regulated at this, but not at other time points. There is a steep decline in the % live cells at around 6 dpi in Figure 1, suggesting that cell lysis could be underway and nutrients available for uptake by the 7 day time point.

### Other changes in gene expression

The clear differentiation in patterns of gene expression between 7 days in water and the three earlier time points are likely to reflect a different set of genes are involved in survival of long-term starvation compared to short term drops in nutrient availability, and the fact that the initial transition from YPD to water also entails osmotic shock. Genes which are differentially regulated at the 1 h time point but not at others are considered candidates for involvement in the osmotic stress response. In line with this idea, 6 of the 10 GO terms over-represented among genes upregulated uniquely at 1 h are cell wall or membrane related. Transporter activity is also represented among the GO terms enriched among down-regulated genes at this time point, indicating that overall transport activity is altered, rather than a simple increase in transport. Among genes uniquely down-regulated after 1 hour in water, several GO terms related to sphingolipid metabolism and catabolism are over-represented. Sphingolipids modulate membrane fluidity, but they have many further roles including in virulence and stress signalling. Their accumulation during starvation, consistent with the down-regulation of sphingolipid catabolism seen here, is linked to increased integration of amino acid permeases into the membrane and thus increase amino acid uptake [62, 63]. GO terms related to sphingolipid metabolism over-represented among genes down-regulated at 1 h also include glucosylceramidase activity. Glucosylceramide is a cell wall component associated with polar growth, hyphal production and germination [64]. Glycosylceramidase activity is responsible for removing glycosyl sidechains from ceramides, whose build up is linked to apoptosis in fungi. The downregulation of glucosylceramidase will reduce the release of ceramides from their glycosylated forms, which may help to reduce apoptotic death in response to stress [65].

Other groups of GO terms over-represented among genes down-regulated at 1 h are linked to nuclear membrane fusion, karyogamy, and meiosis, suggesting that resource allocation to reproduction is also rapidly reduced on transition to a low nutrient environment. Together, these results at the 1 h time point suggest a strengthening or remodelling of walls and membranes to withstand hypo-osmostic stress, changes in transporter activity, possibly involved in modulating cytoplasmic osmotic potential, and a cessation of processes such as germination, anastomsis, or karyogamy.

At 7 days, by contrast, the majority of over-represented GO terms among up-regulated genes are involved in oxidation-reduction processes. PFAM analysis indicated that many of the genes associated with these GO terms were cytochrome P450s (CYPs). CYPs are a superfamily of proteins found all five domains of life and thought to have arisen close to the origin of terrestrial life [66, 67]. They form a multi-component oxygenase system that is involved in a wide range of functions from synthesis of secondary metabolites to detoxification of xenobiotics, adaptation to stress and to new niches [67, 68]. In phytopathogens, they have roles in host-specificity, defence compound detoxification and virulence [67, 69, 70]. In *Saccharomyces cerevisiae* and *Candida glabrata*, CYP functions include fatty acid degradation, as well as synthesis of molecules involved in cell wall and membrane structure [71]. CYPs have previously been shown to be up-regulated under starvation stress in fungi and other organisms [72, 73]. Upregulation of CYPs in response to nitrogen limitation has been shown in both Ascomycete and Basidiomycete fungi [74]. GO terms linked to transport activity and to membrane structure are also over-represented after 7 days, suggesting a possible role for membrane remodelling in starvation tolerance. Over-represented GO terms among down-regulated genes were predominantly related to primary metabolism. Thus, differences in gene expression at the 7 day time point compared to earlier times, revealed in the heatmap in Figure 9, appear to reflect the difference between short-term membrane and wall remodelling and stress responses compared with longer-term changes in metabolism and adaptation to the new environment.

### Comparison to other studies: starvation has commonalities of gene expression with hyphal and growth in minimal media with wheat leaf surface extract, but not with responses to host cues *in planta*

We hypothesised that sudden immersion of blastospores in water following growth on the rich medium, YPD, would provide similar changes to those experienced during the process of a pynidiospore being dispersed from cirrus in rain-splash: a large drop in external osmotic potential and nutrient availability. To determine to what extent our results bear this out, we compared changes in gene expression in this study to two other studies: Rudd *et al*. (2015) [34], who compared the transcriptome of *Z. tritici* cells growing on Czapek-Dox broth (CDB), to either potato dextrose broth (PDB) or 1 day after inoculation onto the wheat leaf and Kilaru *et al*. (2022) [35], who reported genes differentially regulated during hyphal growth across a range of *Z. tritici* isolates (Pan-strain core dimorphism genes, PCDGs) and also compared the transcriptome of *Z. tritici* grown in minimal medium (MM) with and without the addition of a wheat leaf surface extract (WLSE). These studies thus both provided information about *Z. tritici* gene expression under nutrient limitation, with and without additional host cues (either by inoculation onto a wheat leaf or growth on WLSE). PCDGs are association with the transition to hyphal growth, which can occur in response to the host but also to temperature stress and nutrient limitation [35]. Comparison of GO term enrichment in lists of differentially expressed genes (DEGs) in the present work with that in lists of DEGs from these previous studies show some interesting patterns. On comparing GO term enrichment in up-regulated genes, it is clear that the greatest similarity lies between the 7 day timepoint in this study and the patterns of gene expression in Kilaru *et al*.’s PCDGs and WLSE gene lists. Among down-regulated genes, GO terms enrichment is similar for WLSE and both the 1 h and 7 day time points in this study, as well as for GO terms enriched among down-regulated genes in Rudd *et al*’s comparison of CDB to PDB. The least similarity overall was between any time point in this study and Rudd *et al*’s *in planta* transcriptome. Since the list of DEGs used in this comparison compared *in planta* gene expression to that in the low-nutrient CDB, in which *Z. tritici* grows slowly [34], it can be assumed that DEGS here reflect the fungal response to host cues such as leaf hydrophobicity, wheat-specific compounds and defence responses.

The lack of similarity in GO term enrichment between starvation and *in planta* growth, compared to that seen between starvation and minimal media + WLSE suggests that shared function in genes differentially regulated in response to starvation and to the host is linked to nutrient availability, while, by the 1 dpi time point used by Rudd *et al*., the response to the host is linked to more dynamic host responses such as production of defensive compounds in response to pathogen detection - in line with previous findings that the plant also responds to the fungus within hours of contact [75, 76]. By 7 days, the greatest similarities between GO terms in this study and the WLSE GO terms are among functions such as mono-oxygenase activity, likely to relate to CYPs. This suggests similar changes in stress response or secondary metabolite synthesis or degradation during growth on WLSE and starvation. This may indicate that some changes in response to WLSE are a response which prepares cells for the starvation that may be endured during prolonged growth on the wheat leaf surface, a possibility which may bear further investigation. The similarity between GO terms enriched among DEGs in the current study and those enriched among Kilaru *et al*.’s PCDGs suggests that the response to a change in nutrient availability includes functions related to the change to hyphal growth. The most parsimonious explanation here would be that nutrient availability has a greater role in the dimorphism of *Z. tritici* than appreciated.

### Conclusion: Importance for understanding and predicting Septoria Leaf Blotch disease

In this work, we have demonstrated that *Z. tritici* blastospores can utilise lipid stores in order to survive for long periods, spanning the intercrop period for UK winter wheat, with no external nutrition. This survival occurs both in water and in soil, and although a large proportion of spores do not survive for such an extended period, those that do remain as virulent as spores grown on rich media. Coupled with the epiphytic survival of avirulent isolates [6], this suggests that rainsplash-dispersed inoculum on wheat leaves, volunteer and field margin plants and on soil could survive between crops, regardless of their virulence on the planted wheat cultivar. This implies that early infections of newly sown wheat in September-December [36] could be begun by resident asexual inoculum as well as by wind-blown ascospores. We also showed that *Z. tritici* blastospores tolerate drying out, making it more likely that this long-term survival is feasible under field conditions. Moreover, dried spores did not use their lipid stores, indicating that drying could prolong the life of spores and aid in their distribution. Changes in primary metabolism and transport also underpin survival under starvation, and this may be linked to uptake of nutrients from dead cells, potentially following cell lysis, a mechanism which is likely to be most useful to high densities of cells in biofilms, another survival adaptation. In the very short term, transition from YPD to water also involved changes to cell wall and membrane architecture likely linked to withstanding osmotic stress. Survival of both immersion in water and especially longer term starvation are linked to the function of cytochrome P450 mono-oxygenases, which are linked to fungal stress tolerance. Our initial hypothesis was that the drop in blastospore virulence and culturability seen in the first minutes after suspension in water [15] could be required for or involved in longer term survival of starvation. It is clear from the results presented here that *Z. tritici* blastospores have the posited long-term survival ability, and possible that the changes to primary metabolism, trehalose synthesis and CYP expression represent alterations to cell biochemistry that facilitate this. Taken together, and in combination with both survival on resistant hosts [6] and biofilm formation [8], these findings indicate the *Z. tritici* has a suite of exceptional survival strategies which are likely to be important in understanding its population genetics and the selection pressures on virulence and fungicide resistance traits. In turn, this might support development of novel routes for Septoria leaf blotch control.

## Methods

### Fungal isolates used in this study

All experiments in this study used the commonly-studied ‘reference’ isolate, *Z. tritici* IPO323 [2]. IPO323 and IPO323 expressing ZtGFP[77] either cytoplasmically or at the plasma-membrane using an SSO1-GFP fusion protein (SSO1) [78] was kindly provided by Dr Sreedhar Kilaru and Prof Gero Steinberg.

### Confocal microscopy

Confocal images were obtained using a 63x oil immersion lens on a Leica SP8 microscope. GFP fluorescence from *Z. tritici* was detected at 510-530 nm using an argon laser with emission at 500 nm. Images were obtained using LAS-X software and processed as batches using macros written in Adobe Photoshop^®^.

### Assessment of cells per spore and cell live/dead status using propidium iodide

*Z. tritici* IPO323 was plated from ^-^80 °C glycerol stocks onto YPD agar 7 days before use and maintained at 18 °C before resuspending in sterile MilliQ-water at 10^7^ cfu/ml. To measure the percentage of spores with at least one live cell, propidium iodide (PI) was added to *Z. tritici* cell suspensions to a final concentration of 0.1% (w/v), and incubated for 30-60 min before aliquots were diluted by 100x and 10 *µ*l mounted on standard glass microscope slides. Cells were visualised every 24 h using a Leica SP8 confocal microscope, with excitation and detection at wavelengths at 493 and 636 nm, respectively. Fields of view were selected at random for imaging. Maximum projections were obtained from image stacks using Leica’s proprietary software. Cells were scored as dead/non-viable if PI was had completely stained the cytoplasm. The total number of cells per blastospore over time was also assessed from these images, as were the total number of spores in suspension.

### Assessing the culturability of water-suspended blastospores

*Z. tritici* IPO323 was plated from ^-^80 °C glycerol stocks onto YPD agar 7 days before use and maintained at 18 °C before resuspending in sterile MilliQ-water at 10^7^ cfu/ml. Aliquots were then diluted by 1000x before spreading 100 µl (∼1000 spores) onto YPD agar. Changes in culturability were assessed by colony counting after 7 days. Culturability was calculated as: number of colonies/number of spores plated.

### Measurement of blastospore lipid content

*Z. tritici* IPO323 was plated from ^-^80 °C glycerol stocks onto YPD agar 7 days before use and maintained at 18 °C before resuspending in sterile MilliQ-water at 10^7^ cfu/ml. To measure the percentage spore area occupied by lipids, BODIPY^®^ 493/503 (4,4-difluoro-1,3,5,7,8-pentamethyl-4-bora-3a,4a-diaza-s-indacene, Thermo Fisher, Catalogue number: D3922) was used to stain lipid granules. BODIPY^®^ was stored in DMSO at ^-^20 °C at a concentration of 1 mg/ml and added to *Z. tritici* cell suspensions to a final concentration of 10 *µ*M, along with propidium iodide (PI) counterstain at a final concentration of 0.1% (w/v). After the addition of BODIPY^®^ and PI, blastospore suspensions were visualised by confocal microscopy within 30 minutes. Images were taken using excitation/emission maxima of wavelength 493/503 nm. Lipid content was assessed as a percentage of spore area by calculating: (total area of the image made up of lipid granules / total area of the images containing fungal tissue) x 100.

### Measurement of glycogen and trehalose content

Spore populations were tested over time as per [**?** ]. Briefly, 10^8^ *Z. tritici* IPO323 blastospores were heated at 95 °C in 1M acetic acid and 0.2M Sodium acetate before treatment with either *Aspergillus niger α*-amyloglucosidase (for glycogen), porcine trehalase (for trehalose), or no enzyme (controls). Glucose liberated from each reaction was assayed using a Glucose (GO) Assay Kit (Sigma, GAGO-20). Sample optical density was measured at 540 nm using a spectrophotometer and compared against prepared glucose standards.

### Assessment of blastospore survival in dry condi tions

To assess the ability of blastospores to survive periods without water, dry Petri dishes were used. *Z. tritici* IPO323 blastospores were grown on YPD agar for 7 days. blastospores were then suspended in autoclaved MilliQ water and suspension density estimated using a haemocytometer before plating ∼1000 spores onto the dry Petri dishes. Dishes were dried for 60 minutes in a Class II Cabinet before sealing with Parafilm^®^. Individual plates were re-hydrated every 7 days for a 56-day period by flooding with 2 ml of autoclaved MilliQ water. A sterile spreader was then used to suspend blastospores in solution before a small amount was aliquoted onto YPD and incubated at 18 °C under a long-day light cycle. Survival was qualified by assessing plates for *Z. tritici* growth after 7 days. On day 28, aliquots of resuspended blastospores were stained with PI and BODIPY^®^ to assess spore viability and lipid content according to the methods given above.

### Assessment of blastospore survival in soil

For experiments concerning blastospore survival in soil, 25 g of autoclaved John Innes No. 2 soil was added to a petri dish and flooded with 5 ml of autoclaved MilliQ water. Before assessment, *Z. tritici* blastospores were grown on YPD agar for 7 days. Blastospores were then suspended in autoclaved MilliQ water, estimated using a haemocytometer, and pipetted into soil at a rate of 5 ml of 1 x 106 blastospores. Soil plates were sealed with Parafilm^®^ and incubated under standard growth cabinet conditions. Every 7 days, a sterile spreader was placed into the wet soil of an individual plate and spread onto a fresh YPD plate. Plates were incubated at 20 °C under a long-day light cycle. Survival was qualified by assessing plates for *Z. tritici* growth after 7 days.

### Wheat cultivation and inoculation

#### *Triticum aestivum* cv

Consort winter wheat (kindly provided by Nick Palmer of RAGT seeds) was grown on J. Arthur Bower’s John Innes No. 2 Compost. Compost was stored frozen at ^-^20 °C for 3 weeks before use. Two seeds were sown in each cell of a 24-cell modular seed tray containing compost, and loosely covered. Trays were then placed into a Whitefurze 38 cm gravel tray and filled with 750 ml distilled water. Plants were placed onto one of the three shelves of a Panasonic MLR-352-PE growth cabinet. A long-day light cycle (16 hours of light at 20 °C and 8 hours of darkness at 15 °C) was used with 90% relative humidity, using the maximum light setting (∼5 x 10 ∼molm^-2^s^-1^ at leaf level). Plants were left uncovered for 5 days until growth was visible above the soil and subsequently grown for at least 14, and a maximum of 21 days before use.

For all foliar applications, blastospores were suspended in autoclaved MilliQ water. Concentrations were estimated by haemocytometer and adjusted to concentrations described in each experiment. Before use, suspensions were supplemented with 0.01% (v/v) Silwet L77. Suspensions were then applied using a paint-brush to the abaxial side of fully expanded leaves, until complete coverage was obtained. Post-inoculation, all wheat plants were stored under standard growth cabinet conditions for 28 days. For the first 5 days, plants were sealed in autoclave bags to maintain maximum humidity. Plants were assessed for disease by counting pycnidia per cm^2^ of inoculated leaf tissue.

### Soil inoculations and rain-splash plant inoculation experiments

Five ml of a 10^6^ or 10^7^ cfu/ml blastospore suspension was pipetted into each cell of a 24-cell plant tray, each containing two 14-day old wheat plants. Blastospores suspensions were allowed to soak into soil for 10 minutes. To mimic rainfall, trays were watered from a height of 2 metres at a rate of 4 litres of sterile distilled water per 24-cell tray from a Haws No.14 medium rose head watering can. Disease was assessed as pycnidia per cm^2^ of leaf after 28 days incubation. The cotyledon and first two true leaves were assessed. Three controls were conducted: (i) soil with no blastospores added, (ii) plants grown in blastospore inoculated soil without the rain-splash event, and (iii) leaves inoculated by the paint brush method (above).

### RNA extractions

Blastospores were suspended at 1 x 10^7^ cfu/ml in 50 ml tubes of MilliQ water. These tubes were shaken vigorously for 10 seconds every 24 hours to maintain oxygenation. At four time-points (1 hour, 4 hours, 24 hours and 7 days), four randomly selected 50 ml tubes were centrifuged for 1 minute at 2000 x g before freezing pellets in liquid nitrogen. Pellets were subsequently ground to a powder in a pestle and mortar using liquid nitrogen and were not allowed to thaw. Total RNA extractions were carried out using the Qiagen RNeasy Kit (Cat No./ID: 74903) protocol ‘Purification of Total RNA from Plant Cells and Tissues and Filamentous Fungi’ with an on-column DNase step using Qiagen RNase-Free DNase Set (Cat No./ID: 79254). Samples were finally suspended in 50 µl RNase-free water before storing at -80 °C for analysis.

### RNA sequencing

Samples were prepared using the Illumina TruSeq Stranded mRNA kit and sequenced on a HiSeq 2500 generating between 3.3 and 5.2 million reads per sample. Adapter sequences and low quality bases (<Q22) were removed using cutadapt [79] version 2.5. Reads before and after trimming were checked for quality using FactQC Version 0.10.1 [80], and for common contaminants using FastQ Screen [81]. MultiQC [82] was used to collate and visualise the results. A subset of reads was also check with BLAST [83] against the NCBI nucleotide database for other contaminants. Results were visualised using Krona [84]. Trimmed reads were aligned using STAR [85] version 2.7.2b to the reference genomes fungiDB-45_ZtriticiIPO323. DESeq2 [86] version 1.24.0 was used to determine differentially expressed genes between time-points.

### Analysis of changes in gene expression

Lists of gene differentially up- and down-regulated at each time point were analysed for GO term and metabolic pathways enrichment using the tools available at FungiDB (https://www.fungidb.org). PFAM analysis was carried out by extracting PFAM domains associated with genes of interest manually from FungiDB’s gene pages. A global heatmap of differentially expressed genes was produced using SR plot [24]. Comparison of GO term enrichment between lists of differentially expressed genes produced in this and other studies was carried out by using the GO term enrichment analysis tools at FungiDB with our own gene lists, as before, and with gene lists provided in the data associated with Rudd *et al*. (2015) and Kilaru *et al*. (2022) [34, 35].

## Supplementary Information

**Figure S1.**
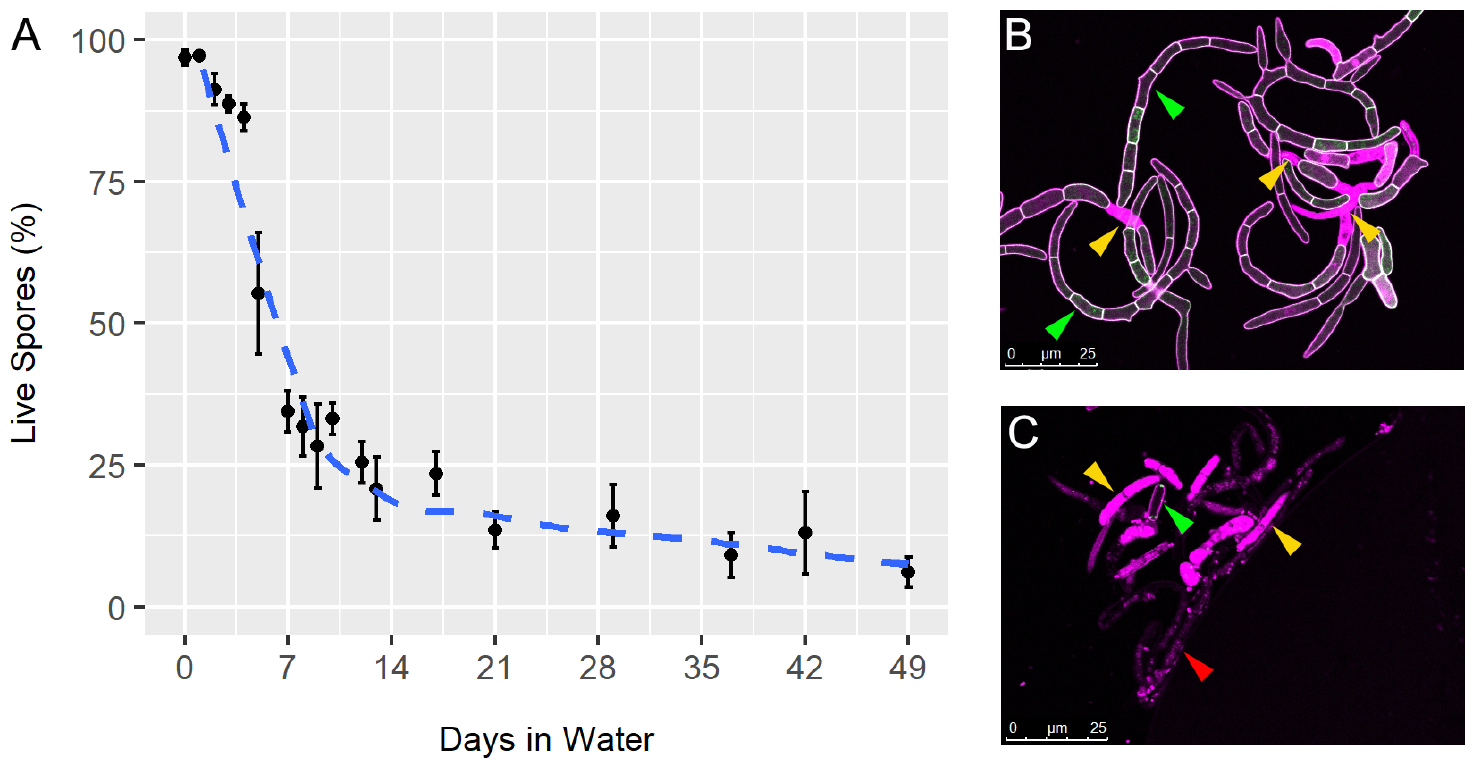
**A** – As Fig. 1, percentage viable blastospores (spores with at least one live cell; assessed by live/dead staining with propidium iodide(PI)) over time in water; B, C: examples of cells sampled and stained with PI after **B**: 5 days in water or **C**: 49 days in water. Yellow arrowheads indicate exemplar dead cells, flooded with PI stain. Green arrowheads indicate exemplar live cells, with staining at plasma membrane only. Red arrowheads indicate matter from dead, lysed cells.

**Figure S2.**
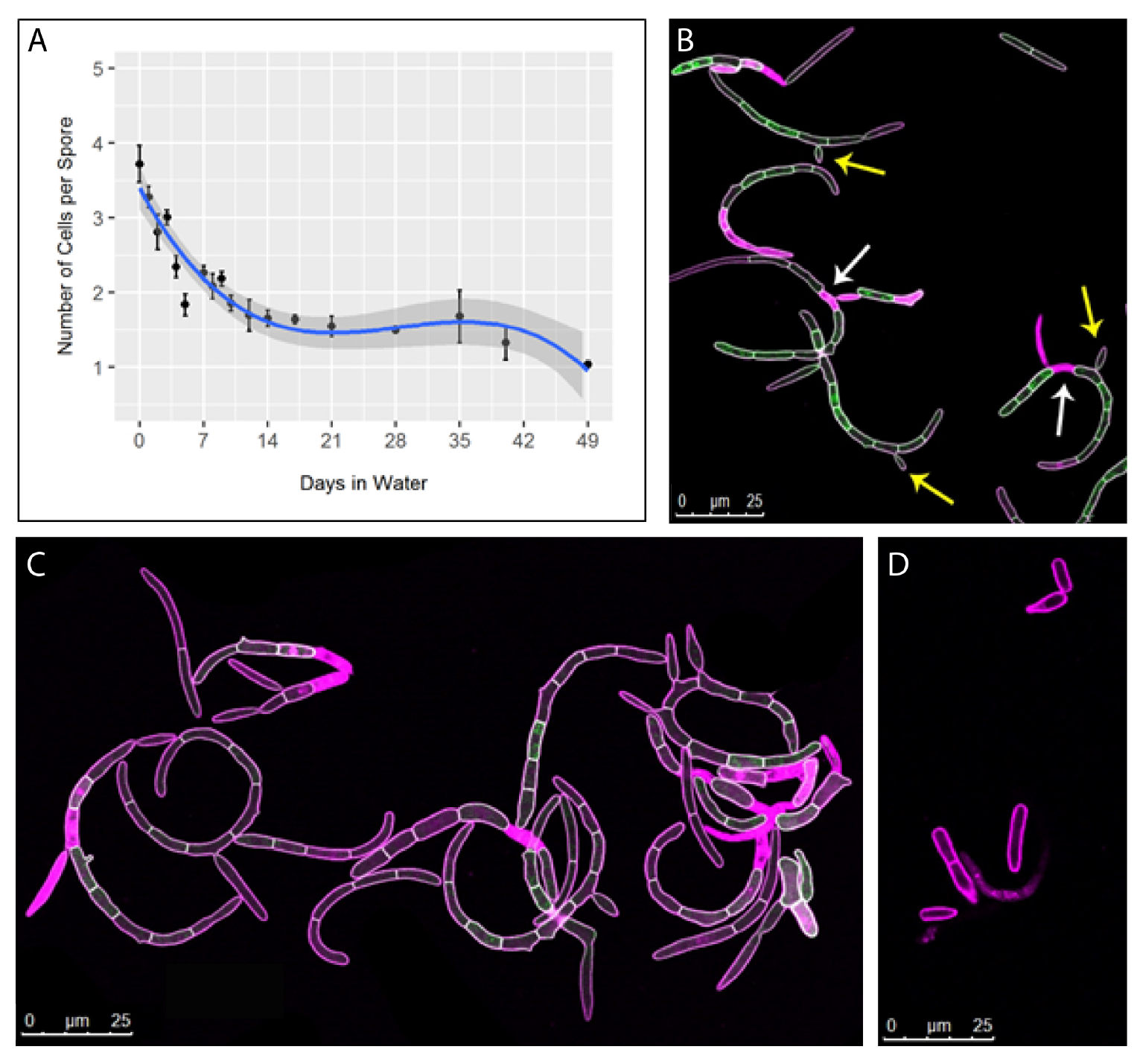
**A** – As Fig. 1D, average number of cells per individual blastospore over time in water; **B** – As Fig. 1C, example of cells sampled and stained with PI after 5 days in water **C**: exemplar blastospores stained with PI after 5 days in water; **D**: exemplar blastospores stained with PI after 42 days in water. Note the reduction in cells per individual blastospore between **B**/**C** and **D**. Arrows in **B** indicate possible reasons for this change: yellow arrows indicate bud formation, which would initially produce one-cell blastospores (buds); white arrows indicate cells in the middle of a blastospore (ie, non-terminal cells) which have died, potentially leading to splitting of the blastospore into two smaller individual blastospores.

**Figure S3.**
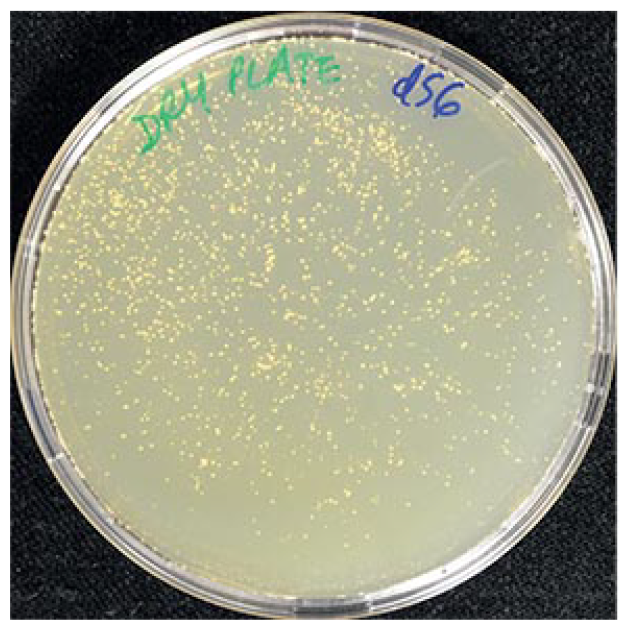
*Z. tritici* cells survive at least 56 days after aqueous suspensions are allowed to dry. An example plate is shown bearing colonies of *Z. tritici* which arose after cells were suspended in water, and then suspensions allowed to dry completely on a sterile petri dish surface before sealing the petri dish for 56 days. Dried cells were then resuspended in water and an aliquot pipetted onto YPD agar and incubated at 18 °C for 7 days. Colonies are clearly visible.

**Table S1:**
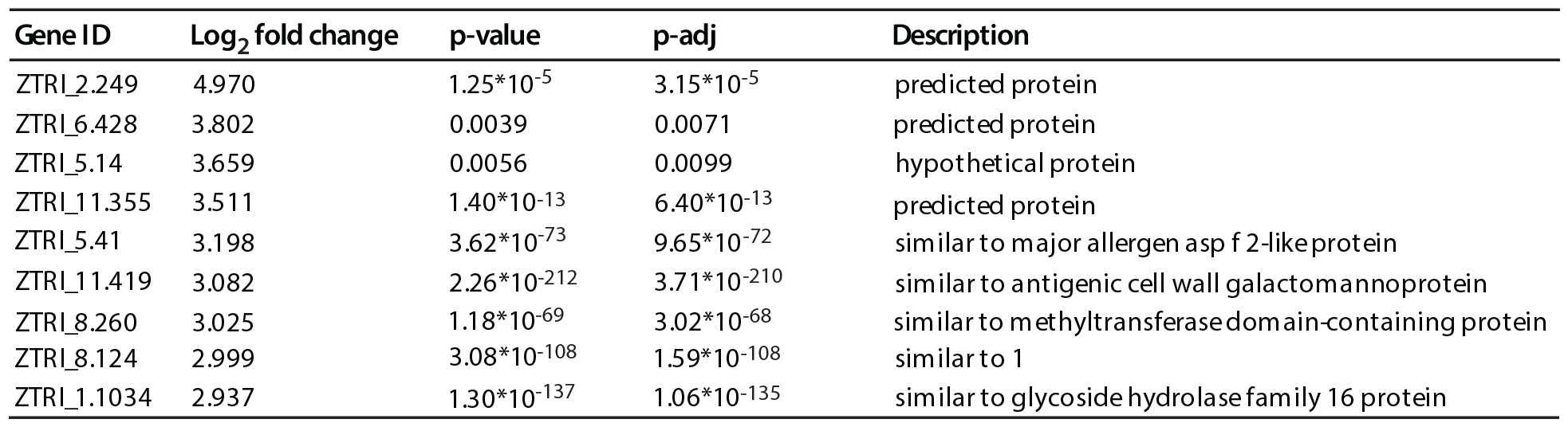
Ten genes most highly up-regulated among those uniquely up-regulated after 1 h in water.

**Table S2:**
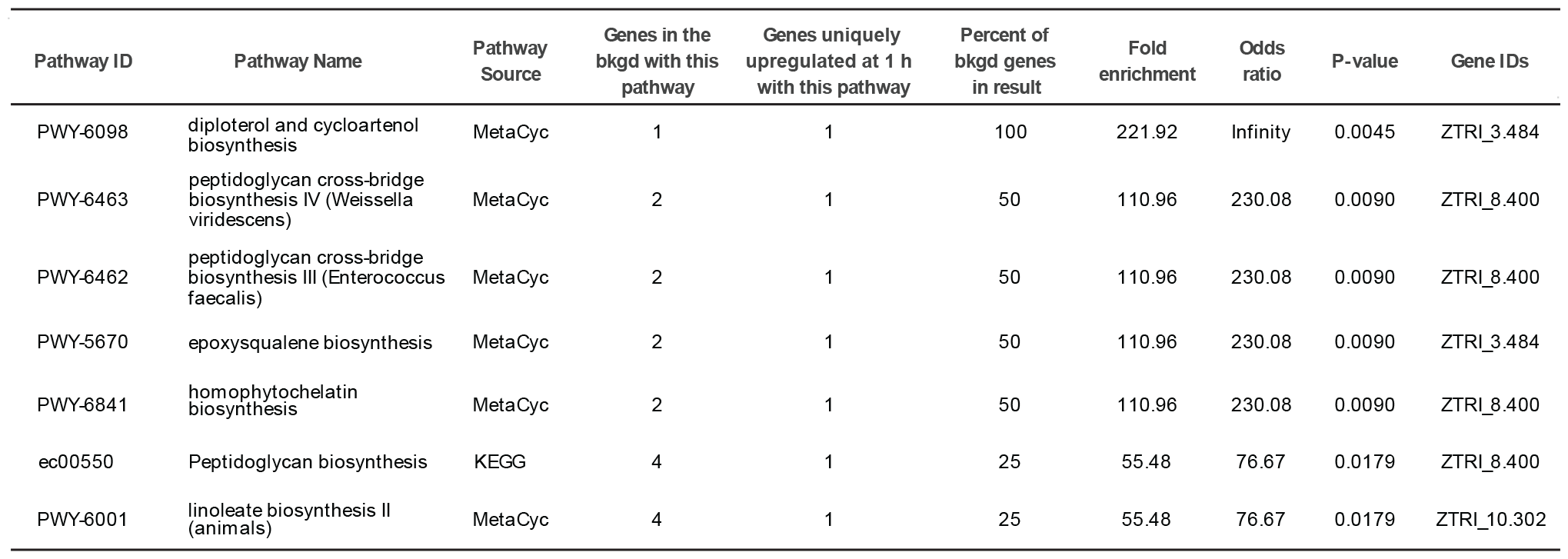
Metabolic pathway enrichment among genes uniquely up-regulated after 1 h in water. Enrichment analysis carried out using tools in FungiDB [1]. Threshold for inclusion in table P < 0.02

**Table S3:** Ten genes most highly up-regulated among all genes up-regulated after 1 h in water.

| Gene_ID | Log<sub>2</sub> fold change | p-value | p-adj | Description |
| --- | --- | --- | --- | --- |
| ZTRI_11.149 | 7.1903 | 4.09*10<sup>-11</sup> | 1.64*10<sup>-10</sup> | hypothetical protein |
| ZTRI_5.833 | 7.0802 | 4.97*10<sup>-11</sup> | 1.97*10<sup>-10</sup> | predicted protein |
| ZTRI_6.402 | 7.0332 | 0 | 0 | similar to heat shock protein 30 |
| ZTRI_11.277 | 6.9153 | 0 | 0 | similar to inorganic phosphate transporter pho84 |
| ZTRI_1.1590 | 6.8658 | 1.12*10<sup>-208</sup> | 1.77*10<sup>-206</sup> | predicted protein |
| ZTRI_11.422 | 6.7621 | 9.89*10<sup>-93</sup> | 3.79*10<sup>-91</sup> | hypothetical protein |
| ZTRI_8.748 | 6.7616 | 8.14*10<sup>-58</sup> | 1.59*10<sup>-56</sup> | similar to major facilitator superfamily transporter |
| ZTRI_1.498 | 6.7381 | 2.41*10<sup>-205</sup> | 3.69*10<sup>-201</sup> | similar to nadp-dependent alcohol dehydrogenase |
| ZTRI_1.494 | 6.5427 | 0 | 0 | hypothetical protein |
| ZTRI_8.112 | 6.4171 | 7.94*10<sup>-9</sup> | 2.70*10<sup>-8</sup> | predicted protein |

**Table S4:** Metabolic pathway enrichment among all genes up-regulated after 1 h in water. Enrichment analysis carried out using tools in FungiDB [1]. Threshold for inclusion in table P < 0.02

| Pathway ID | Pathway Name | Pathway Source | Genes in the bkgd with this pathway | Genes upregulated at 1h with this pathway | Percent of bkgd genes upregulated at 1 h | Fold enrichment | Odds ratio | P-value | Gene ID(s) |
| --- | --- | --- | --- | --- | --- | --- | --- | --- | --- |
| PWY-7288 | fatty acid β-oxidation (peroxisome, yeast) | MetaCyc | 9 | 5 | 55.6 | 11.85 | 25.9 | 0.0000 | ZTRI_10.223, 12.262, 2.730, 3.498, 5.646 |
| PWY-7337 | 10-*cis*-heptadecenoyl-CoA degradation (yeast) | MetaCyc | 11 | 5 | 45.5 | 9.7 | 17.26 | 0.0001 | ZTRI_10.223, 12.262, 3.264, 3.498, 5.646 |
| PWY-7338 | 10-*trans*-heptadecenoyl-CoA degradation (reductase-dependent, yeast) | MetaCyc | 11 | 5 | 45.5 | 9.7 | 17.26 | 0.0001 | ZTRI_10.223, 12.262, 3.264, 3.498, 5.646 |
| PWY66-391 | fatty acid β-oxidation VI (peroxisome) | MetaCyc | 13 | 5 | 38.5 | 8.21 | 12.94 | 0.0002 | ZTRI_10.223, 12.262, 2.730, 3.498, 5.646 |
| PWY-6001 | linoleate biosynthesis II (animals) | MetaCyc | 4 | 3 | 75 | 16 | 61.72 | 0.0004 | ZTRI_10.302, 2.371, 2.730 |
| PWY-7340 | 9*cis*, 11*trans*octadecadienoyl-CoA degradation (isomerase-dependent, yeast) | MetaCyc | 10 | 4 | 40 | 8.54 | 13.76 | 0.0008 | ZTRI_10.223, 12.262, 3.498, 5.646 |
| PWY-821 | superpathway of sulfur amino acid biosynthesis (*Saccharomyces cerevisiae*) | MetaCyc | 10 | 4 | 40 | 8.54 | 13.76 | 0.0008 | ZTRI_1.1103, 13.91, 3.850, 9.626 |
| PWY-7339 | 10-*trans*-heptadecenoyl-CoA degradation (MFE-dependent, yeast) | MetaCyc | 5 | 3 | 60 | 12.8 | 30.85 | 0.0009 | ZTRI_10.223, 3.498, 5.646 |
| PWY-5345 | superpathway of L-methionine biosynthesis (by sulfhydrylation) | MetaCyc | 18 | 5 | 27.8 | 5.93 | 7.96 | 0.0011 | ZTRI_1.1103, 13.91, 3.830, 3.850, 9.626 |
| PWY-5136 | fatty acid β-oxidation II (peroxisome) | MetaCyc | 11 | 4 | 36.4 | 7.76 | 11.79 | 0.0012 | ZTRI_10.223, 12.262, 2.730, 3.498 |
| SO4ASSIM-PWY | sulfate reduction I (assimilatory) | MetaCyc | 6 | 3 | 50 | 10.67 | 20.56 | 0.0018 | ZTRI_1.1103, 13.91, 9.626 |
| ec00071 | Fatty acid degradation | KEGG | 45 | 7 | 15.6 | 3.32 | 3.82 | 0.0045 | ZTRI_1.218, 1.391, 1.498, 10.223, 12.262, 2.730, 3.498 |
| TRESYN-PWY | trehalose biosynthesis I | MetaCyc | 3 | 2 | 66.7 | 14.23 | 40.98 | 0.0064 | ZTRI_6.64, 7.164 |
| PWY-7094 | fatty acid salvage | MetaCyc | 9 | 3 | 33.3 | 7.11 | 10.28 | 0.0069 | ZTRI_10.223, 2.730, 3.498 |
| PWY-561 | superpathway of glyoxylate cycle and fatty acid degradation | MetaCyc | 28 | 5 | 17.9 | 3.81 | 4.49 | 0.0088 | ZTRI_10.223, 12.262, 2.730, 3.498, 4.646 |
| PWY-6683 | sulfate reduction III (assimilatory) | MetaCyc | 4 | 2 | 50 | 10.67 | 20.49 | 0.0123 | ZTRI_1.1103, 9.626 |
| FAO-PWY | fatty acid β-oxidation I | MetaCyc | 11 | 3 | 27.3 | 5.82 | 7.7 | 0.0127 | ZTRI_10.223, 2.730, 3.498 |
| ec00590 | Arachidonic acid metabolism | KEGG | 31 | 5 | 16.1 | 3.44 | 3.97 | 0.0136 | ZTRI_1.542, 4.396, 7.434, 8.400, 9.547 |
| PWY-6920 | 6-geringol analog biosynthesis (engineered) | MetaCyc | 45 | 6 | 13.3 | 2.85 | 3.18 | 0.0175 | ZTRI_1.1720, 12.262, 13.96, 2.730, 5.646, 7.486 |
| PWY-6277 | superpathway of 5-aminoimidazole ribonucleotide biosynthesis | MetaCyc | 5 | 2 | 40 | 8.54 | 13.66 | 0.0199 | ZTRI_7.26, 8.17 |
| PWY-6121 | 5-aminoimidazole ribonucleotide biosynthesis I | MetaCyc | 5 | 2 | 40 | 8.54 | 13.66 | 0.0199 | ZTRI_7.26, 8.18 |

**Table S5:**
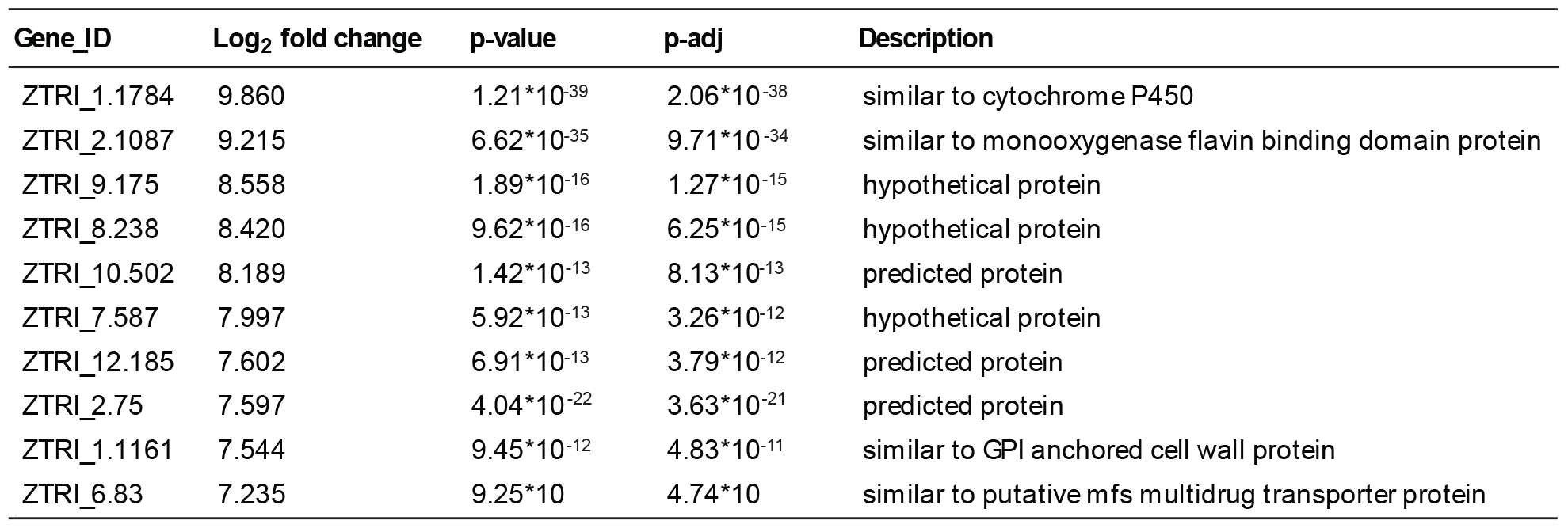
Ten genes most highly up-regulated among those uniquely up-regulated after 7 d in water.

| Gene_ID | Log<sub>2</sub> fold change | p-value | p-adj | Description |
| --- | --- | --- | --- | --- |
| ZTRI_1.1784 | 9.860 | $1.21 \times 10^{-39}$ | $2.06 \times 10^{-38}$ | similar to cytochrome P450 |
| ZTRI_2.1087 | 9.215 | $6.62 \times 10^{-35}$ | $9.71 \times 10^{-34}$ | similar to monooxygenase flavin binding domain protein |
| ZTRI_9.175 | 8.558 | $1.89 \times 10^{-16}$ | $1.27 \times 10^{-15}$ | hypothetical protein |
| ZTRI_8.238 | 8.420 | $9.62 \times 10^{-16}$ | $6.25 \times 10^{-15}$ | hypothetical protein |
| ZTRI_10.502 | 8.189 | $1.42 \times 10^{-13}$ | $8.13 \times 10^{-13}$ | predicted protein |
| ZTRI_7.587 | 7.997 | $5.92 \times 10^{-13}$ | $3.26 \times 10^{-12}$ | hypothetical protein |
| ZTRI_12.185 | 7.602 | $6.91 \times 10^{-13}$ | $3.79 \times 10^{-12}$ | predicted protein |
| ZTRI_2.75 | 7.597 | $4.04 \times 10^{-22}$ | $3.63 \times 10^{-21}$ | predicted protein |
| ZTRI_1.1161 | 7.544 | $9.45 \times 10^{-12}$ | $4.83 \times 10^{-11}$ | similar to GPI anchored cell wall protein |
| ZTRI_6.83 | 7.235 | $9.25 \times 10^{-10}$ | $4.74 \times 10^{-9}$ | similar to putative mfs multidrug transporter protein |

**Table S6:** Descriptions of PFAM domains associated with over-represented GO terms enrichment among genes uniquely up-regulated after 7 d in water. GO enrichment carried out using tools in fungiDB [1]. PFAM domains associated with the 52 genes located on core chromosomes and associated with significantly enriched GO terms (Bonferroni corrected P<0.05) were investigated. A list of PFAM domains (49 domains) was manually curated by inspection of the results of FungiDB’s InterPro Domain search for each gene product.

| PFAM Domain | Occurences | Description |
| --- | --- | --- |
| PF00067 | 15 | Cytochrome P450 |
| PF00264 | 4 | tyrosinase |
| PF00743 | 3 | Flavin-binding monooxygenase |
| PF00550 | 3 | Phosphopantetheine attachment site |
| PF07732 | 3 | Cu-oxidase_3 |
| PF01266 | 3 | DAO FAD dependent oxidoreductase |
| PF00724 | 2 | NADH:flavin oxidoreductase / NADH oxidase family |
| PF03098 | 2 | An-peroxidase |
| PF00501 | 2 | AMP-binding |
| PF07993 | 2 | Male sterility protein |
| PF04116 | 2 | Fatty acid hydroxylase superfamily - 2/52 |
| PF07731 | 2 | Cu-oxidase_2 |
| PF00975 | 1 | thioesterase |
| PF14765 | 1 | polyketide sythase dehydratase |
| PF07992 | 1 | Pyridine nucleotide-disulphide oxidoreductase - 1/52 |
| PF01565 | 1 | FAD-Binding_4 |
| PF08031 | 1 | BBE berberine and Berberine like |
| PF00173 | 1 | Cytochrome b5-like Heme/Steroid binding domain |
| PF00394 | 1 | Cu-oxidase multicopper oxidase |
| PF01794 | 1 | Ferric_reduct |
| PF08022 | 1 | fAD_binding_8 |
| PF08030 | 1 | NAD_binding_6 |
| PF02771 | 1 | Acyl-CoA dehydrogenase, N-terminal domain |
| PF02770 | 1 | Acyl-CoA dehydrogenase, middle domain |
| PF00441 | 1 | Acyl-CoA dehydrogenase, C-terminal domain |
| PF08534 | 1 | redoxin |
| PF04137 | 1 | ERO1 |
| PF00296 | 1 | Luciferase-like monooxygenase |
| PF00389 | 1 | D-isomer specific 2-hydroxyacid dehydrogenase, catalytic domain |
| PF02826 | 1 | D-isomer specific 2-hydroxyacid dehydrogenase, NAD binding domain |
| PF03358 | 1 | NADPH-dependent FMN reductase |
| PF13602 | 1 | Zinc-binding dehydrogenase |
| PF00390 | 1 | Malic enzyme, N-terminal domain |
| PF03949 | 1 | Malic enzyme, NAD binding domain |
| PF12766 | 1 | Pyridoxamine 5'-phosphate oxidase |
| PF08240 | 1 | Alcohol dehydrogenase GroES-like domain |
| PF00107 | 1 | Zinc-binding dehydrogenase |
| PF01494 | 1 | FAD binding domain |
| PF16073 | 1 | Starter unit:ACP transacylase in aflatoxin biosynthesis |
| PF00109 | 1 | Beta-ketoacyl synthase, N-terminal domain |
| PF02801 | 1 | Beta-ketoacyl synthase, C-terminal domain |
| PF00698 | 1 | Acyl transferase domain |
| PF14765 | 1 | Polyketide synthase dehydratase |
| PF00975 | 1 | Thioesterase domain |
| PF00970 | 1 | Oxidoreductase FAD-binding domain |
| PF00175 | 1 | Oxidoreductase NAD-binding domain |
| PF00732 | 1 | GMC oxidred N |
| PF05199 | 1 | GMC oxidred C |

**Table S7:**
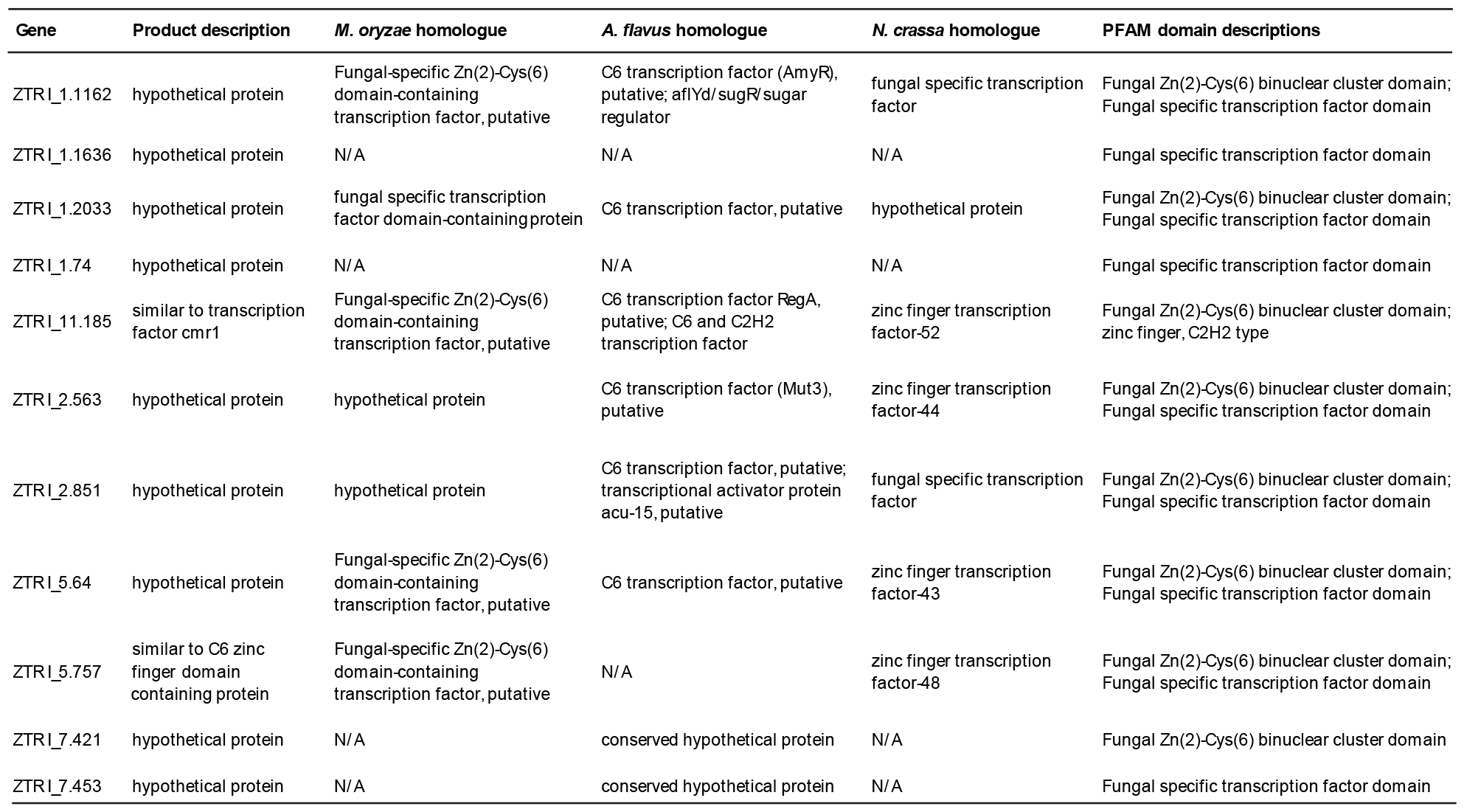
transcription factors among the genes uniquely up-regulated after 7 days in water. The list of 437 genes uniquely up-regulated after 7 days in water was searched using the search strategies function in FungiDB for the string ‘transcr*’. This returned eleven genes, shown.

**Table S8:** Ten genes most highly down-regulated among those uniquely down-regulated after 1 h in water.

| Gene_ID | Log<sub>2</sub> fold change | p-value | p-adj | Description |
| --- | --- | --- | --- | --- |
| ZTR1_13.286 | -4.4646 | 0.0002 | 0.0004 | hypothetical protein |
| ZTR1_15.3 | -4.3526 | 0.0001 | 0.0003 | predicted protein |
| ZTR1_4.50 | -4.2066 | 0.0007 | 0.0014 | predicted protein |
| ZTR1_4.268 | -4.0093 | 0.0007 | 0.0015 | hypothetical protein |
| ZTR1_6.20 | -4.0026 | 0.0017 | 0.0032 | predicted protein |
| ZTR1_1.140 | -3.6359 | 2.54*10<sup>-10</sup> | 9.66*10<sup>-10</sup> | hypothetical protein |
| ZTR1_14.64 | -3.6275 | 0.0089 | 0.0153 | hypothetical protein |
| ZTR1_7.24 | -3.6096 | 0.0001 | 0.0002 | predicted protein |
| ZTR1_20.77 | -3.5823 | 0.0001 | 0.0003 | hypothetical protein |
| ZTR1_3.70 | -3.5751 | 0.0001 | 0.0003 | hypothetical protein |

**Table S9:** transcription factors among the genes uniquely down-regulated after 1 h in water. The list of 207 genes uniquely down-regulated after 1 h in water was searched using the search strategies function in FungiDB for the string ‘transcr*’. This returned twenty genes, shown.

| Gene | Product description | *M. oryzae* homologue | *A. flavus* homologue | *N. crassa* homologue | PFAM domain descriptions |
| --- | --- | --- | --- | --- | --- |
| ZTRI_1.2104 | predicted protein | n/a | n/a | n/a | n/a |
| ZTRI_1.456 | hypothetical protein | ZZ type zinc finger domain-containing protein | ZZ type zinc finger domain-containing protein | zinc finger transcription factor-55 | central domain |
| ZTRI_2.1024 | predicted protein | n/a | n/a | n/a | n/a |
| ZTRI_2.860 | similar to DNA binding regulatory protein AmdX | DNA binding regulatory protein AmdX | C2H2 transcription factor (AmdX), putative | zinc finger transcription factor-31 | Zinc finger C2H2-type; Transcription factor domain |
| ZTRI_3.1064 | similar to ankyrin repeat-containing protein | ankyrin repeat domain-containing protein 28, 29; ankyrin repeat and SOCS box protein 7 | ankyrin repeat and socs box protein, putative | pfs domain-containing protein | Ankyrin repeat-containing domain |
| ZTRI_4.735 | hypothetical protein | n/a | n/a | n/a | Zn(2)-C6 fungal-type DNA-binding domain |
| ZTRI_5.105 | hypothetical protein | Fungal-specific Zn(2)-Cys(6) domain-containing transcription factor, putative | C6 finger domain protein, putative | hypothetical protein | Zn(2)-C6 fungal-type DNA-binding domain |
| ZTRI_5.209 | hypothetical protein | hypothetical protein | n/a | hypothetical protein | n/a |
| ZTRI_5.26 | similar to transcription factor MADS-box | MADS box protein | MADS box transcription factor Mcm 1 | MADS box protein | Transcription factor, MADS-box |
| ZTRI_5.48 | predicted protein | hypothetical protein | F-box and wd40 domain protein, putative; vegetative incompatibility WD repeat protein, putative; G-protein beta WD-40 repeats containing protein, putative | n/a | n/a |
| ZTRI_5.54 | predicted protein | tropomyosin-1 alpha chain | spindle-pole body protein (Pcp1), putative; M protein repeat protein | Septal pore-associated protein, putative | n/a |
| ZTRI_5.787 | similar to C2H2 finger domain protein FlbC | C2H2 finger domain-containing protein FlbC | C2H2 conidiation transcription factor FlbC | C2H2 finger domain-containing protein FlbC | Zinc finger C2H2-type |
| ZTRI_6.248 | hypothetical protein | Fungal-specific Zn(2)-Cys(6) domain-containing transcription factor, putative; ankyrin repeat domain -containing protein 28 | ankyrin repeat domain-containing protein 28; 5-AMP-activated protein kinase, putative; vps9-ankyrin repeat-containing protein, putative; Pfs, NACHT and Ankyrin domain protein; DNA-binding protein rfxank, putative; nf kappa B inhibitor I-kappa-B, putative; m-phase phosphoprotein, putative; serine/threonine-protein kinase ripk4, putative | pfs domain-containing protein | Ankyrin repeat-containing domain |
| ZTRI_7.674 | hypothetical protein | LysM domain-containing protein | muramidase, putative | hypothetical protein | LysM domain |
| ZTRI_8.213 | hypothetical protein | potassium/sodium efflux P-typeATPase | sodium P-type ATPase, putative | calcium-transporting ATPase 3; Na or K P-type ATPase | Cation-transporting P-type ATPase, N-terminal; Cation-transporting P-type ATPase, C-terminal; Fungal transcription factor; Amidase signature domain |
| ZTRI_8.661 | similar to monosaccharide transporter | high-affinity glucose transporters ght2; RGT2; glucose transporter rco3; quinate permease | MFS monosaccharide transporter, putative; MFS glucose transporter, putative; MFS quinate transporter, putative | MFS monosaccharide transporter; sugar transporter-22, 5, 9, 7, 23, 8, 6, 10; RCO3; quinate permease | Major facilitator, sugar transporter-like |
| ZTRI_10.332 | hypothetical protein | medusa | transcriptional regulator Medusa | Acr1 | n/a |
| ZTRI_10.451 | similar to putative c6 zinc finger domain protein | n/a | n/a | n/a | Zn(2)-C6 fungal-type DNA-binding domain |
| ZTRI_10.89 | similar to sodium/phosphate symporter | phosphate-repressible phosphate permease | phosphate-repressible Na+/phosphate cotransporter Pho89, putative | phosphate-repressible phosphate permease | n/a |
| ZTRI_12.279 | hypothetical protein | maltose O-acetyltransferase; nodulation protein L | acetyltransferase, CysE/LacA/LpxA/NodL family | female fertility-7; carbohydrate O-acetyltransferase | Zn(2)-C6 fungal-type DNA-binding domain; hexapeptide repeat |

**Table S10:** Metabolic pathway enrichment among genes uniquely down-regulated after 1 h in water. Enrichment analysis carried out using tools in FungiDB [1]. Threshold for inclusion in table P < 0.02

| Pathway ID | Pathway Name | Pathway Source | Genes in the bkgd with this pathway | Genes down-regulated at 1 h with this pathway | Percent of bkgd genes down-regulated at 1h | Fold enrichment | Odds ratio | P-value | Gene IDs |
| --- | --- | --- | --- | --- | --- | --- | --- | --- | --- |
| METHYLGALLATE-DEGRADATION | methylgallate degradation | MetaCyc | 13 | 3 | 23.1 | 16.28 | 22.15 | 0.0007 | ZTRI_4.700, 6.150, 8.548 |
| P184-PWY | protocatechuate degradation I ( meta-cleavage pathway) | MetaCyc | 6 | 2 | 33.3 | 23.51 | 36.2 | 0.0029 | ZTRI_4.700, 8.548 |
| PWY-6338 | superpathway of vanillin and vanillate degradation | MetaCyc | 7 | 2 | 28.6 | 20.15 | 28.95 | 0.0040 | ZTRI_4.700, 8.548 |
| GALLATE-DEGRADATION-I | gallate degradation II | MetaCyc | 11 | 2 | 18.2 | 12.82 | 16.07 | 0.0100 | ZTRI_4.700, 8.548 |
| ec00600 | metabolism | KEGG | 97 | 5 | 5.2 | 3.64 | 4.09 | 0.0113 | ZTRI_1.1605, 1.726, 3.1013, 10.375, 13.352 |
| PWY-7111 | pyruvate fermentation to isobutanol (engineered) | MetaCyc | 69 | 4 | 5.8 | 4.09 | 4.57 | 0.0157 | ZTRI_2.96, 7.431, 10.375, 10.494 |
| PWY-6339 | syringate degradation | MetaCyc | 70 | 4 | 5.7 | 4.03 | 4.5 | 0.0165 | ZTRI_4.700, 6.150, 8.548, 10.494 |
| PWY-7824 | prunasin and amygdalin biosynthesis | MetaCyc | 40 | 3 | 7.5 | 5.29 | 5.94 | 0.0183 | ZTRI_1.909, 3.1013, 10.352 |
| PWY-5707 | propanethial S-oxide biosynthesis | MetaCyc | 15 | 2 | 13.3 | 9.4 | 11.11 | 0.0184 | ZTRI_8.25, 10.265 |

**Table S11:** Ten genes most highly down-regulated after 1 h in water.

| Gene_ID | Log<sub>2</sub> fold change | p-value | p-adj | Description |
| --- | --- | --- | --- | --- |
| ZTRI_13.56 | -7.199 | 8.43*10<sup>-23</sup> | 5.97*10<sup>-22</sup> | similar to phthalate transporter |
| ZTRI_5.479 | -5.932 | 1.48*10<sup>-68</sup> | 3.68*10<sup>-67</sup> | similar to GPR1/FUN34/YaaH-class plasma membrane protein |
| ZTRI_3.488 | -5.604 | 3.87*10<sup>-07</sup> | 1.14*10<sup>-06</sup> | similar to 15-hydroxyprostaglandin dehydrogenase |
| ZTRI_9.43 | -5.545 | 7.64*10<sup>-07</sup> | 2.19*10<sup>-06</sup> | predicted protein |
| ZTRI_5.399 | -5.431 | 5.34*10<sup>-07</sup> | 1.55*10<sup>-06</sup> | hypothetical protein; homology to *Aspergillus sp.* acetolactate synthase |
| ZTRI_1.1987 | -5.378 | 0 | 0 | similar to homogentisate 1 |
| ZTRI_1.1989 | -5.366 | 8.43*10<sup>-263</sup> | 2.19*10<sup>-260</sup> | similar to cytochrome P450 phenylacetate 2-hydroxylase |
| ZTRI_3.747 | -5.321 | 1.15*10<sup>-102</sup> | 5.46*10<sup>-101</sup> | similar to major facilitator superfamily transporter |
| ZTRI_1.671 | -5.244 | 7.13*10<sup>-48</sup> | 1.14*10<sup>-46</sup> | similar to epoxide hydrolase |
| ZTRI_6.593 | -5.120 | 2.05*10<sup>-06</sup> | 5.60*10<sup>-06</sup> | hypothetical protein |

**Table S12:**
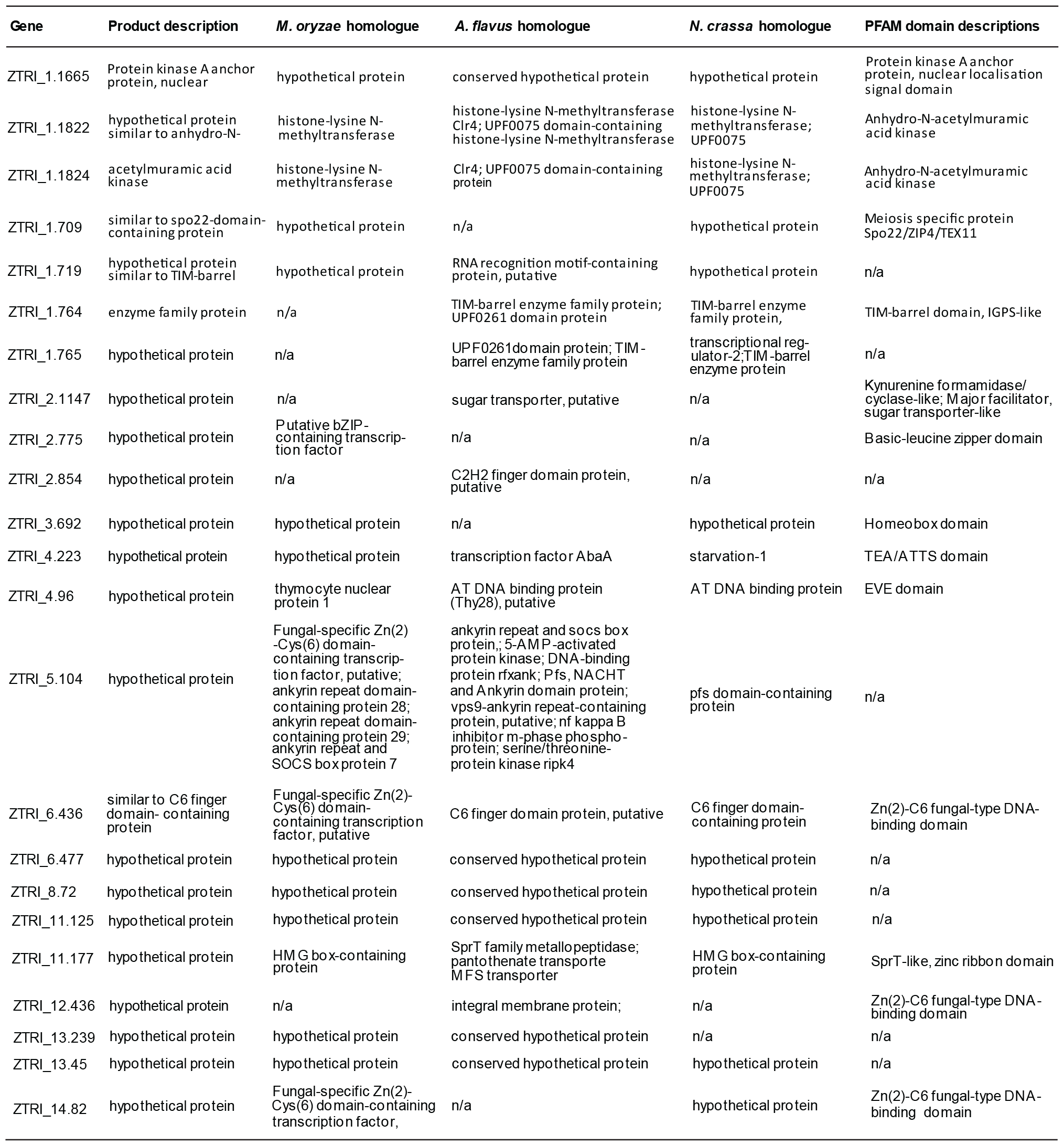
transcription factors among the all genes down-regulated after 1 h in water. The list of 492 genes down-regulated after 1 h in water was searched using the search strategies function in FungiDB for the string ‘transcr*’. This returned twenty-three genes, shown.

**Table S13:**
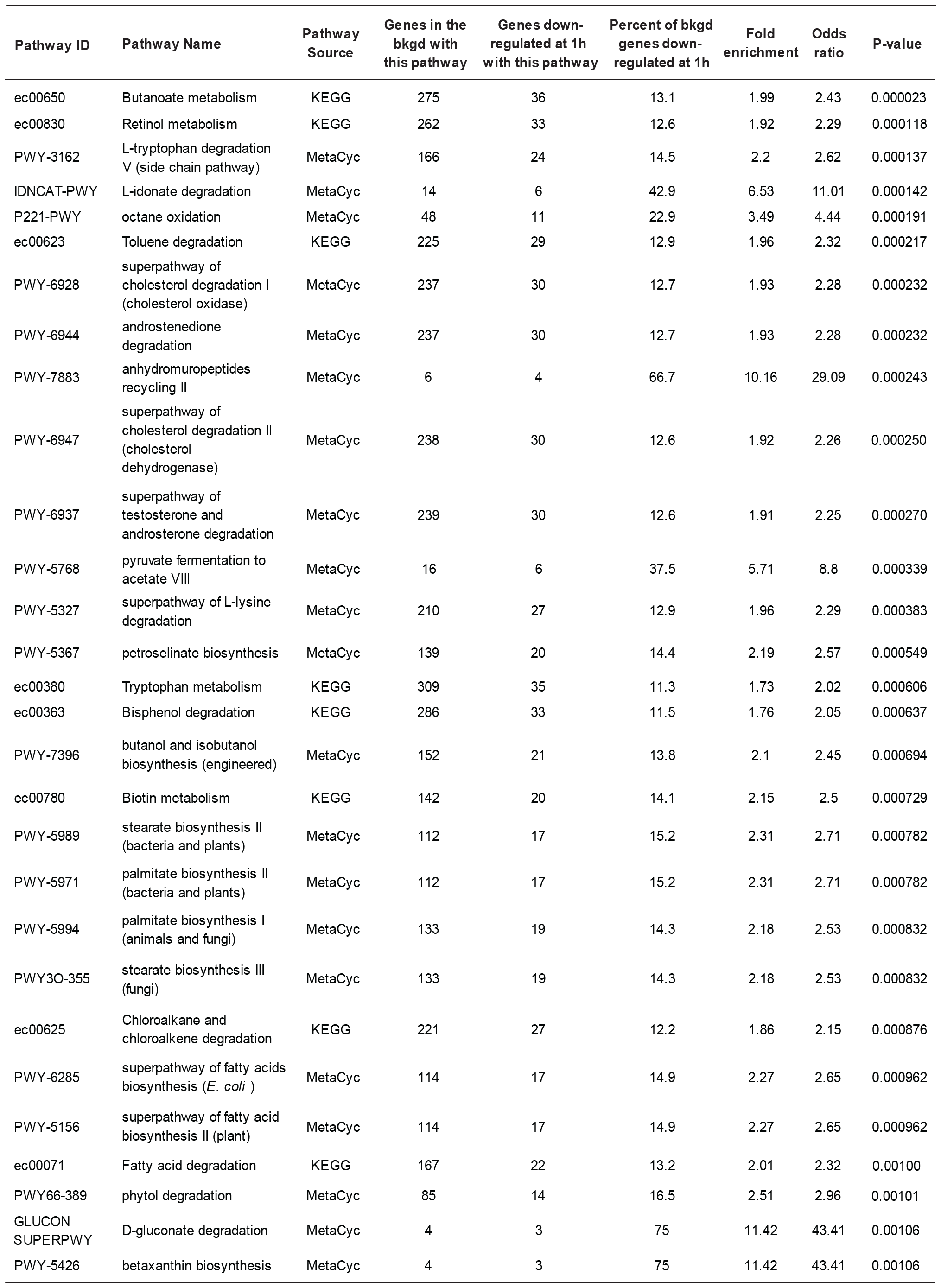
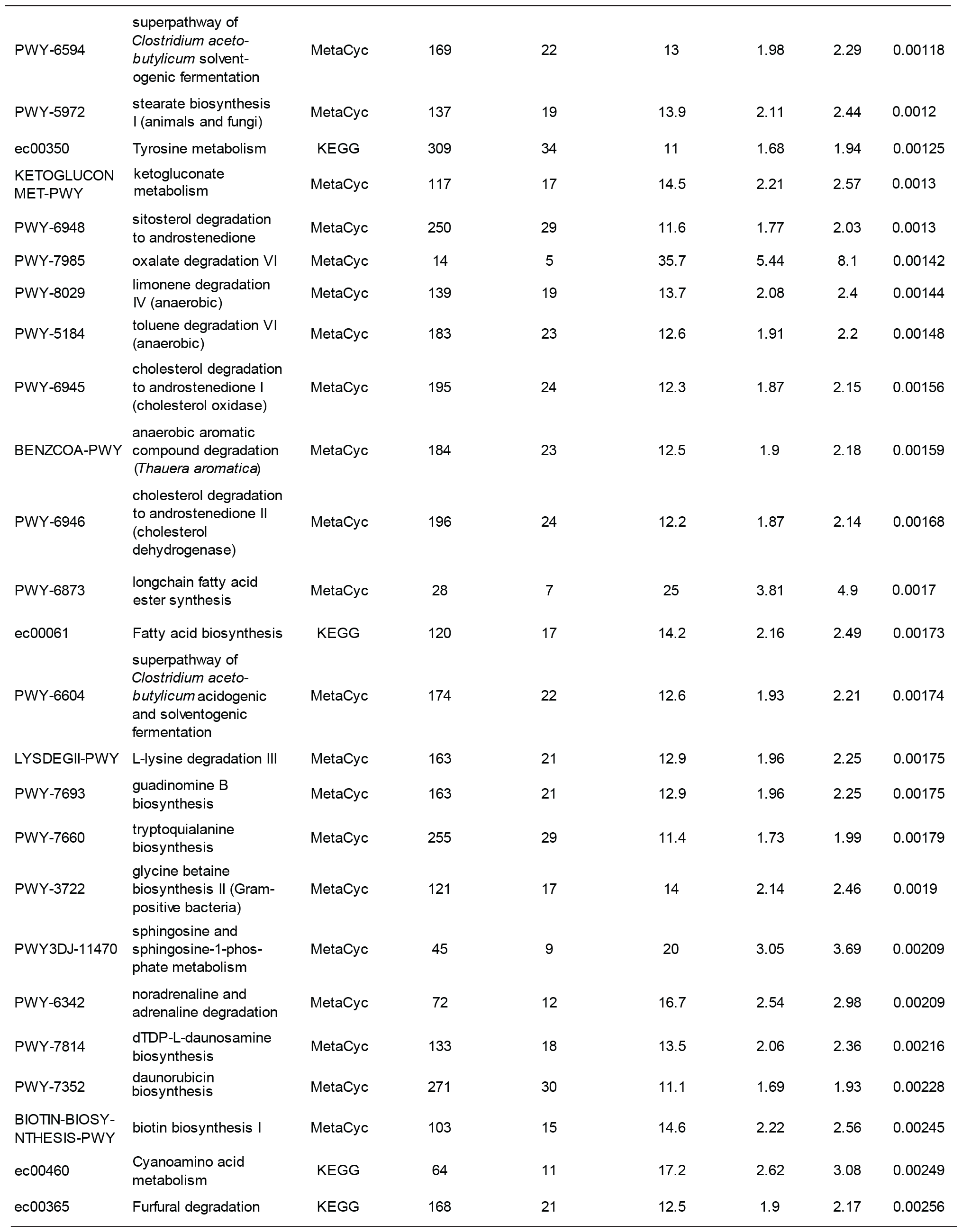
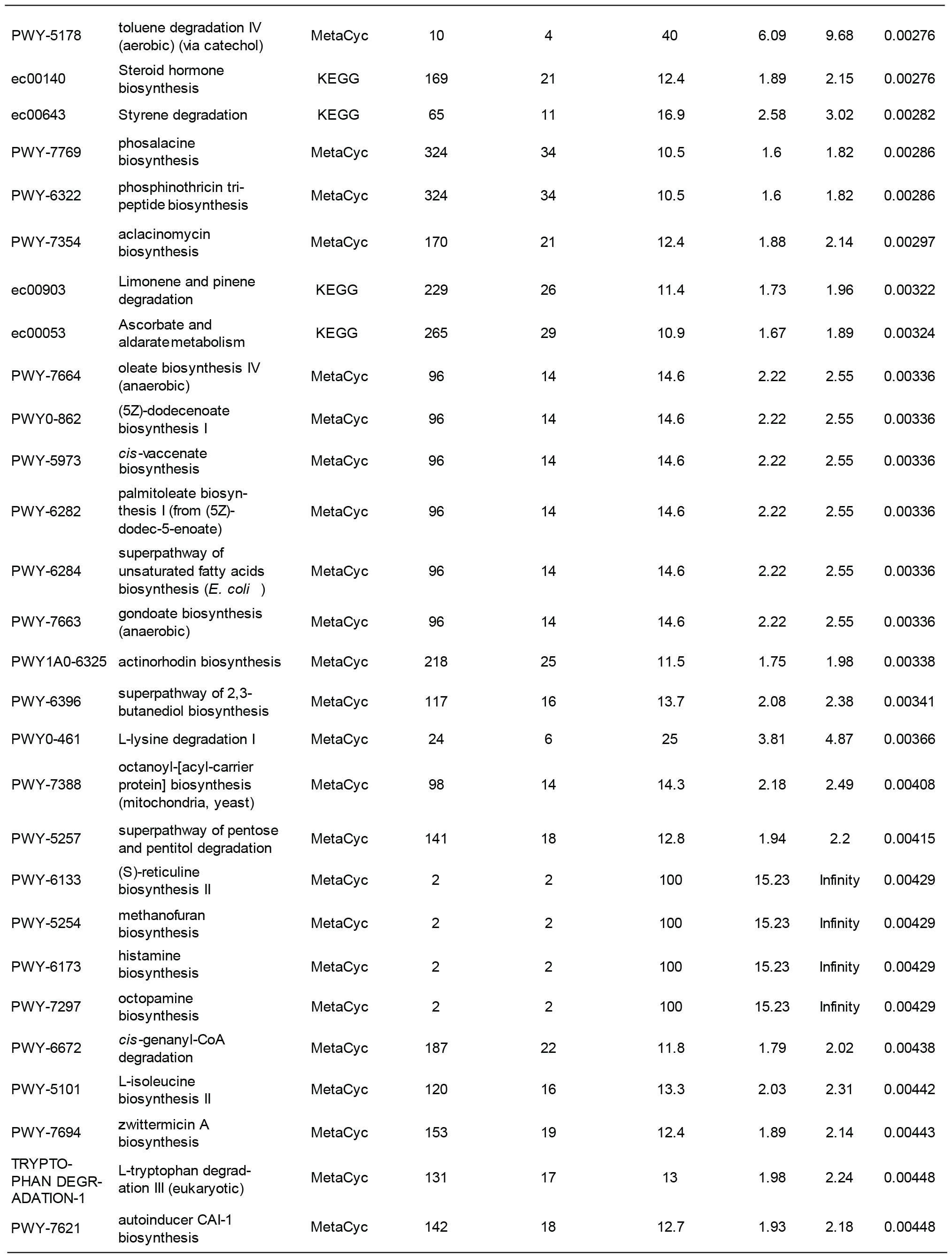
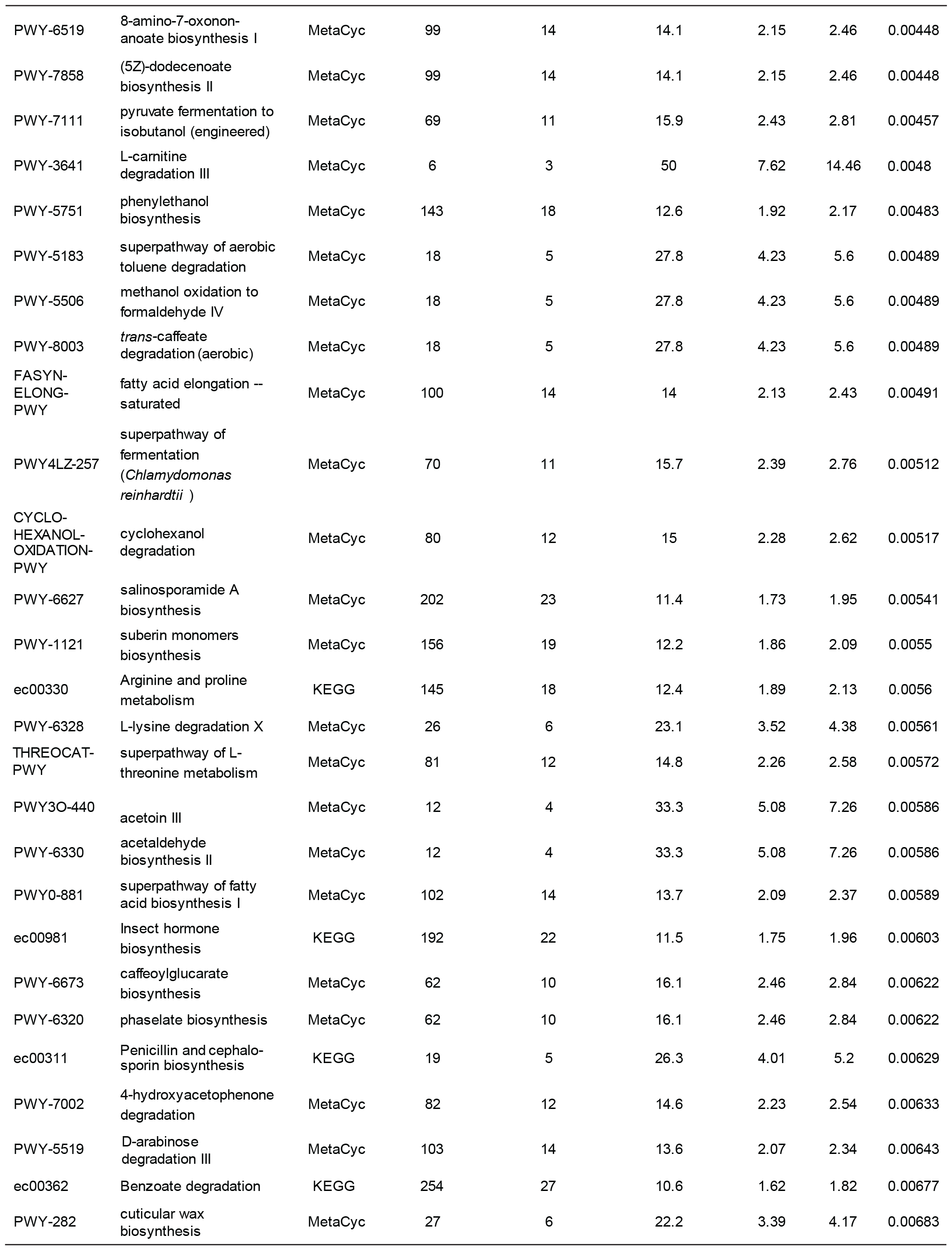
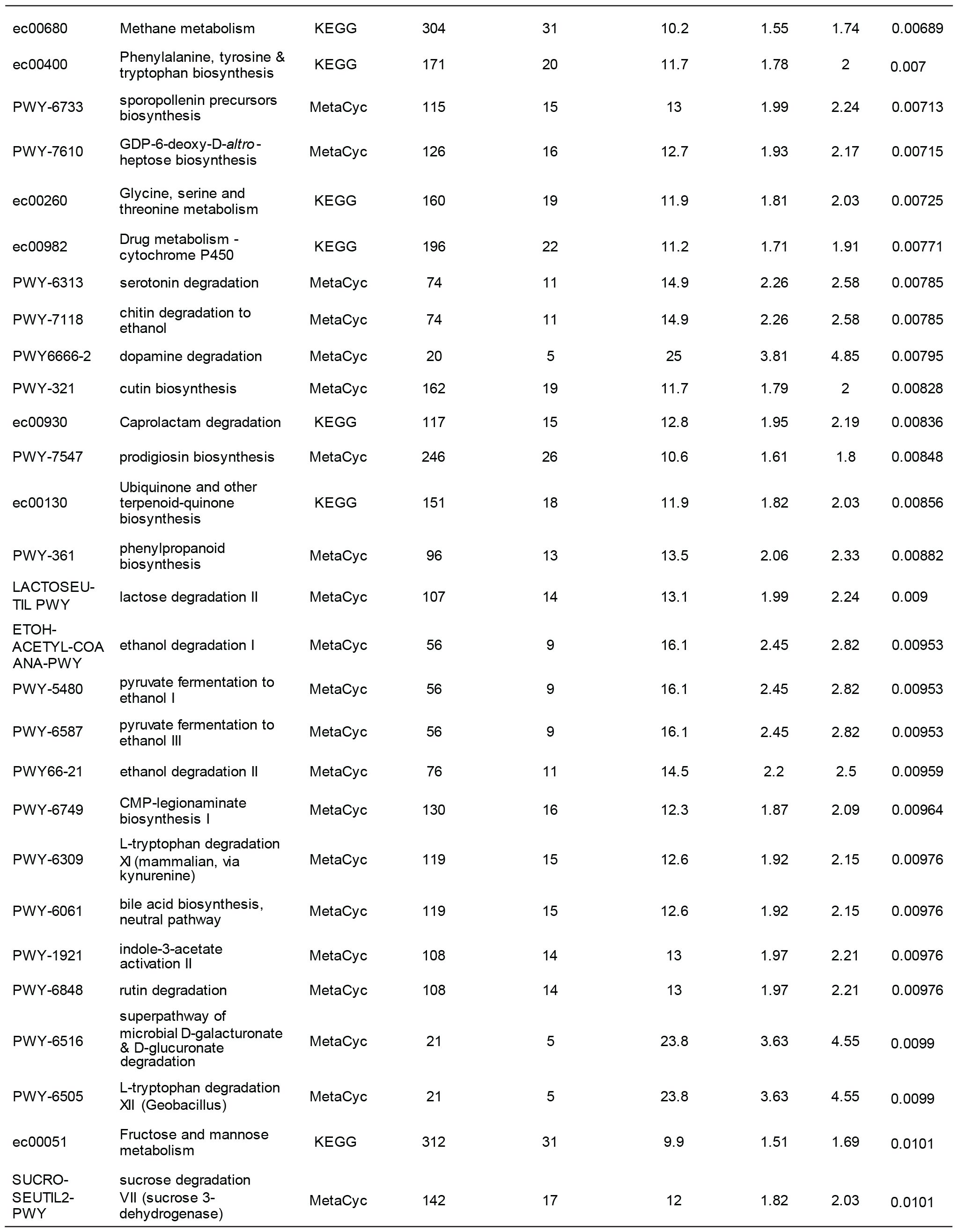
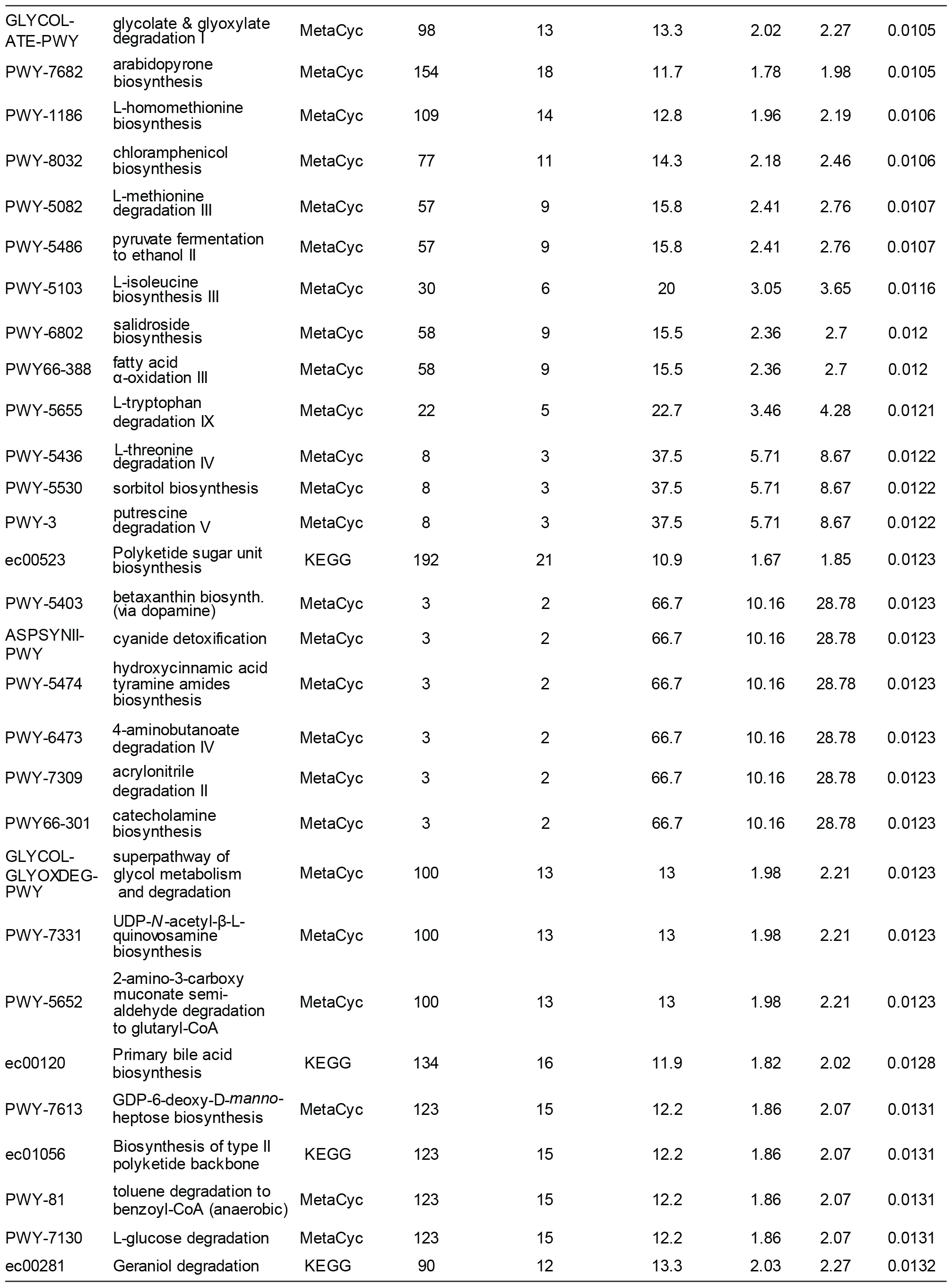
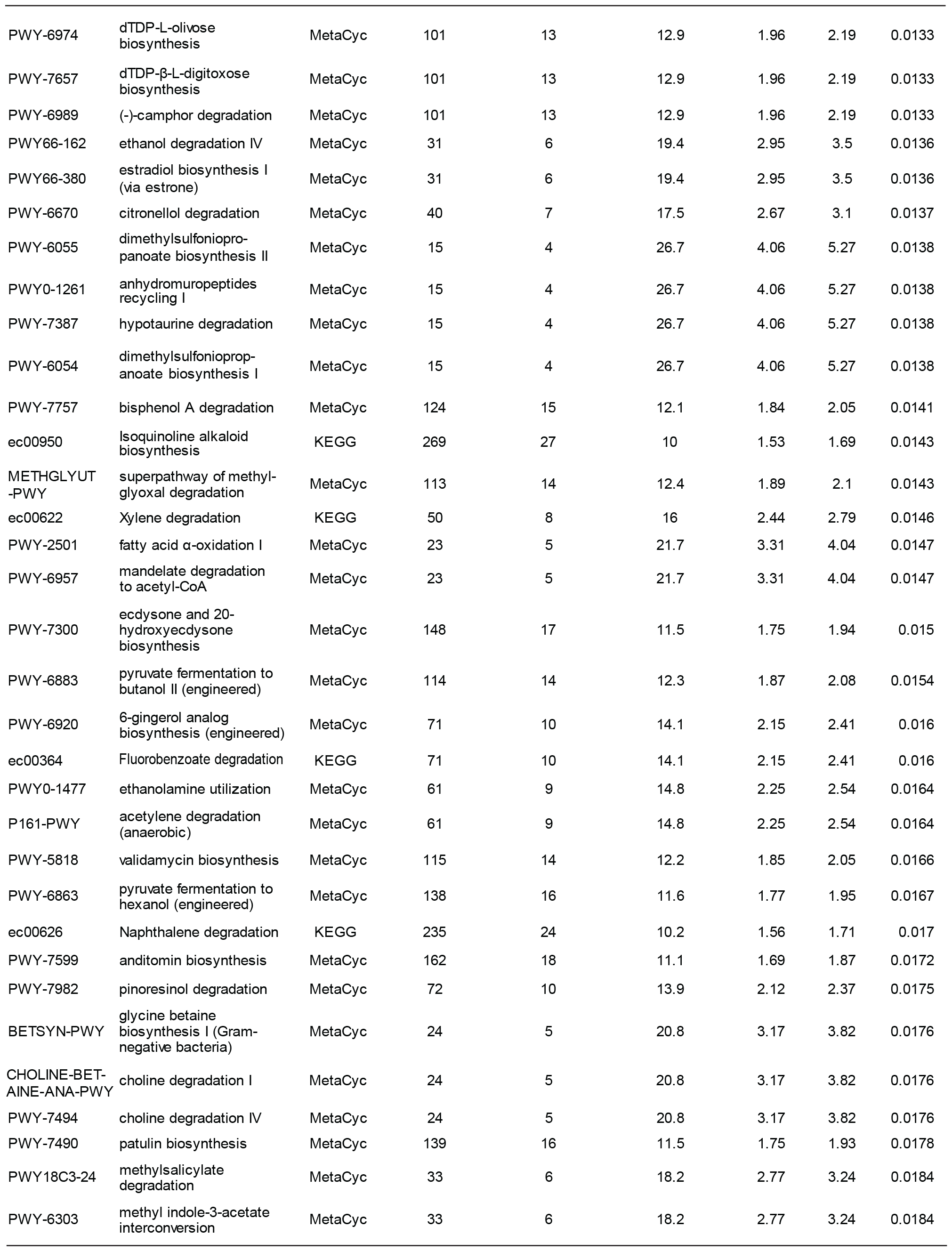
Metabolic pathway enrichment among genes uniquely down-regulated after 1 h in water. Enrichment analysis carried out using tools in FungiDB [1]. Threshold for inclusion in table P < 0.02

**Table S14:**
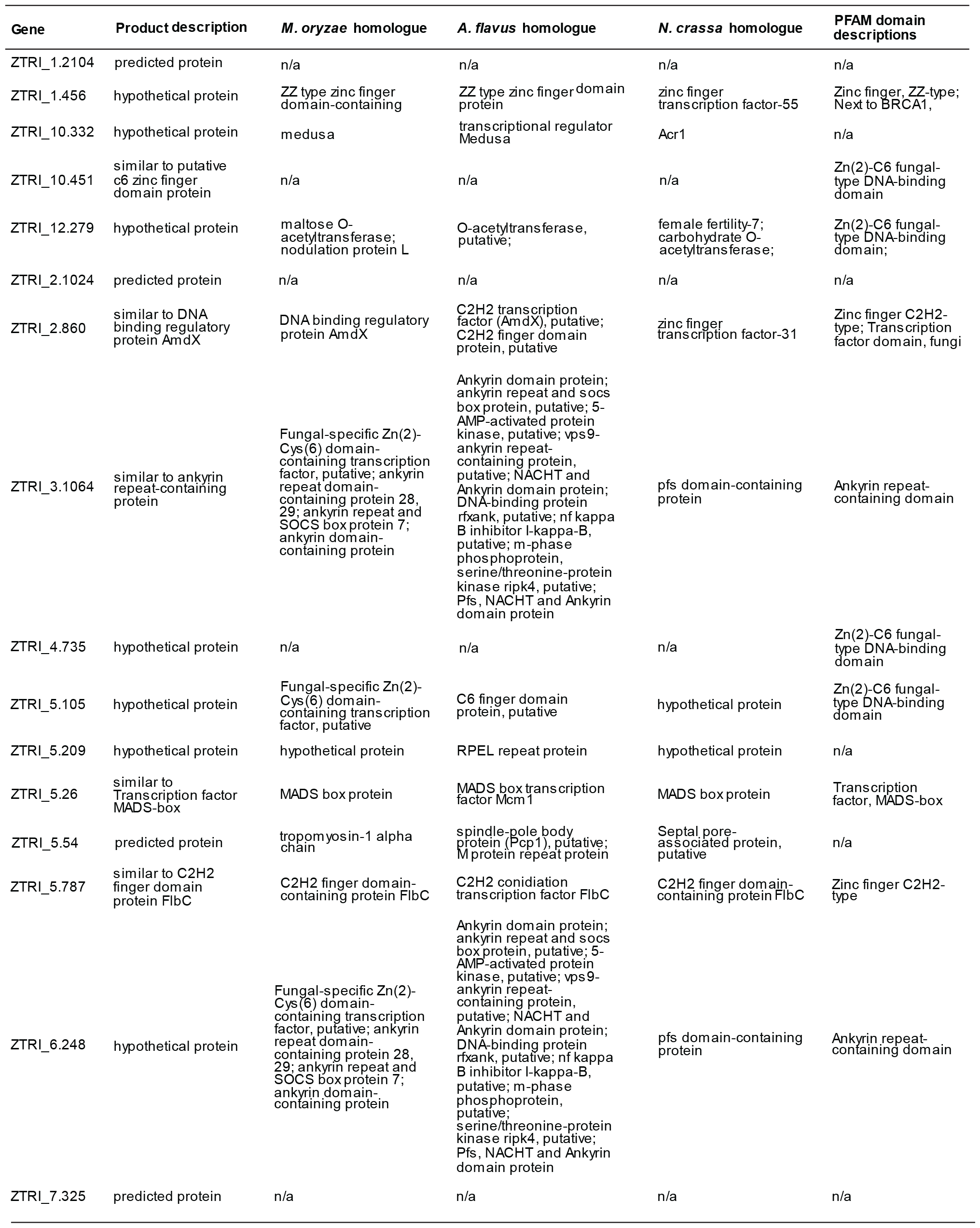
transcription factors among the all genes down-regulated after 1 h in water. The list of genes down-regulated after 7 days in water was searched using the search strategies function in FungiDB for the string ‘transcr*’. This returned twenty-three genes, shown.

**Table S15:**
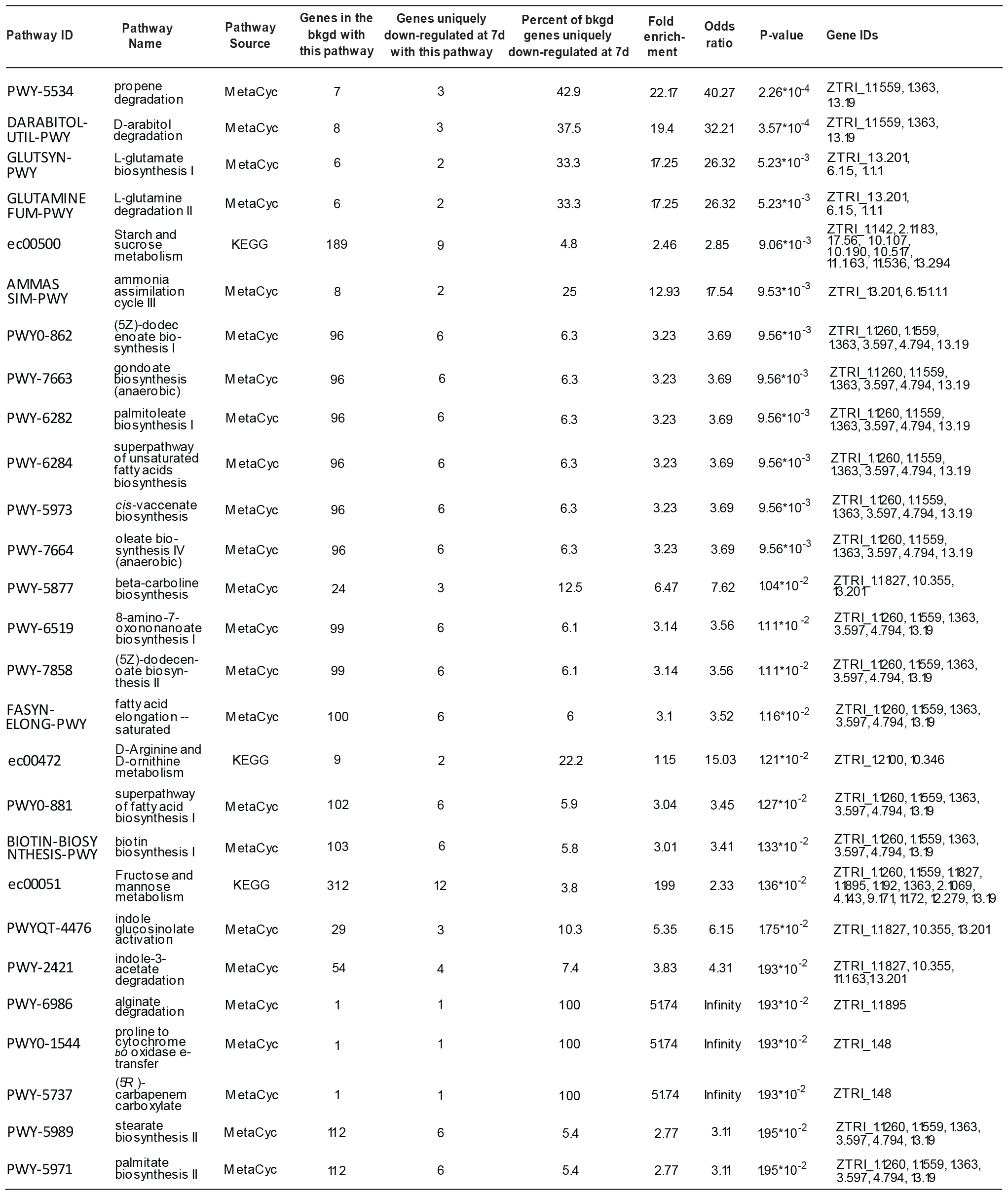
Metabolic pathway enrichment among genes uniquely down-regulated after 7 days in water. Enrichment analysis carried out using tools in FungiDB [1]. Threshold for inclusion in table P < 0.02

